# SVC: A Vision Transformer-based Spatially and Virtually Embedded Cell Model for Deciphering Subcellular Spatial Transcriptomic Heterogeneity

**DOI:** 10.64898/2026.10.07.757371

**Authors:** Hui Wan, Penghui Yang, Siyu Hou, Jade Wang, Xiang Zhou

## Abstract

Subcellular transcript localization shapes gene regulation and cellular function, yet profiling it at scale remains difficult, limiting our ability to study subcellular heterogeneity across tissues and disease. Here, we present SVC, a Vision Transformer-based spatially and virtually embedded cell model that learns subcellular transcript organization within native tissue context directly from subcellular-resolution spatial transcriptomics. SVC represents each gene’s subcellular localization as an image patch and uses other genes within the same cell as context, enabling subcellular contextual learning of spatial dependencies that govern intracellular organization. By integrating gene-function priors, cell morphology, spatial context, and, when available, cell identity, SVC captures unified, spatially grounded representations across genes and cells that connect subcellular organization to cellular state and tissue microenvironment. Across diverse subcellular-resolution spatial transcriptomics datasets, SVC accurately predicts cell-specific subcellular transcript localization and co-organization, generalizes across tissues and technical platforms, and supports subcellular imputation of unmeasured genes and *in silico* perturbation. Leveraging these capabilities, SVC highlights disease-associated patterns of intracellular transcript reorganization and spatially heterogeneous responses to gene perturbations. Beyond prediction, SVC reveals systematic variation in subcellular transcript organization associated with gene function, cellular states, and tissue microenvironments, uncovering *de novo* spatial biology that is not accessible by cell-level analysis. SVC’s learned representations further transfer to cell- and tissue-level tasks, including cell-level expression imputation, cell clustering, and spatial domain detection. Together, SVC provides a general framework for modeling and interrogating subcellular transcript organization *in situ*, enabling scalable discovery of subcellular spatial biology.

## Introduction

Spatial transcriptomics (ST) has undergone rapid development in recent years, evolving from early spot-level technologies, such as Spatial Transcriptomics^1^ and 10x Visium, to single-cell resolution platforms, such as Slide-seq^2^ and Stereo-seq^3^, and more recently to subcellular-resolution methods, such as MERFISH^4^, seqFISH+^5^, and Xenium^6, 7^. These recent subcellular-resolution ST technologies allow the measurement of transcript localization within cells, providing an unprecedented opportunity to characterize spatial gene expression patterns at the intracellular level. This subcellular capability, in turn, helps uncover the biological mechanisms underlying spatial and transcriptional heterogeneity within cells.

Understanding subcellular transcript localization is crucial, as different transcripts exhibit distinct spatial localization patterns within cells, and their proper subcellular positioning is essential for maintaining cellular function and behavior. For example, β– actin mRNA localizes to the leading edge of migrating cells, establishing polarity and promoting persistent directional movement^8^. Similarly, *CaMKIIα* mRNA localizes to dendrites and dendritic spines in mammalian neurons, enabling local translation and supporting spine-specific regulation of synaptic strength^9^. More broadly, variations in subcellular transcript localization across cells can serve as sensitive indicators of cellular state, reflecting polarity, activation status, or early stages of differentiation. For example, a family of human mRNAs has been reported to accumulate at centrosomes and shift localization depending on the cell-cycle stage, revealing a finely orchestrated, translation-dependent program at the centrosome^10^. Similarly, in *S. cerevisiae*, exposure to heat or ethanol stress induces a block in mRNA export, causing bulk poly(A)+ transcripts to accumulate inside the nucleus while stress-response mRNAs are selectively exported, yielding a distinct subcellular signature of the stressed state^11^. Therefore, the ability to characterize and eventually predict the subcellular localization of transcripts is fundamental for advancing our understanding of cellular biology.

While existing subcellular ST methods focus on specific analytic tasks such as identifying genes with distinct subcellular localization patterns^12–14^ or detecting co-localized gene pairs^15^ with shared spatial organization, they fall short of providing a unified framework for comprehensively characterizing and predicting transcript localization in a cell-specific and context-aware manner. This limitation is particularly important because subcellular transcript organization reflects multiple layers of biological context, including coordinated spatial relationships among genes within individual cells, variation in cell-intrinsic states, and context provided by the surrounding tissue microenvironment. Comprehensively modeling subcellular transcript organization therefore calls for several complementary capabilities. First, multi-scale modeling is needed to capture subcellular spatial gene expression while retaining local microenvironmental context, as transcript localization can be influenced by neighboring cells, tissue structure, and extracellular signals. Second, multi-modal integration is required to enable joint modeling of subcellular ST and complementary morphological imaging, providing structural information about cellular and nuclear organization that is not captured by transcriptomic measurements alone. Third, modeling complex multi-gene interactions across hundreds to thousands of genes is critical to enable the discovery of coordinated spatial organization patterns associated with cellular function. Fourth, an ideal model should also represent heterogeneous cellular states rather than being tailored to a single predefined cell type or condition, which requires capturing both population-level variation across cells and within-cell heterogeneity in intracellular organization. Finally, given the complex spatial dependencies underlying subcellular RNA localization, the model should be capable of nonlinear representation learning to establish unified representations across genes and cells and ultimately enable predictive modeling across varying cellular and spatial contexts.

Here, we present Spatially and Virtually Embedded Cell (SVC), a unified artificial intelligence (AI)-based predictive model for subcellular-resolution ST that encompasses the above capacities to capture and predict the subcellular localization of transcripts within individual cells in their spatial tissue context. SVC builds upon a Vision Transformer (ViT)-based framework that enables multi-modal and multi-scale modeling of subcellular ST data. It integrates subcellular transcript localization with cell-level identity and morphological features, while also preserving local spatial context from neighboring cells. In addition, it incorporates prior biological knowledge about gene functional relationships learned from external transcriptomic datasets, while naturally accommodating network connectivity across genes. To learn spatial dependencies underlying subcellular organization in a self-supervised regime, SVC introduces subcellular contextual learning that encodes each gene’s within-cell localization as an image patch and leverages other genes as within-cell context to model cross-gene spatial dependencies. SVC enables spatially grounded virtual representations of individual genes and cells, which is essential for an integrated understanding of cellular function. We demonstrate the utility of SVC through in-depth applications to multiple subcellular-resolution ST datasets.

## Results

### Method overview

SVC builds upon the Vision Transformer (ViT) architecture and adapts it to model subcellular spatial transcriptomics data. A schematic overview of SVC is shown in Fig. 1. At its core, SVC introduces five complementary embeddings to represent genes and cells: (1) gene-level localization embeddings, which capture subcellular spatial expression patterns of individual genes in a cell-specific manner; (2) gene-level functional embeddings, which incorporate prior biological knowledge about gene-gene relationships derived from existing bulk RNA-seq data; (3) cell-level neighbor embeddings, which encode the local environment surrounding each cell through aggregation of expression profiles from its nearest neighbors; (4) cell-level morphology embeddings, which serve as a second modality and encode cellular morphology and nuclear-shape features obtained from parallel DAPI-stained fluorescence or H&E histology image data; (5) cell-level identity embeddings, which are in the form of cell state or cell type labels, capturing heterogeneous cellular characteristics when labels are available. Together, these gene- and cell-level embeddings integrate information across subcellular, cellular, and local tissue scales. These five embeddings provide complementary information, as evidenced by the ablation analyses, with the subcellular gene-level localization embedding serving as the key component that provides predictive subcellular information beyond cell-level features (Supplementary Note 1; Supplementary Fig. 1). Together, they yield a structured representation that enables integrated multi-modal and multi-scale modeling of subcellular spatial transcriptomic data, capturing global tissue architecture as well as fine-grained subcellular molecular localization patterns. A key innovation of SVC lies in analogizing the spatial expression of individual genes within a cell to image patches, enabling the adaptation of ViT’s spatial semantic relationship mining framework towards learning the cell-specific subcellular spatial relationships among genes. SVC learns cell-specific gene relationships through a self-supervised masked gene modeling strategy, supporting a broad range of downstream applications. In particular, SVC predicts subcellular spatial localization patterns for unmeasured genes in new datasets, extending conventional gene imputation and spatial reconstruction to unprecedented subcellular resolution and enabling discovery of biologically meaningful subcellular organization, ranging from polarized transcript distributions and context-dependent shifts in localization to disease-associated remodeling of transcript localization. SVC produces cell-specific gene representations that facilitate subcellular co-localization analysis within individual cells. By capturing multi-gene dependencies, SVC also enables *in silico* prediction of perturbation effects at subcellular resolution with retained spatial variability. Beyond subcellular analysis, by learning how subcellular patterns vary across cells, SVC naturally facilitates cellular- and tissue-scale analyses, distinguishing cellular states and spatial domains and capturing continuous context-associated variation within cell populations and across tissue structures. SVC is computationally scalable, with training time increasing approximately linearly with the number of cells, and can be readily applied to large-scale datasets (Supplementary Table 1; Supplementary Note 2; Supplementary Fig. 2). We demonstrate the utility and benefits of SVC through comprehensive evaluations on multiple subcellular spatial transcriptomics datasets spanning diverse tissue types and technological platforms.

**Fig. 1.**
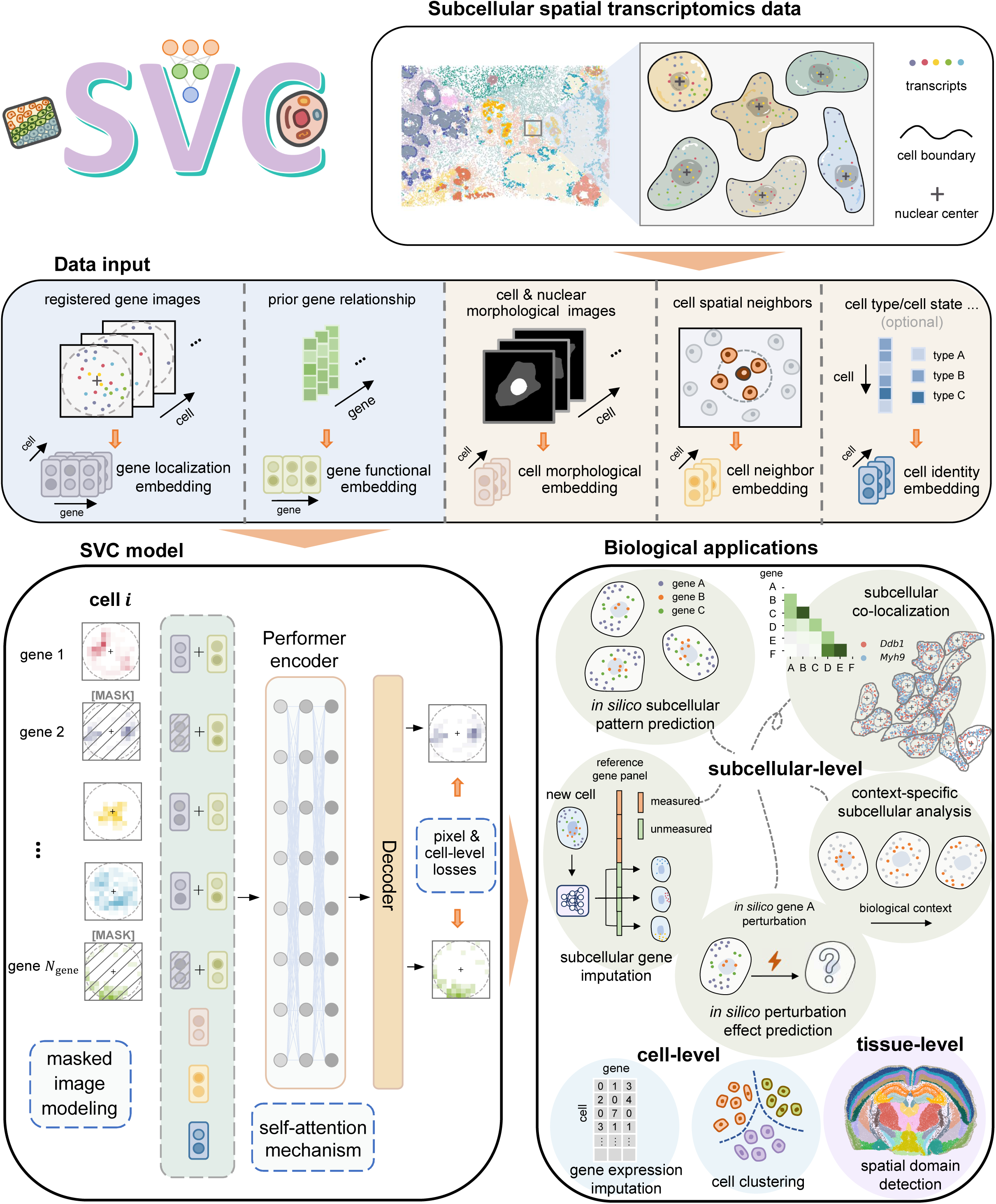
Overview of SVC framework. (1). SVC is a Vision Transformer (ViT)-based spatially and virtually embedded cell model trained on subcellular spatial transcriptomics data that provide fine-grained molecular localization within each cell while retaining higher-level cellular and tissue context. (2). SVC integrates multi-modal and multi-scale information from five complementary inputs: registered gene images that capture gene subcellular spatial localization; prior gene-gene relationships derived from existing pretrained models; images of cell and nuclear morphologies; each cell’s local spatial neighborhood; and optional cell type or state labels. These inputs are mapped to five corresponding embeddings: gene-level localization embeddings and functional embeddings, as well as cell-level morphology, neighbor, and identity embeddings. (3). In SVC, the resulting gene- and cell-level embeddings are combined and fed into a Performer encoder block. SVC is trained using a self-supervised masked image modeling procedure, in which a random subset of gene expression images in each cell is masked. A decoder is then used by SVC to reconstruct their spatial expression patterns by minimizing pixel- and cell-level loss functions, enabling SVC to recover and predict gene-specific subcellular expression patterns and infer spatial relationships across genes. (4). SVC enables a broad range of downstream applications across multiple scales, with a primary focus on subcellular-level analysis. At the subcellular level, it predicts fine-grained expression patterns for unmeasured genes within individual cells; performs spatial expression imputation across genes and cells at subcellular resolution; infers subcellular gene-gene co-localization patterns; characterizes context-specific changes in subcellular organization across different cellular or spatial environments; and supports *in silico* prediction of perturbation-induced changes in subcellular spatial expression. Beyond these subcellular applications, the learned representations can also be used for cell- and tissue-level analyses, including cell-level gene expression imputation, cell clustering, and spatial domain detection.

### SeqFISH+ mouse embryonic fibroblast data

We first applied SVC to a seqFISH+ dataset of NIH/3T3 mouse embryonic fibroblast cells^5^. This dataset measures 171 cells, and we focused the analysis on the top 1,000 most highly expressed genes across 17 field-of-view (FOV) batches. Because this dataset has been extensively studied across prior subcellular ST methods^12–15^, and the original study provides subcellular localization annotations for a subset of genes, we primarily used it for comprehensive benchmarking to evaluate SVC’s capacities in prediction and unified representation learning.

We first assessed SVC’s performance in predicting subcellular spatial patterns for unmeasured mRNAs. Batch 3 was held out as the test set for final evaluation, while batch 1 was used for validation and the remaining batches for training (Methods). Among the 1,000 measured genes in batch 3, we randomly partitioned them into 20 groups and conducted 20-fold masked gene prediction. In each fold, we masked one group of genes, and used the trained SVC model to predict their spatial expression patterns based on the remaining genes. Since no existing methods are capable of predicting subcellular gene expression, we carefully designed four complementary image-based baselines for comparison, representing a randomly selected training-cell image (Baseline 1), a spatially shuffled target-cell image (Baseline 2), and population-average predictions using training cells (Baseline 3) or the remaining test cells (Baseline 4; Methods); Baseline 4 represents the strongest reference baseline but is not feasible in practical imputation settings where the gene is not measured in the test cells. Across three evaluation metrics including root mean squared error (RMSE), Pearson correlation (PCC), and cosine similarity, SVC consistently outperformed all four baselines (Fig. 2a and Supplementary Fig. 3), with median improvements of 29.9-37.1% over Baselines 1-2 and 5.2-21.0% over the more challenging Baselines 3-4. We next quantified similarity between predicted and measured transcript distributions using three distribution-based metrics including Earth Mover’s Distance (EMD), Kullback-Leibler (KL) divergence, and Jensen-Shannon (JS) divergence (Supplementary Fig. 4). SVC again achieved positive median gains over all metrics and baselines, improving EMD by 38.0-39.7% and JS divergence by 25.5-26.7% relative to Baselines 1-2, and by 21.8%/21.3% (EMD) and 12.2%/12.3% (JS divergence) relative to Baselines 3-4. At the gene level, SVC outperformed all baselines for at least 97.3% of genes across all six metrics. Together, these results show that SVC more accurately recovers cell-specific subcellular transcript organization than the baselines.

**Fig. 2.**
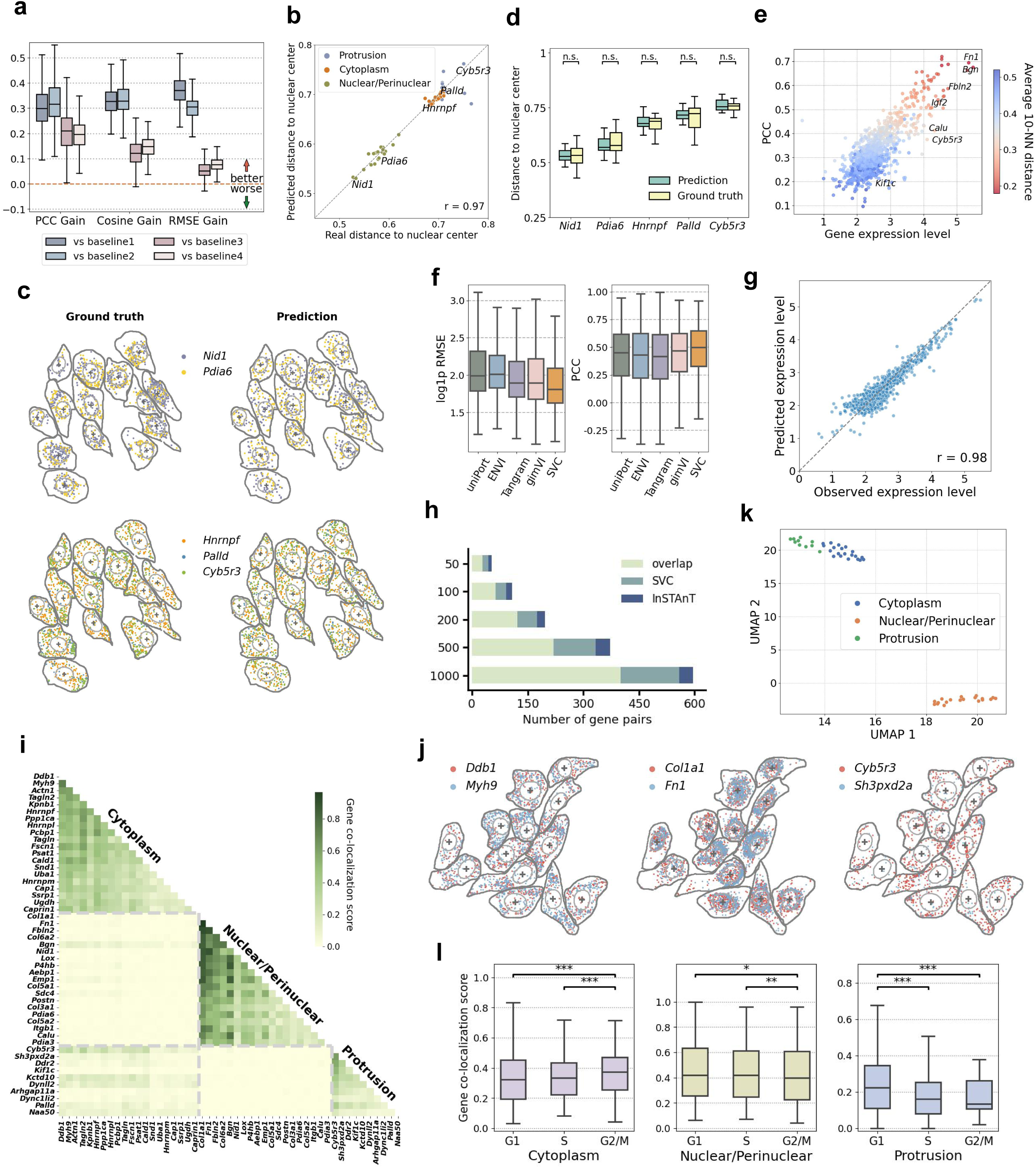
SeqFISH+ mouse embryonic fibroblast data analysis. **a.** Box plots show the per-gene performance gains of SVC over the four baseline models, evaluated using Pearson correlation coefficient (PCC), cosine similarity, and root mean squared error (RMSE). **b.** Scatter plot compares the SVC-predicted relative distance to the nuclear center for 49 genes with well-annotated subcellular localization patterns against the truth values (Pearson correlation = 0.97). Dot colors indicate different types of subcellular patterns. **c.** SVC-predicted transcript profiles are visualized within the original cells alongside the ground truth for five representative genes. **d.** Grouped box plots show the distributions of SVC-predicted relative distance to the nuclear center and the corresponding ground-truth values across cells for each of the five genes. Statistical significance for pairwise comparisons (n.s. ≥ 0.05; * < 0.05; ** < 0.01; *** < 0.001) was assessed using two-sided Wilcoxon signed-rank test *P* values. **e.** Scatter plot shows, for each gene, how subcellular prediction accuracy varies with expression level and latent-space neighborhood similarity, where the *x*-axis shows log1p-transformed mean expression level, the *y*-axis shows the Pearson correlation coefficient between predicted and observed subcellular patterns, and the color scale indicates average distance to the 10 nearest neighbors (10-NN) in latent space. **f.** For the cell-level gene expression prediction task, box plots show the per-gene RMSE (log1p-transformed) and PCC scores between observed and predicted expression counts using 20 held-out gene folds. SVC is compared with uniPort, ENVI, Tangram and gimVI. **g.** Scatter plot shows SVC-predicted vs observed cell-level expression across all genes on a log-transformed scale, with each dot representing a gene. Pearson correlation was calculated on the original scale. **h.** Comparison of the top 50, 100, 200, 500, and 1,000 co-localized gene pairs detected by SVC and InSTAnT. The stacked bars show the number of gene pairs in three categories: (i) overlapping pairs: gene pairs detected by both SVC and InSTAnT; (ii) verified SVC-only pairs: among the non-overlapping pairs detected only by SVC, those in which both genes are among the 49 well-annotated genes and share the same subcellular localization type; and (iii) verified InSTAnT-only pairs: the corresponding category for non-overlapping pairs detected only by InSTAnT. **i.** Heatmap visualizes subcellular co-localization scores for pairs of these 49 genes, displaying three distinct co-localization clusters. **j.** Observed transcript profiles within original cells for three representative co-localization gene pairs. **k.** UMAP visualization of these 49 genes, obtained from the average latent representation of each gene across training cells. Point colors indicate different types of subcellular patterns. **l.** Box plots show the distributions of co-localization scores for cytoplasmic-localized, nuclear/perinuclear-localized, and protrusion-related genes averaged across cells in G1, S, and G2/M phase. Statistical significance for pairwise comparisons (n.s. ≥ 0.05; * < 0.05; ** < 0.01; *** < 0.001) was assessed using one-sided Wilcoxon signed-rank tests.

Next, we systematically examined whether SVC recovered previously characterized subcellular localization patterns. Specifically, among the analyzed genes, 49 had been previously documented^5^ to exhibit one of three distinct subcellular localization patterns: nuclear/perinuclear, cytoplasmic, and protrusion. To facilitate the comparison with these previously documented patterns, we computed the relative distance of the predicted expression values from the nuclear center for each gene: a smaller distance indicates a nuclear/perinuclear pattern, an intermediate distance corresponds to a cytoplasmic pattern, and a larger distance suggests protrusion localization (Methods). As expected, SVC’s predicted subcellular spatial patterns closely matched these known patterns (Fig. 2b), demonstrating SVC’s ability to accurately recover gene-specific localization. In addition, we observed a strong correlation of the predicted distances with those computed from ground truth spatial patterns (Pearson correlation = 0.97). For example, SVC successfully recapitulated the subcellular spatial distributions of several genes: *Nid1* and *Pdia6*, which were previously documented to concentrate near nuclear centroids; *Cyb5r3*, which is known to localize to the cellular peripheries; and *Hnrnpf* and *Palld*, which are known to distribute in intermediate zones (Fig. 2c); all with similar predicted and observed relative distances to the nuclear center (Paired Wilcoxon tests, all *P* values > 0.05; Fig. 2d). Importantly, SVC is particularly effective at predicting genes that are highly expressed or share similar latent representations with other genes (Fig. 2e). To assess whether SVC prediction performance extends to sparse genes, we further evaluated prediction accuracy across the full expression range. SVC maintained a consistent performance advantage over all four baselines across expression groups, including low-expression genes (Supplementary Note 3; Supplementary Fig. 5). Comparisons with gene-specific mean-distance baselines further showed that SVC captures cell-to-cell variation in nuclear distance beyond gene-level average localization (Supplementary Note 4; Supplementary Fig. 6). These results highlight SVC’s high fidelity in capturing gene-specific subcellular localization.

Because SVC predicts fine-grained subcellular patterns that can be aggregated to obtain cell-level expression, we also assessed its performance on cell-level gene imputation, a related but less complex task that provides complementary evidence for its predictive performance across scales. Specifically, we compared SVC with four state-of-the-art spatial-transcriptomic integration methods, ENVI^16^, Tangram^17^, gimVI^18^, and uniPort^19^, all of which leverage single-cell RNA-seq data to impute unmeasured gene expression for spatial transcriptomics at the cellular level. For a fair comparison, key hyperparameters for all competing methods were selected using the validation data before evaluation on the test set (Methods; Supplementary Note 5; Supplementary Fig. 7). Notably, although SVC was not explicitly designed for cell-level imputation, it demonstrated competitive performance across all genes, achieving the lowest median RMSE and highest median PCC among the evaluated methods (Fig. 2f): the median RMSE values across all the genes for uniPort, ENVI, Tangram, gimVI and SVC were 6.33, 6.47, 5.64, 5.66 and 5.12, respectively, while the corresponding median PCC values were 0.45, 0.43, 0.42, 0.47, and 0.50. Additionally, the predicted cellular expression values by SVC closely aligned with ground truth across genes (Pearson correlation = 0.98; Fig. 2g), compared to uniPort (Pearson correlation = 0.90; Supplementary Fig. 8), ENVI (Pearson correlation = 0.97), Tangram (Pearson correlation = 0.95), and gimVI (Pearson correlation = 0.87). The cell-level expression prediction result highlights SVC’s predictive performance, even in settings beyond its primary design scope.

Beyond predicting subcellular spatial expression, SVC integrates multi-scale and multi-modal information to learn unified, spatially grounded virtual representations of cells and genes. The similarities among the inferred representations across genes enable subcellular gene co-localization analysis to capture spatial co-organization within cells, which is distinct from conventional cell-level co-expression or covariation analysis conducted across cells (Methods). The resulting co-localization analysis in SVC also differs from that of InSTAnT^15^ in two key aspects. First, SVC assesses co-localization based on each gene’s overall subcellular localization pattern within cells, whereas InSTAnT primarily captures short-range co-localization between individual transcripts. Second, SVC captures co-localization across multiple genes, as its latent space supports simultaneous modeling of multi-gene relationships, whereas InSTAnT is restricted to pairwise gene analysis. Despite these differences in analytical focus, the two methods show substantial concordance, with 401 of the top 1,000 gene pairs shared (Fig. 2h). For the gene pairs unique to either method, we compared them against the 49 genes with previously annotated subcellular localization patterns, including 19 nuclear/perinuclear, 20 cytoplasmic, and 10 protrusion-associated genes. These annotations define 406 possible gene pairs with known co-localization patterns as ground truth. Among these, a substantially larger number of SVC-specific pairs were consistent with the ground truth (158 by SVC vs 36 by InSTAnT), supporting SVC’s ability to capture gene-specific overall spatial patterns within cells. Furthermore, the gene co-localization relationships inferred on the 49 genes by SVC grouped into three distinct spatial localization clusters, consistent with their annotated patterns (Fig. 2i-j; Supplementary Fig. 9). UMAP visualization of their latent representation reveals corresponding separation (Fig. 2k), while cosine distances between cluster-average representations in the learned latent space (nuclear/perinuclear vs cytoplasmic: 0.13, nuclear/perinuclear vs protrusion: 0.35, cytoplasmic vs protrusion: 0.18) quantify the relative proximity among clusters. Beyond these 49 genes, the original study identified 11 additional genes that formed three nuclear-associated subclusters^5^ (Supplementary Fig. 10). The SVC-inferred latent representations preserved these three subclusters, with closer distances among genes within the same subcluster compared to those from different subclusters (Supplementary Figs. 11 and 12). These results indicate that SVC’s virtual representation effectively preserves key subcellular localization features of genes.

An important feature of the dataset is that the NIH/3T3 embryonic fibroblast cells are actively undergoing the cell cycle, with individual cells captured at different phases. This variability provides an opportunity to examine whether SVC captures biologically plausible cell-cycle-associated variation in subcellular transcript organization. Therefore, we further incorporated ELLA^14^ cell-cycle annotations into SVC’s cell-identity embeddings, enabling SVC to effectively capture cell-cycle-dependent transcriptional differences. Notably, SVC revealed clear differences in subcellular gene co-localization across G1, S, and G2/M phases, consistent with expected transcript redistribution during cell-cycle progression (Fig. 2l). Cytoplasm-localized genes exhibited the highest total co-localization scores during the G2/M phase (Paired Wilcoxon tests: G2/M vs G1, *P* value = 1 × 10^−4^ ; G2/M vs S, *P* value = 6 × 10^−5^), consistent with cell-cycle-associated remodeling during mitosis. Conversely, nuclear-localized genes exhibited stronger co-localization during the G1 and S phases compared to G2/M (G1 vs G2/M, *P* value = 0.025; S vs G2/M, *P* value = 7 × 10^−3^), which may reflect greater nuclear transcriptional and RNA-processing activity during G1 and S relative to G2/M. Additionally, protrusion-related genes showed significantly higher co-localization in G1 cells (G1 vs S, *P* value = 1 × 10^−7^ ; G1 vs G2/M, *P* value = 1 × 10^−7^), consistent with enhanced cell motility and cytoskeletal activity during this phase. These results provide biological validation that SVC captures dynamic changes in subcellular gene co-localization across the cell cycle that are consistent with phase-specific cellular organization and activities. Because cell-cycle progression is closely accompanied by changes in cell and nuclear morphology, we further examined whether the observed co-localization differences can be accounted for by morphology. This is important because systematic shape and size changes across cell-cycle stages could, in principle, alter measured subcellular localization patterns (e.g., via shifts in compartment boundaries or segmentation) even if the underlying RNA organization were unchanged. We performed complementary morphology controls by (i) removing morphology inputs from SVC or (ii) residualizing the latent representations against morphology-derived features (Supplementary Note 6; Supplementary Figs. 13-15). Most cell-cycle-associated differences persisted, with 11 of the 12 comparisons remaining significant; the main exception was the G2/M-versus-S difference for cytoplasm-localized genes, which was attenuated after morphology residualization, suggesting a contribution from morphology-associated changes. Overall, these results show that SVC reliably captures population-level variation in subcellular transcript localization across cell-cycle states.

### MERFISH human osteosarcoma data

Next, we applied SVC to a MERFISH dataset of human osteosarcoma U2-OS cells^20^, from which we analyzed 1,022 cells and 119 genes across 70 imaging FOV batches.

We began by evaluating SVC’s predictive accuracy across scales, starting at the subcellular level and then extending to the cellular level. To assess its ability to capture subcellular localization patterns on the test data, we visualized spatial predictions and evaluated quantitative performance across genes (Fig. 3a-b; Supplementary Figs. 16 and 17). Across six evaluation metrics, SVC consistently outperformed all four baselines, achieving median performance gains of 26.4-87.0% over Baselines 1-2 and 3.4-14.3% over the stronger Baselines 3 and 4. At the gene level, SVC outperformed all baselines for at least 99.2% of genes. Additionally, the predicted relative distances to the nuclear center matched the observed ones across genes (Pearson correlation = 0.99; Supplementary Fig. 18), with mean-distance baseline comparisons further supporting SVC’s ability to capture cell-to-cell variation in subcellular distribution beyond gene-level averages (Supplementary Fig. 19). Predicted and observed spatial patterns for five example genes (*SLC5A3*, *SRRM2*, *COL5A1*, *USP9X*, and *TLN1*) showed strong concordance, capturing both their progressively increasing radial distances from the nuclear center and gene-specific concentration patterns (Fig. 3a; Supplementary Fig. 20). Besides reliably predicting subcellular distributions of unmeasured genes, SVC’s aggregated predictions across subcellular locations accurately captured cell-level expression, correlating strongly with observed measurements across all genes (Pearson correlation = 0.99; Fig. 3c). In comparison with existing cell-level imputation methods, SVC showed competitive performance in both RMSE and PCC, with a median RMSE of 22.59 and PCC of 0.60, compared with 26.46/0.60 for uniPort, 24.77/0.61 for ENVI, 28.57/0.39 for Tangram, and 25.43/0.57 for gimVI (Supplementary Fig. 21).

**Fig. 3.**
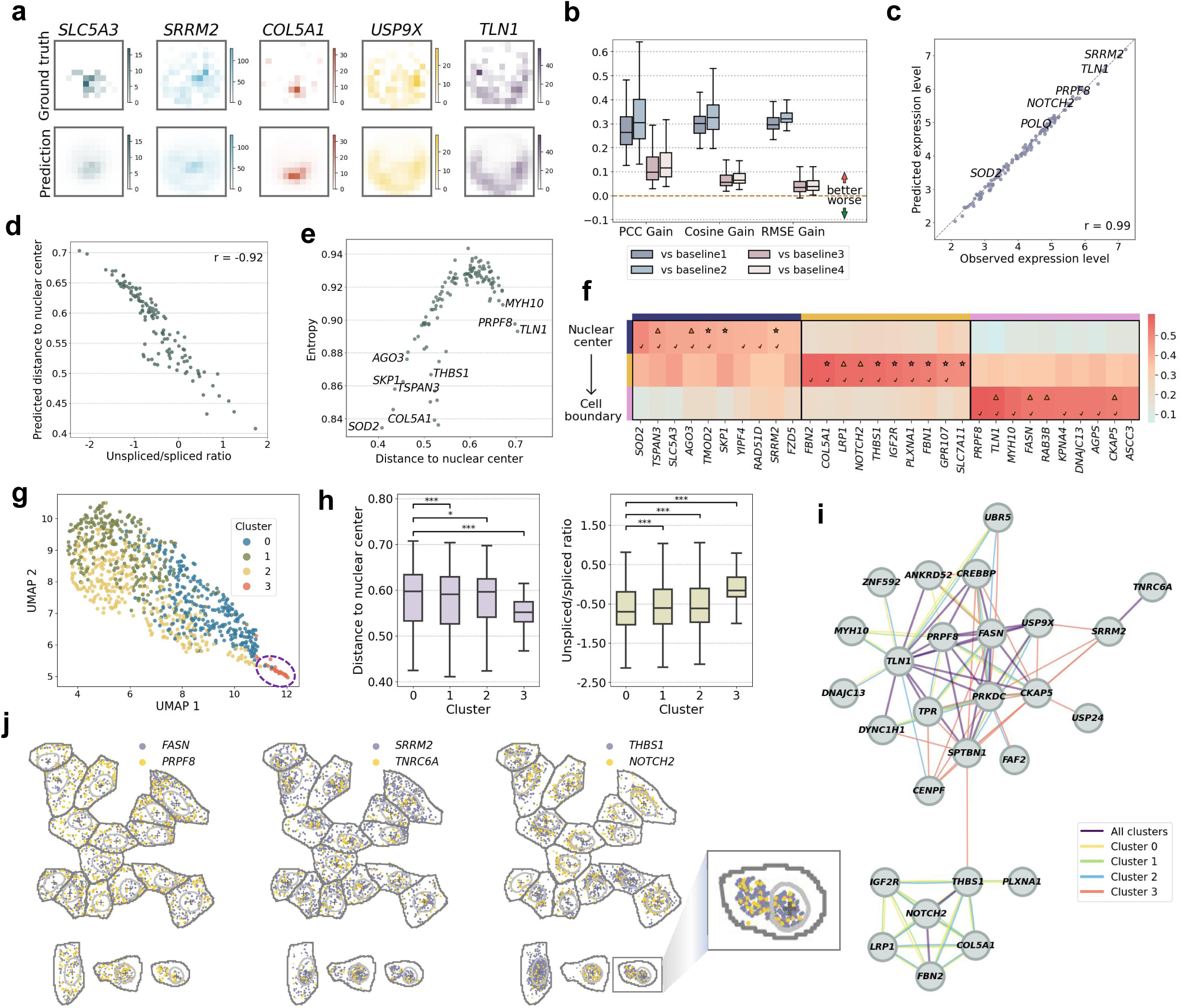
MERFISH human osteosarcoma data analysis. a. Visualization of SVC-predicted subcellular expression patterns, represented as 12 × 12 registered images, alongside the ground truth for five genes within a cell. **b.** Box plots show the per-gene performance gains of SVC over the four baseline models, evaluated using Pearson correlation coefficient (PCC), cosine similarity, and root mean squared error (RMSE). **c.** Scatter plot shows SVC-predicted vs observed cell-level expression across all genes on a log-transformed scale, with each dot representing a gene. Pearson correlation was calculated on the original scale. **d.** Scatter plot shows each gene’s SVC-predicted relative distance to the nuclear center against its average log1p-transformed unspliced-to-spliced ratio (Pearson correlation = −0.92). **e.** Scatter plot depicts each gene’s entropy vs its SVC-predicted relative distance to the nuclear center. Entropy was calculated as the normalized Shannon entropy of SVC-predicted expression across image pixels, ranging from 0 to 1, with low values indicating stronger concentration within specific subregions and high values indicating a more diffuse distribution across subregions. **f.** Heatmap displays the top 10 genes with the highest subregion-specific enrichment. Enrichment is computed for each gene as the average proportion of SVC-predicted transcripts within radial subregions from the nuclear center to the cell boundary across cells. Check marks in the heatmap indicate genes that overlap with the top 10 subregion-specific genes obtained from the observed real images, whereas stars and triangles denote genes whose encoded protein localizations in the Human Protein Atlas (HPA) are consistent with our annotated subregions in U2-OS cells or in other cell lines, respectively. **g.** UMAP visualizes cell clustering results from SVC. For each cell, SVC computed a gene-gene cosine similarity matrix from its inferred latent representation, flattened it into a feature vector, and used this vector as input to UMAP. **h.** Box plots show the average relative distance to the nuclear center (left) and unspliced/spliced expression ratio (log1p-transformed; right) across genes in each of the clusters 0-3. Statistical significance for pairwise comparisons (n.s. ≥ 0.05; * < 0.05; ** < 0.01; *** < 0.001) is based on one-sided Wilcoxon signed-rank tests. **i.** Gene co-localization network for the top 50 gene pairs in each cell cluster, with edge colors denoting the cluster assignment of each identified gene pair. Gene pairs shared among all clusters are connected by purple edges. **j.** Observed transcript profiles in the original cells for three representative co-localized gene pairs shared by all cell clusters. Top panels show cells from clusters 0, 1, and 2; bottom panels show three example cells from the abnormal cluster 3.

We further validated SVC’s accuracy in predicting subcellular mRNA localization by examining its biological implications. First, unspliced transcripts are known to remain near their transcription sites within the nucleus. Accordingly, for each gene, we calculated its predicted relative distance to the nuclear center and its average unspliced-to-spliced ratio. These two values are strongly negatively correlated (Pearson correlation = −0.92; Fig. 3d), confirming the accuracy of SVC. Second, we characterized each gene’s predicted subcellular distribution using spatial localization entropy. Lower entropy indicates that expression is more strongly concentrated within specific subcellular regions, whereas higher entropy reflects a more diffuse distribution across the cell. We found that genes concentrated near the nuclear region generally exhibit lower entropy; however, genes with similar relative distances to the nuclear center can display substantially different entropy values, reflecting finer-grained variation in their subcellular distribution patterns (Fig. 3e). To further characterize these concentration patterns, we performed GO enrichment analysis using Metascape^21^ on the top 10 low-entropy and high-entropy genes (Supplementary Fig. 22). Low-entropy genes were enriched for functions related to apoptotic regulation, proteoglycan binding, serine/threonine receptor signaling, and calcium ion binding (*P* value < 0.01), indicating that genes with more spatially concentrated mRNA distributions are associated with diverse signaling, binding, and regulatory functions. In contrast, no GO terms with *P* value < 0.01 were identified for the high-entropy genes, suggesting that no dominant functional program was evident among genes with more diffuse mRNA distributions.

We next assessed subregion-specific enrichment of SVC-predicted patterns across nuclear, cytoplasmic, and membrane-proximal regions (Methods). The top 10 genes with the highest enrichment scores in each category are shown in Fig. 3f. Notably, most of these genes (25 out of 30; 83.3%) also appear among the top 10 compartment-specific genes identified using observed, rather than predicted, gene expression. A substantial fraction of these genes show patterns consistent with their protein localizations reported in the Human Protein Atlas (HPA, Methods). Specifically, 10 genes (33.3%) exhibit concordant localization in the U2-OS cell line and an additional 8 genes (26.7%) match protein locations in at least one other cell line, supporting the notion of localized translation^22^. Some genes, however, display discrepancies between mRNA and protein localization. For example, the secreted protein FBN2 has mRNA that is localized intracellularly, as it is synthesized inside the cell, but the protein itself is secreted and found in the extracellular matrix^23^. In conclusion, protein-localization data from HPA provide independent and supportive evidence for the accuracy of SVC’s predicted mRNA localization patterns, while also underscoring the diverse and intricate mechanisms that link gene transcription to protein translation. Robustness analyses further showed that SVC prediction performance remained stable under substantial perturbations in RNA coordinates and morphological inputs (Supplementary Note 7; Supplementary Figs. 23-26).

Importantly, SVC learns cell-specific representations of multi-gene subcellular organization that can be used to characterize cell-state heterogeneity. Given the genomic instability and pronounced subpopulation heterogeneity of U2-OS cells, we explored the utility of SVC’s representations for studying cell states and cell-to-cell variability through cell clustering (Methods). Unlike conventional clustering methods relying solely on cell-level gene expression, SVC incorporates subcellular spatial localization information across genes, enabling the identification of cell subpopulations with distinct subcellular spatial organization. The clustering results visualized by UMAP (Fig. 3g) show four clusters, with cluster 3 forming a relatively small group. We annotated these clusters using multiple sources of information, including phase-specific marker genes, *CKAP5* (a G2/M phase marker) and *BRCA2* (an S/G2 phase marker) (Supplementary Fig. 27), as well as genes’ relative distance to the nuclear center and their average unspliced/spliced ratio (Fig. 3h). Cluster 0 is annotated as G2/M phase, showing high *CKAP5* and *BRCA2* expression, a modest shift toward more peripheral transcript localization, and the lowest unspliced/spliced ratio, consistent with mitosis-associated nuclear reorganization and a greater proportion of mature, fully spliced transcripts during mitosis. Cluster 1 corresponded to S phase, exhibiting high *BRCA2* expression and a slightly more nuclear-proximal transcript distribution. Cluster 2 is annotated as G1 phase, characterized by low expression of both *CKAP5* and *BRCA2*, together with an intermediate radial transcript distribution and unspliced/spliced ratio. However, cluster 3 exhibited concentrated nuclear-proximal transcript localization, a markedly elevated unspliced/spliced ratio, and increased *CKAP5* expression. These mixed features suggest that cluster 3 represents an atypical or undefined cell state rather than a canonical cell-cycle phase. Subcellular gene co-localization analysis further highlighted the distinct spatial organization of cluster 3 (Fig. 3i). Notably, cluster 3 contains 19 unique gene pairs, while clusters 0, 1, and 2 contain 1, 0, and 1 unique pairs, respectively, indicating that the co-localization patterns of cluster 3 are distinct from the others. Visualization of three co-localized gene pairs further showed distinct spatial distributions in cluster 3 cells (Fig. 3j). In clusters 0-2, for example, the *FASN*-*PRPF8* pair was concentrated near cell boundaries, *SRRM2*-*TNRC6A* was primarily distributed in the nucleus with some cytoplasmic transcripts, and *THBS1*-*NOTCH2* was localized predominantly at the nuclear edge and in cytoplasmic regions. While these same gene pairs are also co-localized in cluster 3 cells, their spatial distributions differ substantially, showing consistent nuclear-proximal enrichment. A zoomed-in example reveals a cell that appears to be multinucleated, suggesting a potential abnormal or stressed state. Finally, we performed GO enrichment analysis via Metascape^21^ on genes involved in the 19 cluster-3-specific gene pairs (Supplementary Fig. 28). The enriched GO terms were primarily related to mitotic cell-cycle regulation and microtubule cytoskeleton organization, suggesting altered mitotic regulation and supporting the interpretation of cluster 3 as a potentially atypical cell-cycle-associated state. The SVC-defined clusters are not primarily explained by cell and nuclear morphology, as morphology-only features showed little separation among these clusters, while cluster-associated information persisted after accounting for morphology, supporting that the SVC clusters capture variation beyond morphology (Supplementary Note 8; Supplementary Figs. 29 and 30). This case study illustrates SVC’s capacity to sensitively discern cell states via distinctive subcellular organization.

### Xenium mouse brain data

Next, we focused on two large-scale Xenium tissue datasets to highlight SVC’s ability to jointly capture fine-grained molecular localization within cells and higher-order spatial organization across cellular and tissue contexts, thus enabling distinct downstream analysis.

We first examined the 10x Xenium mouse brain dataset. Tissues consist of spatially organized and functionally distinct anatomical structures, often referred to as spatial domains. Because SVC’s latent representations capture subcellular expression localization information across cells within a tissue, we explored their utility for spatial domain detection. In particular, we combined SVC-derived gene-gene similarity with the physical proximity between cells to infer spatial domains (Methods). We found that the domains identified by SVC aligned well with major anatomical brain regions (Fig. 4a-b and Supplementary Fig. 31) and were supported by the expression patterns of canonical marker genes (Fig. 4c), highlighting its ability to resolve biologically meaningful structures from the data. For instance, *Rasgrf2*, *Rorb*, *Fezf2*, *Tle4*, and *Cplx3*, which mark isocortical layers 2/3, 4, 5, 6, and 6b, respectively, were highly enriched in domains 1, 14, 15, 3, and 24. The CA1-specific marker *Wfs1* showed strong enrichment in domain 11 relative to domains 13 and 8, whereas CA3-specific marker *Prss35* was predominantly expressed in domain 13 and the dentate gyrus (DG) marker *Prox1* in domain 8. We further evaluated SVC on the held-out section 2 to assess its predictive performance in this heterogeneous tissue setting. SVC consistently outperformed the four baselines in subcellular localization prediction (Supplementary Note 9; Supplementary Fig. 32), preserved cell-type-specific variation in both gene expression and transcript localization, and substantially improved over the gene-mean-distance baseline (Supplementary Figs. 33-35). It also achieved competitive performance relative to existing methods for cell-level gene-expression imputation (Supplementary Fig. 36). Beyond predictive accuracy, bidirectional cross-modal retrieval between cellular morphology and subcellular gene organization further showed that the learned representations in SVC capture relationships across modalities (Supplementary Note 10; Supplementary Figs. 37 and 38).

**Fig. 4.**
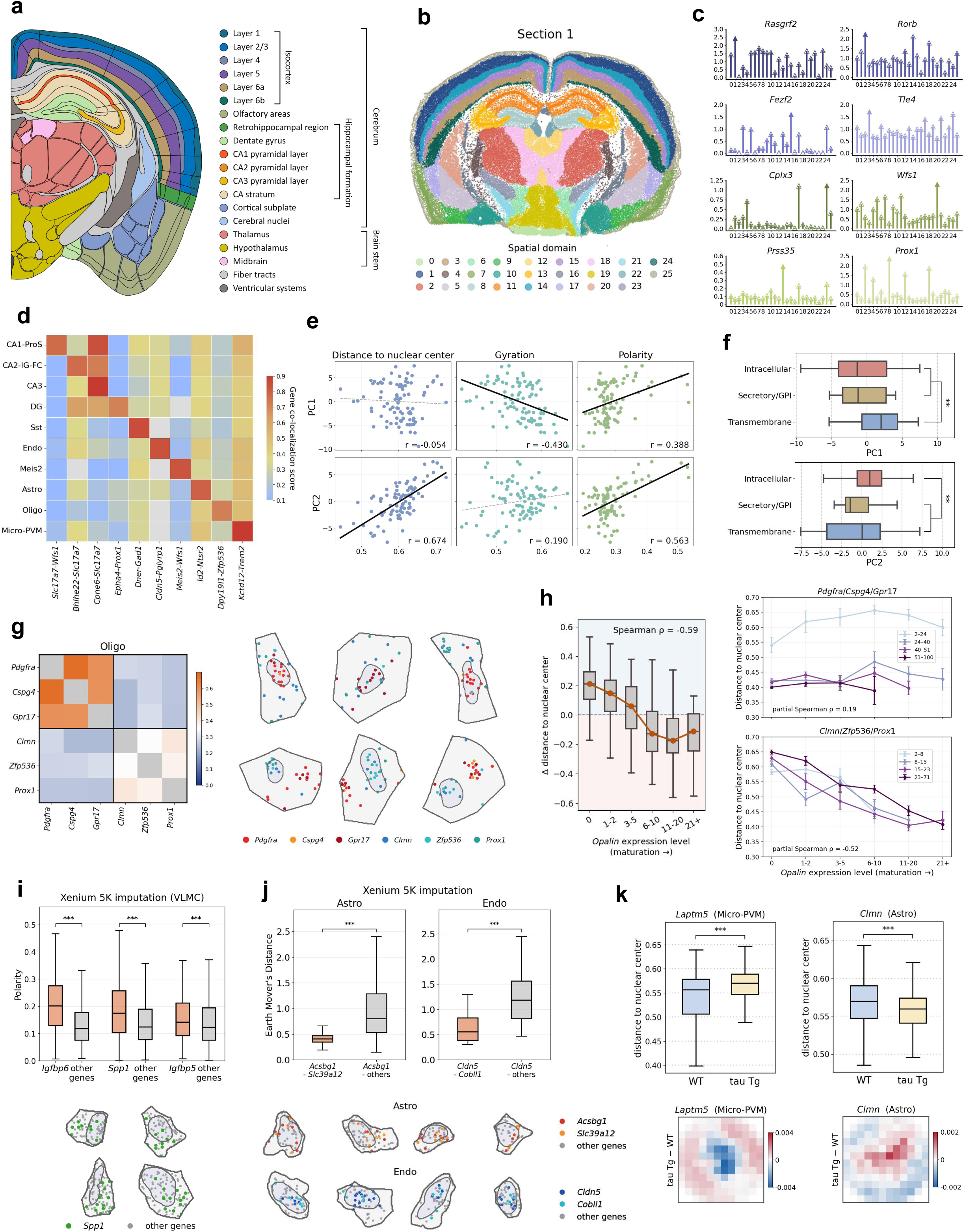
Xenium mouse brain data analysis. **a.** Coronal section of the mouse brain with major tissue regions annotated from the Allen Reference Atlas. **b.** Spatial domains detected by SVC in tissue section 1; results for section 2 are shown in Supplementary Fig. 31. **c.** Lollipop plots display the mean expression (log1p-transformed) of eight marker genes (*y* axis) in each spatial domain (*x* axis). The solid arrow indicates the highest value. **d.** Heatmap visualizes gene co-localization scores of cell-type-specific gene pairs (columns) across ten cell types (rows). For each cell type, the pair shown is the one whose score most exceeds its average across cell types. Grey indicates pairs with an insufficiently detected gene in that cell type. **e.** Scatter plots show the first two principal components (PCs) of the SVC gene-level latent representation in VLMCs and three measured descriptors: relative distance to the nuclear center, gyration and polarity. Each panel is annotated with the Pearson correlation coefficient (r); the line shows the ordinary least-squares fit of PC score on the corresponding descriptor, with solid lines for |r| ≥ 0.30 and dashed lines for |r| < 0.30. **f.** Box plots show the distributions of PC1 and PC2 scores for genes grouped by protein targeting class, annotated using reviewed mouse Swiss-Prot entries as transmembrane, secretory/GPI-anchored, or intracellular. Significant comparisons are indicated for each PC. Statistical significance for the one-versus-rest comparisons (n.s. ≥ 0.05; * < 0.05; ** < 0.01; *** < 0.001) was assessed using two-sided Mann-Whitney U tests. **g.** Left panel: heatmap showing the mean per-cell cosine similarity between SVC latent representations of six oligodendrocyte-lineage genes, calculated across oligodendrocytes in which all six genes were detected. The first three genes constitute the OPC-associated module (*Pdgfra*, *Cspg4*, *Gpr17*), whereas the latter three constitute the maturation-associated module (*Clmn*, *Zfp536*, *Prox1*). Right panel: observed transcript positions of the six genes in six representative oligodendrocytes; the top row shows representative cells with more nuclear-proximal localization of the OPC-associated module, whereas the bottom row shows cells with more nuclear-proximal localization of the maturation-associated module. **h.** Left panel: box plots show the per-cell difference in the relative distance to the nuclear center between the two modules (maturation-associated minus OPC), stratified by *Opalin* expression level. *Opalin*, a canonical myelin gene excluded from both modules, serves as an independent marker of oligodendrocyte maturation. The orange line connects the median of each bin; across individual cells, the difference correlates with *Opalin* transcript count at Spearman correlation = −0.59. Right panel: mean relative distance to the nuclear center for the OPC module (top) and maturation-associated module (bottom) across *Opalin* expression levels, with cells stratified by quartiles of the corresponding module transcript count; error bars indicate the standard error of the mean. Partial Spearman correlations with *Opalin* expression, controlling for module transcript-count rank, are 0.19 and −0.52 for the OPC and maturation-associated modules, respectively. **i**. Upper panel: box plots showing the predicted per-cell polarity of three unmeasured genes in VLMCs from the Xenium 5K section. Each gene was compared with a count-matched background constructed from other imputed genes by resampling to match the target gene’s per-cell transcript-count distribution. Statistical significance (n.s. ≥ 0.05; * < 0.05; ** < 0.01; *** < 0.001) was assessed using one-sided Mann-Whitney U tests, with Benjamini-Hochberg correction across genes. Lower panel: predicted transcript positions of *Spp1* and of every other imputed gene in four representative cells. **j**. Upper panels: box plots showing the per-cell Earth Mover’s Distance between the predicted subcellular distributions of each imputed gene pair in astrocytes (Astro, left) and endothelial cells (Endo, right) from the Xenium 5K section. Each pair was compared with a count-matched background of the same gene paired with every other imputed gene, resampled to match the pair’s per-cell total transcript count. Statistical significance (n.s. ≥ 0.05; * < 0.05; ** < 0.01; *** < 0.001) was assessed using one-sided Mann-Whitney U tests, with Benjamini-Hochberg correction across gene pairs. Lower panels: predicted transcript positions of each pair and of every other imputed gene in four representative astrocytes and four endothelial cells. **k**. Upper panels: box plots showing the predicted relative distance to the nuclear center of one imputed gene in Micro-PVM cells (left) and astrocytes (Astro, right) from wild-type (WT) and P301S tau-transgenic (tau Tg) sections. Statistical significance (n.s. ≥ 0.05; * < 0.05; ** < 0.01; *** < 0.001) was assessed using two-sided Mann-Whitney U tests, with Benjamini-Hochberg correction across genes within each cell type. Lower panels: differences in the SVC-predicted subcellular distributions between tau Tg and WT cells, averaged over cells and shown as the per-pixel change in mass fraction; each cell’s map was normalized to sum to one before averaging.

SVC further enables cell-type-resolved analysis of subcellular gene co-localization, allowing us to identify gene pairs with pronounced cell-type co-localization specificity in the mouse brain that may reflect distinct cellular functions and molecular programs. Across the ten cell types analyzed, we observed marked variation in subcellular co-localization patterns among cell populations (Fig. 4d). For example, *Dner*-*Gad1* showed preferential co-localization in Sst interneurons, likely reflecting coordinated subcellular organization of dendritic and GABAergic inhibitory programs characteristic of this interneuron population^24, 25^. *Cldn5* and *Pglyrp1* exhibited prominent co-localization in endothelial cells, capturing the joint organization of barrier-maintenance and innate-defense programs in brain endothelium^26, 27^. A within-cell spatial-shuffle test showed that all ten pairs exhibited significant subcellular co-localization beyond that expected from cell-level co-expression alone (empirical *P* value < 0.001), consistent with our finding that evidence for subcellular co-localization and for cell-level co-expression is not strongly correlated across gene pairs within individual cell types (Supplementary Note 11; Supplementary Figs. 39 and 40).

The gene-specific representations inferred by SVC are biologically interpretable and can help uncover fine-grained continuous variation in subcellular RNA organization within a single cell population *de novo*. To see this, we focused on 6,613 vascular and leptomeningeal cells (VLMCs) and performed principal component analysis (PCA) on gene-specific SVC representations after removing shared cell-level and transcript-abundance-associated variation (Methods). The first two principal components explained 17.8% and 12.0% of the variation, respectively, revealing two dominant continuous axes of gene-level variation in subcellular organization (Supplementary Fig. 41). PC2 captured a radial localization gradient, with strong associations with nuclear distance and polarity (Pearson correlation = 0.674 and 0.563, respectively), whereas PC1 showed little association with nuclear distance but was associated with polarity and gyration (Pearson correlation = −0.054, 0.388, and −0.430, respectively; Methods; Fig. 4e). Given the prominent role of VLMCs in extracellular-matrix synthesis and matrix-mediated interactions^28, 29^, we grouped genes into intracellular, secretory/GPI-anchored, and transmembrane protein-targeting classes based on UniProt annotations^30^ to examine the functional relevance of the SVC-identified organization signatures (Methods). These protein-targeting classes separated along the SVC-derived axes (Fig. 4f): transmembrane genes showed higher PC1 scores than the other two classes (*P* value = 0.0084), whereas intracellular genes showed higher PC2 scores than secretory/GPI-anchored and transmembrane genes (*P* value = 0.0095). Consistently, transmembrane genes showed lower mean gyration (0.525 vs 0.560 and 0.558 for intracellular and secretory/GPI-anchored genes), whereas intracellular genes showed greater mean nuclear distance (0.614 vs 0.600 and 0.586 for secretory/GPI-anchored and transmembrane genes). These patterns are biologically insightful given the preferential ER association of mRNAs encoding secretory and membrane proteins and the spatially restricted localization of actively translated secretome mRNAs at specific ER junctions^4, 31, 32^. Their separation along the SVC-derived principal components therefore suggests that SVC uncovers biologically interpretable continuous variation in subcellular RNA organization that aligns with distinct protein-biogenesis and targeting requirements.

We focused on oligodendrocytes as a case study to further illustrate the benefits of SVC for discovering subcellular organization of gene programs. Specifically, we examined two biologically motivated gene modules along the oligodendrocyte lineage: an oligodendrocyte precursor cell (OPC)-associated module comprising *Pdgfra*, *Cspg4*, and *Gpr17*, and a maturation-associated module comprising *Clmn*, *Zfp536*, and *Prox1*. These genes have established roles in precursor identity, differentiation, and mature myelinating function^33–36^. Gene-specific SVC representations in oligodendrocytes showed greater similarity among genes within each module than between modules (Methods; Fig. 4g, left). Despite within-module similarity, visualization of representative cells illustrated variability in the relative radial positioning of the two modules (Fig. 4g, right). In particular, the relative positioning score progressively decreased with increasing *Opalin* expression (Methods), an independent marker of mature oligodendrocytes, shifting from positive to negative values (Spearman correlation = −0.59; Fig. 4h, left), indicating continuous radial repositioning of the two modules during maturation. *In situ* mapping further showed that this variation was spatially organized across the tissue (Supplementary Fig. 42), with positive scores enriched in deep central regions and negative scores toward the outer regions. Because oligodendrocyte maturation is accompanied by expected changes in module expression (Supplementary Fig. 43), we asked whether the observed radial shifts persisted after accounting for cell-level transcript abundance (Fig. 4h, right). The maturation-associated module remained strongly associated with *Opalin* expression after controlling for module abundance (partial Spearman correlation = −0.52), whereas the OPC-associated module showed only a modest positive association (partial Spearman correlation = 0.19) and greater dependence on transcript abundance. Together, these results indicate coordinated remodeling of subcellular transcript localization during maturation beyond changes in cell-level transcript abundance. This pattern is directionally consistent with prior protein-level reports that PROX1 becomes increasingly nuclear^37^, whereas GPR17 redistributes from intracellular/perinuclear compartments toward cellular processes and the plasma membrane during oligodendrocyte lineage progression^38^. Our findings provide complementary mRNA-level evidence for coordinated maturation-associated spatial reorganization, potentially involving mRNA sorting and transport^39, 40^.

Lastly, we demonstrate that SVC extends conventional gene imputation from the cell level to subcellular resolution. Cross-dataset transfer of mouse brain data from Xenium to MERFISH and cross-tissue transfer from human breast cancer to human lung cancer demonstrated robust generalization of SVC for subcellular localization prediction across independent datasets and biological contexts (Supplementary Note 12; Supplementary Figs. 44-47). We also applied the trained model to a newly released mouse brain hemisphere dataset generated with the 10x Xenium Prime 5K Mouse Pan Tissue and Pathways Panel to evaluate its ability to impute subcellular localization patterns of unmeasured genes, addressing a challenging yet practical generalization task. The Xenium 5K dataset was substantially sparser than the Xenium mouse brain dataset, with an average of 0.56 vs 1.74 transcripts per gene per cell. In addition, because the 5K panel was designed for broad cross-tissue applications, several brain-specific genes relevant to regional and cell-type-specific biology were not included in the measured panel. In the analysis, SVC successfully predicted the subcellular expression patterns of 84 unmeasured genes across the entire tissue section. As a qualitative assessment of prediction plausibility, several unmeasured anatomical marker genes showed the expected tissue-level enrichment patterns: *Rasgrf2* and *Rprm* were enriched in isocortical layers 2/3 and 6, respectively, whereas *Wfs1* and *Prss35* were enriched in hippocampal CA1 and CA3 regions (Supplementary Fig. 48). We next examined whether the imputed genes also recovered gene-specific asymmetric intracellular organization beyond tissue-level anatomical patterns (Methods). Among 24 imputed genes with sufficient predicted abundance in VLMCs, five showed significantly greater polarity than count-matched backgrounds (all FDR < 1 × 10^−4^), and all five were also significantly polarized in the corresponding measured transcript maps from the original Xenium mouse brain dataset (all FDR < 0.05). Notably, three of these genes, *Spp1*, *Igfbp5*, and *Igfbp6*, encode secreted or extracellular-matrix-associated factors, consistent with the matrix-producing role of VLMCs. The predicted and measured polarity distributions of these genes and representative predicted *Spp1* localization in individual VLMCs are shown in Fig. 4i and Supplementary Figs. 49 and 50. We further extended the analysis from individual-gene polarity to pairwise subcellular organization (Methods). In astrocytes and endothelial cells, *Acsbg1*-*Slc39a12* and *Cldn5*-*Cobll1*, respectively, were the top-ranked co-localized pairs among the eligible imputed genes in the measured Xenium mouse brain data. These relationships were also biologically coherent: *Acsbg1* and *Slc39a12* are associated with astrocyte-mediated maintenance of the neural microenvironment, whereas *Cldn5* and *Cobll1* are associated with endothelial junctional and cytoskeletal organization. In the Xenium 5K imputation, *Acsbg1* showed significantly lower EMD to *Slc39a12* than to its count-matched background in astrocytes (FDR = 1.3 × 10^−8^^1^ ; Fig. 4j), while *Cldn5* showed significantly lower EMD to *Cobll1* in endothelial cells (FDR = 2.0 × 10^−6^). The same spatial proximity relationships were observed in the corresponding measured transcript maps (astrocyte FDR = 2.0 × 10^−1^^62^, endothelial FDR = 3.1 × 10^−1^^47^ ; Supplementary Fig. 51). A within-cell spatial-shuffle control further confirmed that these recovered gene-pair relationships reflected subcellular spatial organization beyond cell-level co-expression (Supplementary Note 11; Supplementary Fig. 52).

We further assessed the benefits of subcellular imputation in a disease context using matched WT and P301S tauopathy mouse brain datasets profiled with the Xenium Prime 5K panel. Focusing on Micro-PVM cells and astrocytes (Supplementary Fig. 53), SVC predicted significant tauopathy-associated localization shifts for three imputed genes (Methods). Specifically, in Micro-PVM cells, the predicted localizations of *Laptm5* and *Kctd12* shifted away from the nucleus in P301S samples, with median increases in nuclear distance of 0.0131 (FDR = 4.1 × 10^−4^) and 0.0176 (FDR = 4.0 × 10^−3^), respectively. In astrocytes, *Clmn* showed a predicted shift toward the nuclear region with a median decrease of 0.0098 (FDR = 4.6 × 10^−4^). Fig. 4k shows the corresponding predicted changes and aggregated subcellular distributions for *Laptm5* and *Clmn*; across genes, localization changes were only weakly correlated with predicted cell-level expression changes (Pearson correlation = 0.19; Supplementary Fig. 54). The predicted outward redistribution of *Laptm5* provides a clear, biologically interpretable illustration of a disease-associated RNA localization shift identified by SVC. P301S tauopathy is accompanied by pronounced microglial activation and lysosomal remodeling^41^, and LAPTM5 protein shows predominantly perinuclear localization in cultured microglia but a broader cytoplasmic distribution during neuronal-death-associated microglial activation^42^. SVC therefore extends this established inflammatory biology by predicting a previously uncharacterized RNA-level spatial response in tauopathy. In astrocytes, *Clmn* provides a complementary example. *Clmn* is a mature astrocyte-associated gene^43^, while CLMN has recently been identified as an ER-actin tether involved in the spatial organization of ER-plasma membrane contacts and the actin cytoskeleton^44^. Given that P301S tauopathy suppresses neuron-regulated astrocytic gene programs associated with astrocyte maturation and function^45^, the predicted nuclear-proximal redistribution of *Clmn* suggests an additional spatial dimension of astrocyte remodeling in tauopathy. More broadly, these results demonstrate that SVC can use subcellular imputation to uncover disease-associated localization changes in unmeasured transcripts, generating biological hypotheses beyond cell-level expression measurements.

### Xenium human breast cancer data

Finally, we evaluated SVC on the 10x Xenium human breast cancer dataset^6^ to demonstrate its applicability to tumor sections with pronounced spatial heterogeneity and microenvironmental gradients. Trained exclusively on tissue section 1, SVC identified highly consistent spatial domains in the overlapping region of the unseen section 2, demonstrating its ability to recover consistent spatial organization across sections (Fig. 5a, b). The identified domains corresponded closely to manually annotated tumor regions, including invasive ductal carcinoma (IDC; domains 1 and 14), ductal carcinoma in situ (DCIS) #1 (domains 3 and 16), and DCIS #2 (domains 5 and 15). Their biological identities were further supported by established markers (Fig. 5c): the invasive-tumor marker *FASN* was enriched in IDC-associated domain 1; the DCIS marker *CEACAM6* in domains 3 and 15; the myoepithelial marker *MYLK* in tumor-peripheral domain 5 and tumor-boundary domain 16; the stromal marker *LUM* in domains 0, 4, and 6; and the T-cell marker *CD8A* in immune-related domain 9. Complementary benchmarks and region-level out-of-distribution (OOD) analysis further showed that SVC maintained strong predictive performance for both subcellular transcript localization and cell-level gene expression while recovering coherent spatial domains in anatomically held-out regions (Supplementary Notes 9 and 13; Supplementary Figs. 55-60). SVC also retained predictive performance for cell-type-specific OOD localization patterns that differed substantially from those represented in the training cell types (Supplementary Note 14; Supplementary Figs. 61-63). Together, these results demonstrate the robustness and generalizability of SVC in capturing biologically coherent spatial organization at subcellular, cellular, and tissue-domain scales.

**Fig. 5.**
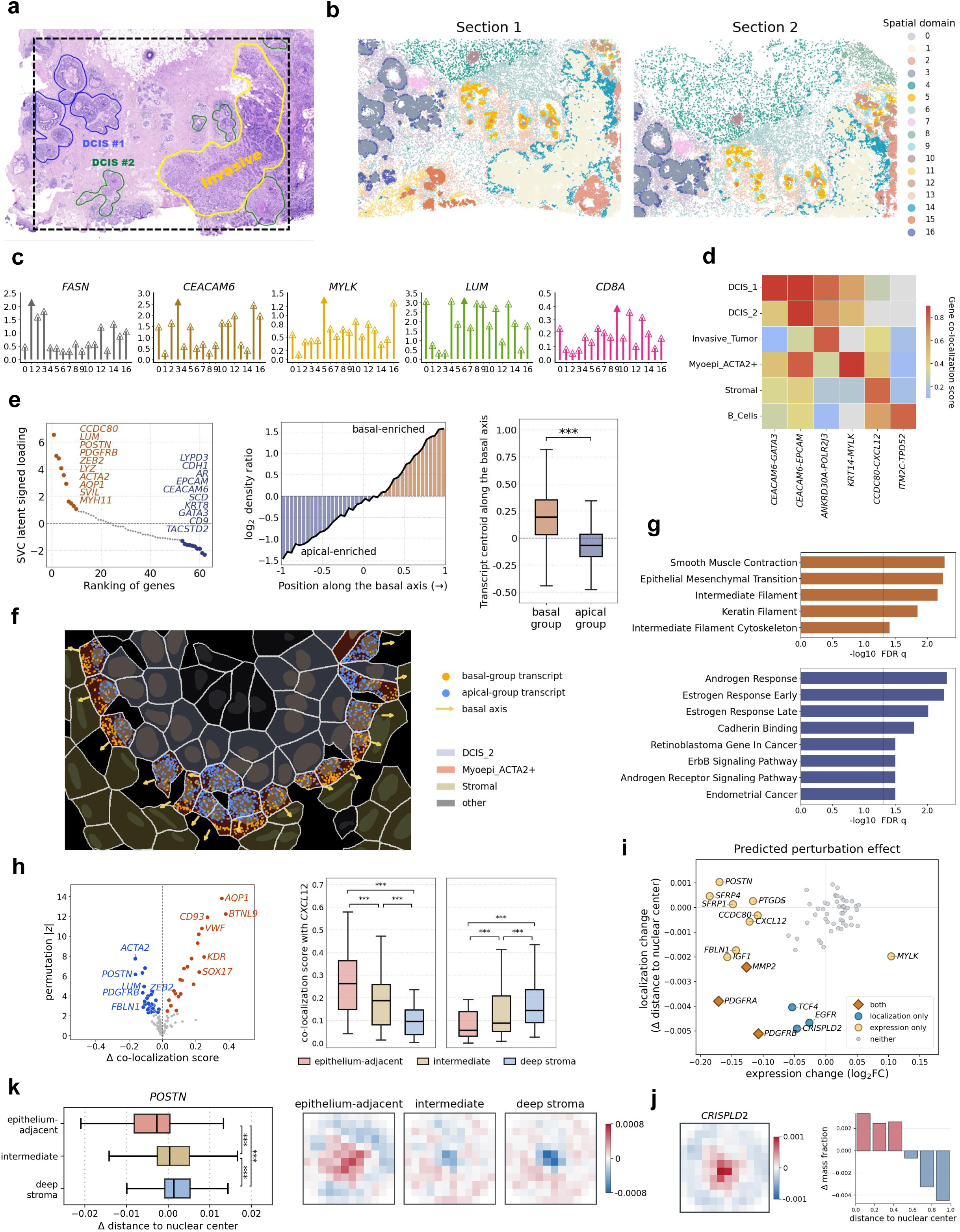
Xenium human breast cancer data analysis. **a.** Manually annotated tumor regions in tissue section 1 of the Xenium human breast cancer dataset. **b.** Spatial domains detected by SVC in both sections. **c.** Lollipop plots display the mean expression (log1p-transformed) of five marker genes (*y* axis) in each spatial domain (*x* axis). The solid arrow indicates the highest value. **d.** Heatmap visualizes gene co-localization scores of cell-type-specific gene pairs (columns) across six cell types (rows). For each cell type, the pair shown is the one whose score most exceeds its average across cell types. Grey indicates pairs with an insufficiently detected gene in that cell type. **e.** Left panel: signed loading of each gene on the basal-apical axis of ACTA2+ myoepithelial cells; the ten most basal (red) and ten most apical (blue) genes are labeled. Middle panel: transcript positions were projected onto the anatomy-derived axis from −1 (apical) to 1 (basal). The curve shows the log2 ratio between the transcript densities of the two gene groups along the axis. Right panel: box plots show the per-cell transcript centroid of each group along the same axis. Statistical significance (n.s. ≥ 0.05; * < 0.05; ** < 0.01; *** < 0.001) was assessed using a two-sided Wilcoxon signed-rank test. **f.** Representative ROI from section 1 illustrates observed transcript positions of the two ten-gene groups; arrows mark the basal axis of each myoepithelial cell. **g.** Bar plots show the terms enriched in the genes with positive (basal, 24 genes) and negative (apical, 38 genes) loading. Enrichment was computed with Enrichr^71^, querying GO (biological process, cellular component and molecular function), MSigDB Hallmark, Reactome and WikiPathways. *P* values were adjusted separately within each gene-set library using the Benjamini-Hochberg procedure; up to eight terms are shown per group. The x-axis is −log10 of the adjusted *P* value. **h.** Left panel: volcano plot of the change in co-localization with *CXCL12* in endothelial cells between epithelium-adjacent and deep-stromal regions, across 132 partner genes. The x-axis shows the difference in co-localization score and the y-axis shows the absolute z score from 2,000 group-label permutations; point color indicates genes with significantly stronger co-localization with *CXCL12* in epithelium-adjacent cells (22) or in deep stroma (29) at FDR < 0.05. Right panel: box plots show the co-localization scores of these two partner-gene sets across the three spatial contexts. Statistical significance for pairwise comparisons (n.s. ≥ 0.05; * < 0.05; ** < 0.01; *** < 0.001) was assessed using two-sided Wilcoxon signed-rank tests across genes. **i.** Scatter plot compares the cell-level and the subcellular response of 65 genes to an *in silico* depletion of *LUM* in stromal cells. The x-axis shows the log2 fold change of predicted counts and the y-axis the change in relative distance to the nuclear center (inward, negative; outward, positive); point color and shape indicate which of the two readouts is significant at FDR < 0.05. **j.** Left panel: change in the SVC-predicted subcellular distribution of *CRISPLD2* upon *LUM* depletion, averaged over cells and shown as the per-pixel change in mass fraction on the 12 × 12 image; each cell’s map was normalized to sum to one before averaging. Right panel: the same change summed within six concentric rings from the nuclear center to the cell boundary. **k.** Left panel: box plots show the per-cell change in the relative distance of *POSTN* transcripts to the nuclear center upon *LUM* depletion, for stromal cells located in epithelium-adjacent, intermediate and deep-stromal contexts. Right panel: the corresponding change in the predicted distribution of *POSTN*, computed as in **j** and averaged within each context. Statistical significance for pairwise comparisons (n.s. ≥ 0.05; * < 0.05; ** < 0.01; *** < 0.001) was assessed using two-sided Mann-Whitney U tests.

Given the pronounced cellular heterogeneity of breast tumors, we next examined cell-type-specific subcellular gene co-localization across six major cell populations and identified many co-localized pairs (Fig. 5d). For example, *KRT14*-*MYLK* showed preferential co-localization in ACTA2+ myoepithelial cells, potentially reflecting coordinated subcellular organization of the basal epithelial program and actomyosin contractile machinery that underlies myoepithelial contractile function^46, 47^. *CCDC80*-*CXCL12* preferential co-localization in stromal cells may reflect coordinated subcellular organization of ECM-associated and paracrine chemokine programs, consistent with the secreted ECM-associated function of *CCDC80* and fibroblast-derived *CXCL12* signaling in breast tumors^48, 49^. Complementary within-cell shuffle and co-expression analyses further showed that these co-localization relationships captured subcellular spatial organization beyond cell-level co-expression (Supplementary Figs. 64 and 65).

Apicobasal polarity is a fundamental organizing principle of epithelial tissues, coordinating directional cell-cell and cell-matrix interactions^50^. Yet how intracellular RNAs are coordinately organized to support this polarity remains largely uncharacterized, particularly within myoepithelial cells at DCIS boundaries. We therefore asked whether SVC could uncover continuous, polarity-aligned intracellular organization. Specifically, we analyzed 4,678 ACTA2+ myoepithelial cells, which were preferentially distributed along the periphery of DCIS epithelial regions at epithelial-stromal interfaces (Supplementary Fig. 66). Using local tissue geometry, we defined a cell-specific anatomical reference direction independently of SVC pointing outward from the densely cellular region toward surrounding lower-density tissue, consistent with the expected apicobasal orientation of myoepithelial cells (Methods; Supplementary Fig. 67). We then examined whether the gene-specific SVC representations varied coherently along this anatomical direction. After removing shared cell-level variation, joint modeling of gene-specific SVC representations with the anatomical axis identified a dominant shared directional component and a signed loading for each gene (Methods). This component explained 62.8% of the variation in gene-specific directional effects, far exceeding the 8.5-12.8% observed across 100 random-rotation controls (empirical *P* value = 0.0099; Supplementary Fig. 68), indicating that the SVC-derived directional signal was specifically aligned with local tissue geometry. Ranking genes by their signed loadings ordered them continuously along the shared directional component, with the top 10 genes at each end highlighted in Fig. 5e (left panel). Measured transcript distributions of the two loading-defined ten-gene groups showed preferential localization toward opposite ends of the anatomy-derived axis (Fig. 5e, middle panel), defining an outward-enriched basal group and an oppositely enriched apical group; this separation was also evident within individual cells, with the basal-group centroid positioned farther along the basal direction than the apical-group centroid (median 0.196 vs −0.066; paired Wilcoxon signed-rank test, *P* value < 1 × 10^−300^ ; Fig. 5e, right panel). *In situ* visualizations further showed opposite localization of the two gene groups along the anatomy-derived basal axis within individual myoepithelial cells (Fig. 5f). Functional enrichment analysis for genes with positive and negative loadings (Fig. 5g) showed that basal-associated genes were enriched for contractile and cytoskeletal programs, including smooth muscle contraction, intermediate filaments, keratin filaments, and epithelial-mesenchymal transition. Apical-associated genes were enriched for hormone-response, epithelial-adhesion, and signaling programs, including androgen and estrogen responses, cadherin binding, and ErbB signaling. Overall, SVC uncovered a coordinated, continuous, multi-gene apicobasal RNA localization axis within ACTA2+ myoepithelial cells at DCIS boundaries, revealing directional partitioning of distinct functional programs.

SVC also helped uncover context-dependent subcellular co-localization variation across tumor microenvironments. Here, we focused on partners of the well-studied endothelial chemokine *CXCL12*^51^ in endothelial cells across epithelium-adjacent, intermediate, and deep-stromal contexts using transcript-abundance-matched groups for each candidate partner (Methods). Among 132 evaluable genes, 51 showed significantly different co-localization with *CXCL12* between epithelium-adjacent and deep-stromal contexts (FDR < 0.05), including 22 with stronger co-localization in epithelium-adjacent cells and 29 with stronger co-localization in deep stroma (Fig. 5h; left panel). The partners with stronger *CXCL12* co-localization in epithelium-adjacent cells showed a progressive decrease in co-localization score from the epithelium-adjacent group to the intermediate and deep-stromal groups, whereas the deep-stroma-enriched partners showed the opposite trend. For both partner sets, gene-level co-localization scores differed significantly between every pair of spatial contexts (paired Wilcoxon test, all *P* values < 1 × 10^−4^ ; Fig. 5h; right panel). These results indicate a progressive, context-dependent shift in the subcellular co-localization partner profile of *CXCL12* along the epithelial-to-stromal axis.

Comparison of context-dependent changes in *CXCL12* co-localization with the corresponding changes in cell-level co-expression revealed only a moderate correlation (Pearson correlation = 0.47; Supplementary Fig. 69), indicating that the co-localization shifts reflect within-cell spatial organization beyond cell-level expression changes. Finally, functional enrichment analysis revealed distinct biological programs associated with the two context-specific partner sets (Supplementary Fig. 70). Partners preferentially co-localized with *CXCL12* near the epithelium were enriched for blood vessel morphogenesis, whereas deep-stromal partners were enriched for angiogenesis and epithelial-mesenchymal transition-related programs. Together, these results suggest context-dependent subcellular organization of endothelial *CXCL12* across microenvironments, consistent with its established roles in vascular development and remodeling^52, 53^.

Given that gene regulatory relationships in human breast cancer are highly complex and heterogeneous across distinct cell states and transcriptional programs^54^, we next conducted *in silico* perturbation experiments using SVC to characterize predicted effects of key gene depletions on cell-level expression and subcellular transcript organization within native tissue context (Methods). First, we conducted an *in silico* depletion of a canonical gene *LUM* in stromal cells across the tissue and systematically quantified the predicted downstream responses of all broadly expressed genes. Joint visualization of the predicted cell-level and subcellular effects revealed significant but only partially overlapping gene responses (Fig. 5i). Six genes showed significant subcellular localization changes, all characterized by shifts toward the nuclear center (Supplementary Fig. 71). Using *CRISPLD2* as a representative example, we visualized these predicted subcellular effects with aggregated spatial maps and radial profiles (Fig. 5j). Given that *CRISPLD2* encodes a secreted mesenchymal protein implicated in fibroblast regulation and inflammatory responses^55^, this inward redistribution may reflect spatial reorganization of secretory-protein biosynthesis following *LUM* depletion^32^. We next systematically examined context-dependent *in silico* perturbation effects by assessing, across all analyzed genes, whether the predicted subcellular perturbation effects varied among three stromal spatial contexts: epithelium-adjacent, intermediate, and deep stroma (Methods). Two genes showed significant context-dependent localization effects: *POSTN* (FDR < 1 × 10^−1^^5^), which showed the strongest difference, and *CCPG1* (FDR = 0.02). The predicted localization response of *POSTN* shifted progressively from inward in epithelium-adjacent stroma to outward in deep stroma, with intermediate stroma showing an intermediate effect (all *P* values < 0.001; Fig. 5k, left panel). Aggregated normalized maps and radial profiles illustrated the same context-dependent transition (Fig. 5k, right panel; Supplementary Fig. 72). This phenomenon may reflect spatial differences in stromal cell state and cell-matrix signaling. *LUM* depletion could therefore differentially reorganize the cytoskeletal-ER network across microenvironments^56^, favoring association of *POSTN* transcripts with perinuclear ER in epithelium-adjacent cells but their redistribution toward peripheral ER domains in deep-stromal cells^32^. We also conducted an *in silico* depletion of the *ERBB2* gene within tumor-related cells, revealing pronounced heterogeneity in downstream gene expression responses (Supplementary Note 15; Supplementary Figs. 73-75). Overall, SVC provides a new perspective for predicting spatial perturbation effects, offering valuable insights for the design of subsequent targeted perturbation experiments.

## Discussion

Here, we developed SVC, a ViT-based spatially and virtually embedded cell model for learning and predicting subcellular transcript organization within native tissue context. By integrating multi-scale and multi-modal information, including subcellular localization, gene-function priors, cell morphology, cell identity, and local tissue context, SVC learns unified representations across genes and cells that capture cell-specific spatial relationships among transcripts. The learned relationships support prediction of subcellular localization for unmeasured genes and *in silico* perturbation-associated changes at subcellular resolution, extending spatial transcriptomic analysis beyond characterization of measured transcripts. Across diverse datasets, SVC further revealed subcellular organization associated with cellular states, gene function, and local tissue microenvironments, facilitating our understanding of cellular function and behavior.

Despite these advances, SVC has several limitations that point to important directions for future development. First, SVC is currently trained on static datasets, reflecting the limitations of existing subcellular-resolution spatial transcriptomics technologies. Consequently, it is not yet capable of modeling dynamic responses over time in distinct environments at the single-cell level. Dynamic modeling would allow for time-resolved prediction of transcript localization changes in response to environmental or genetic stimuli, which is essential for capturing cell behavior in more realistic, physiologically relevant contexts. Second, we currently focus on targeted gene panel platforms, where predesigned probes enrich for moderately to highly expressed genes to support reliable subcellular pattern learning or imputation. As high-throughput spatial transcriptomics technologies continue to improve in depth and quality, extending SVC to more comprehensive datasets will enable modeling of a broader range of genes. Third, the available subcellular spatial transcriptomics datasets are observational in nature, such that the variation learned by SVC primarily reflects endogenous biological processes rather than targeted perturbations. As spatially resolved perturbation assays, such as Perturb-FISH^57^ and Perturb-map^58^, continue to mature, integrating perturbation-based data will be essential to generate causal, within-cell transcript relocalization responses under defined external stimuli. At the moment, SVC can predict the effects of individual gene perturbations *in silico*, but broadening the perturbation types and integrating perturbation-specific training data in the future would enable a key capability envisioned for fully functional AI-powered virtual cells (AIVCs). Fourth, while SVC currently models two modalities, spatial gene expression and morphology, the emergence of spatial multi-omics presents future opportunities to expand its capabilities. Modalities such as spatial proteomics, spatial chromatin accessibility, and spatial metabolomics could eventually reach subcellular resolution, providing richer cellular context for improved model interpretability. Fortunately, SVC’s modular architecture is inherently flexible and can readily accommodate additional subcellular resolution spatial omics data as additional embeddings, making it well-suited for future multi-omics extensions when these data become available. Fifth, although our sensitivity analysis suggested that SVC is relatively tolerant to moderate differences in cell segmentation (Supplementary Note 16; Supplementary Figs. 76 and 77), inaccuracies in upstream cell segmentation and transcript assignment remain an important consideration. We expect that continued advances in membrane-marker imaging and cell-segmentation algorithms will reduce such upstream errors and improve the reliability of SVC and downstream analyses. Finally, subcellular ST data vary widely across tissue types, disease contexts, gene panel designs, and experimental noise profiles. Although SVC demonstrated cross-dataset transferability, jointly integrating heterogeneous datasets to learn shared and robust representations across platforms, conditions, and gene panels remains an important direction for future development.

## Methods

### Data input and preprocessing

SVC is designed to characterize gene spatial expression patterns at the subcellular level within individual cells, while preserving their tissue spatial context, in high-resolution spatial transcriptomics studies. The model adopts a multi-modal framework that integrates two complementary data modalities: (i) gene expression profiles derived from subcellular ST data and (ii) cell and nuclear morphological features extracted from corresponding DAPI microscopy or H&E histology images. The subcellular ST data provide for each cell its gene expression and localization information, represented by the Cartesian coordinates (*x*,*y*) of all measured transcripts within the cell. The morphological images provide additional geometric context for each cell, including cell boundary information, defined by the coordinates of polygonal cell boundary points, the coordinates of the nuclear center, and pixel-wise masks that enumerate all coordinates belonging to the cell and its nucleus.

With this input information, we begin data preprocessing using a robust cell registration procedure based on ELLA^14^ to establish a unified cellular coordinate system, enabling spatial expression patterns shared across cells. Specifically, for each transcript within a cell, we compute two spatial parameters in polar coordinates: angle (*θ*) and relative position (*ρ*). The angle *θ* of a transcript is measured relative to the positive x-axis, with the nuclear center as the origin. The relative position *ρ* of a transcript is calculated as *ρ* = *l*_1_/(*l*_1_ + *l*_2_), where *l*_1_ is its distance to the nuclear center and *l*_2_ is its distance to the cell boundary in the opposite direction from the nuclear center. Using coordinate transformations that preserve *θ* and *ρ*, we project all transcripts within a cell onto a unit circle, providing a normalized coordinate system centered at the nuclear origin. After registration, we represent the subcellular spatial expression of each cell as a 48×48-pixel square image, with transcript counts recorded at the pixel level. Additionally, as complementary morphological inputs, for each cell, we constructed two 48×48-pixel binary masks for the cell body and nucleus (1 = structure, 0 = background). Both masks were centered at the nuclear centroid, with relative morphological sizes preserved across cells.

### Vision Transformer

After data preprocessing, we develop a Vision Transformer (ViT)^59^ framework to model subcellular gene spatial expression patterns within individual cells, while capturing cellular heterogeneity and the broader spatial tissue context. Unlike convolutional neural networks (CNNs), which rely on local receptive fields and hierarchical feature extraction, the ViT architecture is capable of leveraging self-attention mechanisms and transformer architecture to capture long-range dependencies in spatial expression patterns across genes. This capability makes ViT particularly well suited for modeling the complex subcellular spatial relationships among genes revealed by high-resolution spatial transcriptomics data. To fully harness the power of ViT, we introduce five embeddings that comprehensively capture distinct biological features at both the gene and cell levels, ensuring accurate and comprehensive characterization of subcellular spatial expression patterns. Below, we describe each of these embeddings in detail.

Gene-level subcellular localization embedding – We conceptualize the spatial expression pattern of each gene within a cell as analogous to an image patch. This analogy allows us to adapt ViT to encode the spatial expression pattern of each gene through localization embeddings, which serve as key inputs to the transformer architecture. Specifically, each cell is first divided into *N*_gene_ sub-images of size 48× 48, where *N*_gene_ denotes the number of genes. This structured representation not only allows explicit modeling of subcellular spatial localization patterns for individual genes, but also facilitates the learning of cross-gene spatial relationships within the ViT framework. Due to the inherently low read counts in common subcellular-resolution spatial transcriptomics technologies, these high-resolution images often contain sparsely distributed gene expression counts across pixels. This sparsity presents computational challenges when learning effective feature representations from spatial transcriptomics data. To address this, we apply convolution to locally aggregate expression counts per pixel, thereby generating a lower-resolution transcript map for each cell while preserving essential spatial expression features. Specifically, we conduct a resolution reduction operation to convert each high-resolution image to a 12×12 low-resolution image via a 4×4 sum pooling kernel with stride 4. This resolution reduction operation not only alleviates the computational challenge posed by data sparsity but also enhances robustness to the noise nature of subcellular transcript localization. By doing so, it enables our model to focus on learning the overall spatial distribution patterns of transcripts rather than enforcing precise coordinate prediction, helping mitigate overfitting while maintaining biological interpretability. We denote the observed expression count data at pixel (*h*,*w*) in the 12 × 12 image for cell *i* and gene *j* as *y_i_*_,*j*,ℎ,*w*_ . For library size normalization, we used a cell-specific size factor *s_i_*, normalized relative to the median total transcript count across cells, to account for differences in total transcript abundance. We then used *s_i_* to normalize the observed counts *y_i_*_,*j*,ℎ,*w*_, correcting for variability in mRNA capture efficiency across cells. Afterwards, we flattened each image per gene within each cell into a 144-dimensional vector. This vector is then projected into a *d*-dimensional (default 384) latent vector *g_i_*_,*j*_, for *i*-th cell and *j*-th gene, using a trainable linear transformation, with the weights learned from the data. This projection into the *d*-dimensional latent space ensures that a consistent latent vector size is provided to the transformer architecture across all its layers.

Gene-level functional embedding – In parallel, we extract semantic embeddings of genes based on transcriptome-wide co-expression relationships, to provide gene-level functional embeddings to our ViT model. These embeddings are derived from the Gene2vec^60^ model, which integrates external knowledge about gene function and encodes prior information on biological relationships among genes, thereby enhancing both the biological interpretability and effectiveness of our model. The use of the gene-level embeddings also addresses a key limitation of conventional ViT architectures when applied directly to gene expression data. Specifically, in standard ViT models, the order of image patches is encoded through positional embeddings, enabling the model to leverage spatial proximity and semantic relationships among patches. In contrast, in gene expression modeling, shuffling the order of genes should not alter their biological meaning, rendering traditional positional encoding inappropriate. To overcome this, we replace positional embeddings with gene-level functional embeddings, which capture functional relevance between genes. By incorporating these embeddings, our model integrates broad functional relationships among genes, effectively complementing the spatial information encoded in the gene-level localization embeddings. In the Gene2vec model, each gene is represented by a 200-dimensional vector learned from bulk RNA-seq data; these vectors are commonly employed as key inputs to single-cell large language models (LLMs) such as scBERT^61^ and GeneCompass^62^. To enforce consistent vector dimensionality across transformer layers, we map each pre-defined 200-dimensional Gene2vec vector into a *d*-dimensional latent vector *p_j_* for *j*-th gene using a trainable linear transformation, with the weights learned from the data.

Cell-level neighbor embedding – To capture local spatial context, we characterize the microenvironment surrounding each cell and generate cell-level neighbor embeddings. These embeddings incorporate information from neighboring cells to represent the local tissue context, enabling the model to account for the cellular microenvironment when modeling subcellular gene spatial expression within each cell. Additionally, modeling subcellular spatial expression patterns within the broader tissue context enables the capture of biological processes across scales, encoding a potential hierarchical structure of gene expression that links microenvironmental context to subcellular organization. To set up the embedding, for the *i*-th cell we identify its *k* nearest cells by Euclidean distance between nuclear centers, and represent each neighbor by its normalized, log-transformed cell-level gene expression profile. Neighbors are then pooled by a distance-weighted average in which a neighbor’s contribution decays exponentially with its distance to the focal cell, with the decay set by a temperature equal to the median distance to the *k*-th nearest neighbor across cells. To fit inputs of the transformer architecture, we further map the pooled neighbor profile of the *i*-th cell into a *d*-dimensional latent vector *n_i_*.

Cell-level morphological embedding – Subcellular-resolution ST data are often accompanied by parallel morphological images (e.g., DAPI-stained fluorescence or H&E histology images), which provide valuable cell boundary and morphological information highly complementary to spatial gene expression data. Jointly modeling both these modalities facilitates integrative analyses that capture the multi-modal nature of cellular processes and behaviors. To leverage such multi-modal features of the data, we provide each cell with additional embeddings extracted from binary morphological images. These embeddings encode cell-specific geometric and structural features of individual cells, capture continuous variation in cellular phenotypes, and offer complementary cell and nuclear morphology information to spatial transcriptomics data. For each cell *i*, the 48×48 cell and nuclear morphology images are each divided into non-overlapping 8 × 8 patches; each patch is then projected by a trainable linear transformation, and the resulting patch embeddings are averaged into two d-dimensional latent vectors, *m_i_*^cell^ and *m_i_*^nuclear^, respectively.

Cell-level identity embeddings – Finally, when additional cell identity information, such as cell type or cell state labels, is available, it can be leveraged to construct cell-level identity embeddings. These embeddings encode categorical information that distinguishes different cellular populations, allowing the model to explicitly account for cellular heterogeneity. By integrating cell identity features, our model can more accurately capture cell-identity-specific gene expression patterns and variations that arise from distinct functional states or differentiation stages of cells. These embeddings complement the spatial and morphological embeddings above, providing an additional layer of biological context that enhances interpretability and predictive performance. Here, each cell label is first encoded as a one-hot vector and then mapped into a shared d-dimensional latent space via a trainable linear transformation with the weights learned from the data, resulting in a cell identity embedding vector *f_i_* for *i*-th cell.

These five complementary embeddings enable the ViT architecture to perform comprehensive feature extraction in a multi-modal and multi-scale manner, learning the contextual dependencies and spatial semantic associations across genes, thereby enhancing the model’s ability to capture complex spatial expression patterns of genes within cells.

### Subcellular contextual learning via masked image modeling

We employ a self-supervised Masked Image Modeling^63^ strategy to learn subcellular spatial patterns of gene expression without requiring any pixel-level labels or task-specific annotations. The key idea is to randomly mask a subset of gene expression images in each cell during training and task the model with reconstructing the masked content based on the remaining unmasked information. This strategy facilitates subcellular contextual learning, in which the subcellular pattern of a target gene is learned in the context of the remaining genes within the same cell, enabling effective gene-specific subcellular inference while capturing complex cross-gene spatial dependencies.

Specifically, during each training iteration, we randomly select 10% of the genes in each cell and mask their corresponding spatial expression images. For each cell *i*, we represent its spatial gene expression pattern as a subcellular localization embedding matrix *G_i_*^mask^ of size *N*_gene_×*d*, where *N*_gene_ denotes the total number of genes and *d* is the embedding dimension described in the previous section. The matrix *G*^mask^ is identical to the gene-level localization embeddings *G_i_* = (*g_i_*_,1_ *g_i_*_,2_ . . . *g_i_*_,*N*_)*^T^* with *g_i_*_,*j*_ defined earlier, except that the rows corresponding to masked genes are replaced with a learnable mask token. For the selected genes, only the subcellular localization embeddings are replaced with mask tokens, while the gene functional, morphology, and cell-identity information is treated as observed conditioning information that helps the model infer the masked localization patterns. To prevent information leakage through the neighborhood input, the neighbor embedding *n*^mask^ is computed as *n_i_* defined earlier, except that the expression profile of each neighbor is restricted to the unmasked genes of cell *i*, so that masked genes remain unobserved in the neighbor context as well.

We then train our ViT model to reconstruct the full spatial expression of the masked genes by leveraging information from the unmasked genes as well as auxiliary contextual inputs. During this process, SVC integrates comprehensive information from all five embeddings to learn cell- and gene-specific representations of subcellular transcript organization. Specifically, we denote the gene-level functional embeddings *P_i_*, cell-level neighbor embeddings *n_i_*^mask^, cell-level morphological embeddings *m_i_*^cell^ and *m_i_*^nuclear^, and cell-level identity embeddings *f_i_* (when available). To explicitly model the shared effect of cell-level information across genes, we introduce feature-wise linear modulation (FiLM)^64^. Specifically, each cell-level embedding is mapped by a two-layer perceptron to a scale and a shift vector of dimension *d*; these are summed across the cell-level embeddings, each weighted by its own learnable gate, yielding a single pair (*γ_i_*, *β_i_*) per cell that is applied to every row of *G_i_*^mask^ as 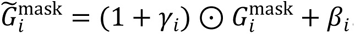. We employ a Performer encoder^65^ **F**, a scalable Transformer architecture based on linear attention,

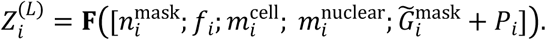

The semicolon in the equation denotes vector concatenation by row. Since the Performer encoder consists of *L* successive blocks, we denote the representation after the *l*-th block as 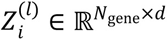, with the final output given by *Z_i_^(L)^*, where each row corresponds to one gene in cell i . The cell-level tokens serve as conditioning inputs to the encoder and, for notational simplicity, are not included in *Z_i_^(l)^*, which contains only the gene-level representations. We add *Ḡ_i_*^mask^ and *P_i_* following the standard ViT embedding formulation^59^. These two embeddings encode complementary information for the same gene token. As previous work has shown comparable performance for additive and concatenative embedding fusion^66^, we use addition because it preserves the token dimensionality without requiring an additional projection layer. Importantly, the Performer encoder captures long-range gene-gene dependency through a multi-head self-attention mechanism while preserving the token embedding dimension. Compared with standard Transformer encoders, whose self-attention scales quadratically with sequence length, Performer’s linear-attention mechanism also empowers scalable processing of thousands of genes simultaneously for spatial transcriptomic analyses.

The inferred final-layer representation *Z_i_^(L)^* is then used to predict the spatial gene expression patterns of masked genes in the training data. To account for the overdispersed nature of count data, for gene *j* in cell *i*, we model *y_i_*_,*j*,ℎ,*w*_ using a negative binomial (NB) distribution, parameterized by a mean *μ_i_*_,*j*,ℎ,*w*_ and an inverse-dispersion *φ_i_*_,*j*,ℎ,*w*_, where *y_i_*_,*j*,ℎ,*w*_ denotes the observed count at pixel (*h*,*w*) of the 12 × 12 image defined earlier. To recover the spatial probabilistic patterns of masked transcripts, we use two lightweight one-layer decoders applied to *Z_i_^(L)^* to estimate the pixel-wise NB mean and inverse-dispersion parameters. Specifically, for a masked gene *j* in cell *i*, let *μ̄_i_*,*_j_*_,ℎ,*w*_ and *φ*^*_i_*,*_j_*_,ℎ,_*_w_* denote the outputs of the NB mean and inverse-dispersion decoders at pixel (*h*,*w*), respectively. We calculate the negative log-likelihood of the NB distribution and obtain the pixel-level loss function in the form of:

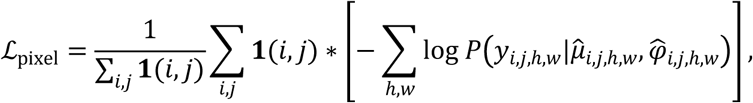

where **1**(*i*, *j*) = 1 if gene *j* is masked in cell *i*, otherwise 0. Here, the summation over (*h*,*w*) is taken only over foreground grid locations within the registered unit circle, corresponding to pixels inside the cell mask. To fit the distribution parameters of raw count *y_i_*_,*j*,ℎ,*w*_, *μ̄_i_*,*_j_*_,ℎ,_*_w_* is multiplied by the corresponding cell-specific size factor *s_i_* to obtain the estimated mean parameter *μ*^*_i_*,*_j_*_,ℎ,_*_w_*, accounting for library-size differences during optimization while retaining the original count scale of the data. For masked-gene training and prediction, *s_i_* was estimated from the unmasked genes, with their total transcript count adjusted for the proportion of genes remaining observed. To explicitly capture the total abundance of a gene within a cell, we add a cell-level expression loss. For each masked gene, the predicted and observed counts are summed over the foreground pixels and compared on a log scale with a smooth L1 loss:

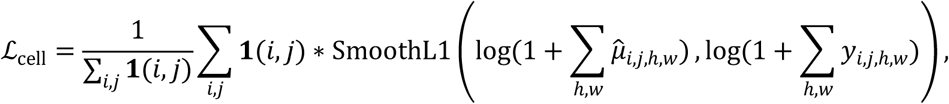

where SmoothL1(·) denotes the Huber loss with a transition point of 1. The final training objective combines the two terms:

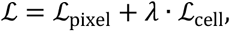

where λ is a hyperparameter balancing the two losses.

### Output and postprocessing

Our model generates two core outputs: (1) a unified latent representation for individual genes and cells and (2) predicted spatial gene expression for new genes in the test data.

For downstream representation-based analyses, we define the latent representation as the average of the representations from the first three encoder blocks, 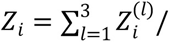. We use representations from the early encoder blocks because lower Transformer layers tend to preserve more general features, whereas deeper layers become increasingly specialized to the training objective^67^. The resulting representation is both cell-specific and gene-specific, fusing rich multi-scale, multi-modal information learned from the data. Our transformer architecture accommodates the intrinsic complexity of cellular processes and captures nonlinear subcellular spatial variation of gene expression. This integrated representation facilitates the inference of complex multi-gene interactions and relationships and uncovers spatial organization patterns that underlie the fundamental molecular logic of cells. Consequently, we can explore gene-specific subcellular localization patterns and also characterize cellular heterogeneity by quantifying within-cell localization characteristics across genes.

On the other hand, our trained ViT model can also be directly applied to predict subcellular spatial expression patterns for genes that are absent in the test data but present in the training data. Notably, the model generates such maps for unmeasured genes in the test data and estimates the corresponding 12×12 NB mean parameters for each cell, thereby producing gene-specific predicted transcriptomic maps. These 12× 12 predicted expression maps serve as the primary input for most downstream analyses. Additionally, the 12×12 maps can be probabilistically converted back to the original 48 × 48 resolution within each cell’s morphology, enabling visualization within the native cell geometry at higher spatial resolution. To reconstruct the 48×48 high-resolution spatial map, we partition each pixel in the 12×12 low-resolution map into sixteen 4×4 subregions. Let *μ*^*_i_*,*_j_*_,ℎ,_*_w_* and *φ*^*_i_*,*_j_*_,ℎ,*w*_ denote the estimated NB mean and inverse-dispersion parameters at pixel (*h*,*w*) for cell *i* and gene *j*. Assuming additivity of counts across subpixels, we model the expression count at each subpixel of (*h*,*w*) as an independent and identically distributed NB variable with mean *μ*^*_i_*,*_j_*_,ℎ,_*_w_*/16 and inverse-dispersion *φ*^*_i_*,*_j_*_,ℎ,_*_w_*/16 . Using these parameters, we perform probabilistic sampling at the subpixel level to generate a 48 × 48 high-resolution expression count map, thereby preserving the distributional characteristics of the original 12×12 representation.

To preserve the spatial expression patterns within the actual cell shape, we map the predicted high-resolution spatial expression from the unified cellular coordinate system back to the original cell morphology. To achieve this, for each predicted transcript in the 48 × 48 spatial map, we compute two spatial parameters using the polar coordinate system: the relative radial distance *ρ*’ ∈ [0,1] and the angular position *θ*’ ∈ [− 180, 180] . Let the nuclear center in Cartesian coordinates be denoted as *c*_0_ = (x_0_, y_0_) . We identify the cellular boundary point corresponding to angle *θ*’ and denote its Cartesian coordinates as *c*’ = (*x*’, y’), such that *θ*’ represents the angle of *c*’ relative to *c*_0_. This way, we determine the Cartesian coordinates (*x*,*y*) of the transcript within the original cell as follows:

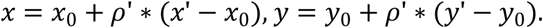

We refer to our method as SVC, which is implemented as a Python package. The SVC software and model implementation are freely available at https://github.com/aster-ww/SVC.

### Downstream analysis

### K-fold masked gene prediction task

To evaluate the model’s ability to predict the expression patterns of unobserved genes for benchmarking analysis, we designed a K-fold masked gene prediction task. In this case, the training and test datasets shared the same set of genes. We first trained the model on the full gene set. During testing, the genes were randomly partitioned into K groups. In each fold, one group was masked from the test data, and the trained model was directly applied to predict the spatial gene expression map of the masked genes based on the remaining observed genes. This process was repeated across all the groups, resulting in predictions for the entire gene set. Performance metrics were then aggregated across folds to provide a comprehensive evaluation of the model’s capability to infer unmeasured gene expression patterns. For the seqFISH+ mouse embryonic fibroblast dataset, K was set to 20; for the MERFISH U2-OS dataset and the two Xenium tissue datasets, K was set to 10.

### Subcellular prediction baselines and evaluation metrics

Since SVC is the first method to predict subcellular spatial gene expression, there are no existing methods available for direct comparison. To benchmark its performance, we constructed four baseline models. *Baseline 1* predicts each target cell’s gene expression image using a randomly selected image of the same gene from a training cell, thereby preserving gene-specific global patterns while disrupting cell-specific spatial organization. *Baseline 2* randomly permutes the pixel locations of the corresponding ground-truth subcellular image for each target cell-gene pair, preserving the per-cell, per-gene pixel-value distribution and total transcript count while disrupting its subcellular spatial organization. *Baseline 3* predicts each gene using its mean subcellular image across all training cells. *Baseline 4* predicts each target cell-gene pair using the mean ground-truth subcellular image of the same gene across the remaining test cells, with the target cell excluded from the average to avoid information leakage. Note that Baseline 4 is not feasible in practical imputation settings where the target gene is not measured in the test cells. For datasets containing multiple cell types, *Baselines 1*, *3*, and *4* were constructed within each cell type, such that the randomly selected training cell in *Baseline 1* and the population-average images in *Baselines 3* and *4* were restricted to cells of the same cell type as the target cell.

We evaluated prediction performance using six complementary metrics: root mean squared error (RMSE), Pearson correlation coefficient (PCC), cosine similarity, Earth Mover’s Distance (EMD), Kullback-Leibler divergence (KL), and Jensen-Shannon divergence (JS). RMSE quantifies the average prediction-error magnitude, PCC evaluates the overall linear trend consistency, and cosine similarity measures the directional alignment of vectors. EMD, KL divergence, and JS divergence further evaluate similarity at the spatial-distribution level. EMD measures the spatial displacement required to transform one transcript distribution into another, KL divergence quantifies the discrepancy of the predicted distribution from the ground truth distribution, and JS divergence provides a symmetric and bounded measure of distributional dissimilarity. For each gene in turn, we calculated these metrics between each predicted and ground-truth image pair and then averaged the values across cells. EMD, KL, and JS were calculated on unit-normalized predicted and ground-truth images with non-zero total mass. We ran the stochastic *Baselines 1* and *2* with 10 different random seeds and averaged the results for comparison. Lower values of RMSE, EMD, KL, and JS indicate better prediction quality, whereas higher values of PCC and cosine similarity indicate better prediction quality. For all metrics, performance gain was defined such that positive values indicate improvement over the corresponding baseline. For RMSE, EMD, KL, and JS, the relative gain of SVC over each baseline was calculated as the difference between the baseline score and SVC score divided by the baseline score. For PCC and cosine similarity, the gain was defined as SVC’s score minus the corresponding baseline score.

### Relative distance to the nuclear center

For each gene, we first computed the average relative subcellular position of each cell. This was done by calculating the weighted average distance of each pixel to the image center, with weights proportional to the gene’s expression level at that pixel. Because all images were registered within a unit circle, the resulting metric ranges from 0 to 1. Values approaching 0 indicate a gene’s preferential localization near the nuclear center, while values nearing 1 suggest enrichment near the cell boundary. Afterwards, we averaged these values across cells to derive each gene’s mean relative distance to the nuclear center.

### Concordance with subcellular localization of encoded protein

For the MERFISH U2-OS data analysis, since we lack well-annotated mRNA subcellular localization categories, we applied an alternative scheme to validate the accuracy of predicted patterns. Each cell image was segmented into three concentric zones — inner, middle, and outer — defined by increasing radial distance from the nuclear center and approximating the nucleus, cytoplasm, and cell membrane regions, respectively (see Supplementary Fig. 78 for illustration). For each gene, we quantified its enrichment within specific subregions by calculating the average predicted proportion of transcripts in each subregion. Genes exhibiting the highest enrichment in each subregion were taken as the most likely representatives of the corresponding subregion. In addition to comparing the top 10 subregion-specific genes with those obtained from the observed real images, we also cross-referenced the predicted mRNA locations with the subcellular localizations of the encoded proteins reported in the Human Protein Atlas (HPA, https://www.proteinatlas.org). A prediction was considered concordant with its encoded protein when at least one HPA annotation term matched the predicted compartment: nuclear terms for nuclear-enriched genes, plasma-membrane terms for membrane-enriched genes, and cytoplasmic or cytoplasmic-organelle terms for cytoplasm-enriched genes. We also recorded whether the supporting protein-localization evidence in HPA was provided from a U2-OS study (treated as strong evidence) or from other cell lines (treated as weak evidence).

### Subcellular gene co-localization analysis

The latent representation derived by SVC captures gene-specific subcellular patterns within individual cells. To quantify subcellular co-localization, we extracted the gene-specific latent representations from SVC and calculated pairwise cosine similarities within each cell. Specifically, we used *Z_i_* to compute cosine similarity scores between gene pairs within each cell, providing a quantitative measure of their co-localization. For each gene, we first identified its 10-nearest neighbors (10-NN), based on these similarity scores, within cells where it exhibited non-zero expression. Then, for each gene pair, we calculated the proportion of cells in which one gene appears in the 10-NN of the other, and vice versa. The gene co-localization score was defined as the larger value of these two proportions. Gene pairs with the highest co-localization scores were prioritized as top candidates exhibiting subcellular co-localization patterns. Note that the subcellular co-localization analysis captures spatial organization within cells and thus differs from conventional cell-level co-expression or correlation analyses, which reflect co-expression or co-variation across cells.

### Cell clustering

In the MERFISH U2-OS dataset, we used the SVC-derived latent representation to perform cell clustering characterized by consistent subcellular gene organization patterns. For each cell, we first computed a pairwise gene-gene cosine similarity matrix based on *Z_i_* . To quantify the similarity between each cell pair (*i*_1_,*i*_2_), we calculated the Euclidean distance *d_i_*_1,*i*2_ between their respective gene-gene similarity matrices. These cell-cell distances were then transformed into a similarity matrix using a Gaussian kernel 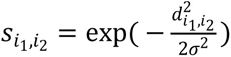, where the bandwidth parameter σ was set to the median of all pairwise distances. Finally, spectral clustering was applied to group cells exhibiting similar subcellular gene localization patterns.

### Spatial domain detection

Beyond cell clustering, the SVC-derived latent representation can also be leveraged to identify spatial domains across the tissue. As in the cell-clustering analysis, we calculated intercellular distances by measuring the Euclidean metric between the gene-gene cosine similarity matrices of each cell pair. To reduce computation burden, we then constructed a sparse latent cell-cell adjacency matrix *A*_latent_ using these distances, where an element is set to one if the corresponding cell pairs are mutual 10-nearest neighbors, and 0 otherwise. In parallel, we built a spatial cell-cell adjacency matrix *A*_spatial_ based on the Euclidean distances between cells’ actual spatial coordinates, again using mutual 10-nearest neighbors. The combined adjacency matrix *A*_latent_ + *A*_spatial_ was then used for spectral clustering to detect spatial domains. For cross-section analysis of the Xenium breast cancer data, SVC was trained on Section 1 and applied to both sections to obtain latent representations. Section 2 was first aligned to the training Section 1 using STalign^68^, after which domain detection was applied to the aligned spatial coordinates.

### Transcript-count filtering for localization-derived analyses

To reduce the influence of sparse cell-gene observations on localization-derived downstream analyses (Supplementary Note 3), we applied transcript-count-based quality-control criteria unless otherwise specified for individual analyses. For downstream analyses in which the results are interpreted in terms of subcellular localization or co-localization, including mean relative distance to the nuclear center and gene co-localization analysis, we retained only genes with at least 20 valid cells within the cell population or selected group of interest, where a valid cell was defined as having at least five transcripts for that gene. For subcellular localization analyses based on the mean relative distance to the nuclear center, cell-gene images with total subcellular transcript counts below five were further excluded.

### Gene-specific subcellular RNA organization analysis in VLMCs

To characterize fine-grained variation in subcellular RNA organization within a single cell type, we analyzed 6,613 vascular and leptomeningeal cells (VLMCs) from the Xenium mouse brain dataset. For each cell, we centered the 384-dimensional gene-specific representations across genes to remove the shared cell-level component. We retained genes with at least five detected transcripts in at least 100 cells across the two sections, resulting in 91 genes. For each gene, we then averaged its centered representations across VLMCs in which the gene had at least five detected transcripts. The median per-cell transcript count and total transcript count were calculated over the same cells, log-transformed, and regressed out of the gene-level representations, followed by principal component analysis (PCA) on the residuals.

To interpret the major axes identified by PCA, we quantified three subcellular localization characteristics for each gene across VLMCs: relative distance to the nuclear center, gyration, and polarity. For polarity and gyration, let *δ*_ℎ,*w*_ denote the normalized spatial offset vector of pixel (h, w) from the image center, with distances expressed relative to the grid half-width (6 pixels), and *π*_ℎ,*w*_ the fraction of transcripts of a given gene assigned to that pixel, with ∑_ℎ,*w*_ *π*_ℎ,*w*_ = 1 . Polarity was defined as the magnitude of the transcript-weighted mean offset, ∥ ∑_ℎ,*w*_ *π*_ℎ,*w*_ *δ*_ℎ,*w*_ ∥, corresponding to the displacement of the transcript centroid from the image center, reflecting asymmetric transcript localization within cells. Gyration was defined as the transcript-weighted root-mean-square distance from the transcript centroid, 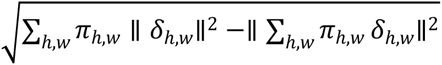, reflecting the spatial spread of transcripts around their centroid. These descriptors were calculated for each gene within each cell and averaged across VLMCs in which the gene had at least five detected transcripts.

For protein-targeting classification, genes were annotated using reviewed mouse Swiss-Prot entries from UniProt^30^. Genes were classified according to the protein-targeting route most relevant to transcript association with the endoplasmic reticulum (ER). Classification rules were applied hierarchically: genes encoding proteins with at least one annotated transmembrane segment were classified as transmembrane; among the remaining genes, those annotated as secreted, extracellular, extracellular-matrix associated, or GPI-anchored were classified as secretory/GPI-anchored; and among those still unassigned, genes without an annotated signal peptide were classified as intracellular. Of the 91 genes included in the VLMC analysis, 27 were classified as transmembrane, 22 as secretory/GPI-anchored, and 40 as intracellular. The remaining two genes lacked reviewed UniProt entries and were excluded from the protein-targeting analysis.

### Oligodendrocyte maturation-associated subcellular localization analysis

For the oligodendrocyte maturation analysis, we retained 1,101 oligodendrocytes in which all six genes from the OPC-associated and maturation-associated modules were detected. For each cell, pairwise cosine similarities were calculated between the gene-specific SVC representations of the six genes and then averaged across cells. The resulting pairwise similarities were further averaged separately for gene pairs within the OPC-associated module, within the maturation-associated module, and between the two modules. To quantify the relative radial positioning of the two gene modules, we pooled transcripts from all genes within each module and calculated their relative distance to the nuclear center. A relative positioning score was quantified as the distance of the maturation-associated module minus that of the OPC-associated module, such that positive values indicate a more peripheral position of the maturation-associated module relative to the OPC-associated module, whereas negative values indicate the opposite.

### Imputation analysis in the Xenium 5K mouse brain dataset

For the polarity analysis, we used the polarity metric defined in the preceding VLMC subcellular RNA organization analysis. We used a relaxed filtering threshold given the greater sparsity of the Xenium 5K dataset, retaining imputed genes with at least three predicted transcripts in at least 20 VLMCs, resulting in 24 eligible genes of the 84 imputed genes. For each target gene, we constructed a leave-one-out background from all other eligible imputed genes in VLMCs. To control for differences in transcript abundance, background gene-cell observations were subsampled within transcript-count bins, with two background observations drawn for each target observation, to match the target gene’s per-cell transcript-count distribution. The polarity distribution of each target gene was then compared with its count-matched background using a one-sided Mann-Whitney U test, followed by Benjamini-Hochberg correction across genes. To assess whether the localization patterns identified from the Xenium 5K imputation were consistent with independently measured data, we applied the same polarity analysis and count-matched background procedure to the corresponding measured transcript maps in VLMCs from the Xenium mouse brain dataset.

For the gene-pair spatial proximity analysis, we focused on astrocytes and endothelial cells and applied the same abundance filter used for the polarity analysis, retaining 20 and 8 eligible imputed genes, respectively. To define representative reference pairs, we ranked eligible gene pairs by their gene co-localization scores in the corresponding measured Xenium mouse brain data and selected the top-ranked pair in each cell type, *Acsbg1*-*Slc39a12* in astrocytes and *Cldn5*-*Cobll1* in endothelial cells. For each cell, the predicted subcellular distribution of each gene was normalized to sum to one, and pairwise co-localization was quantified using EMD, with lower values indicating stronger spatial overlap. For each target pair, a leave-one-out background was constructed by pairing the anchor gene with all other eligible imputed genes. Background observations were matched to the target pair using the same count-matching procedure, with transcript-count bins defined by the summed per-cell counts of the two genes and two background observations drawn for each target observation. The target-pair EMD distribution was compared with its count-matched background using a one-sided Mann-Whitney U test, followed by Benjamini-Hochberg correction across tested gene pairs. The same analysis was then applied to the corresponding measured transcript maps in the Xenium mouse brain dataset.

### Disease-associated subcellular imputation in P301S tauopathy

For the disease-associated imputation analysis, we applied the SVC model trained on the Xenium mouse brain dataset, without retraining, to matched WT and P301S tauopathy mouse brain sections^69^ profiled with the Xenium Prime 5K panel. We imputed 84 genes that were present in the training panel but absent from the measured target panels and focused on Micro-PVM cells and astrocytes. For each cell type, we retained genes with at least three predicted transcripts in at least 20 cells in both WT and P301S samples, resulting in 5 eligible genes in Micro-PVM cells and 11 in astrocytes. For each eligible gene, we calculated the predicted per-cell relative distance to the nuclear center and compared its distribution between WT and P301S cells using a two-sided Mann-Whitney U test, followed by Benjamini-Hochberg correction across genes within each cell type.

### Apicobasal RNA localization analysis in myoepithelial cells

We analyzed the 4,678 ACTA2+ myoepithelial cells from the two sections of the Xenium human breast cancer dataset. For each cell, the 384-dimensional gene-specific SVC representations were centered by subtracting their mean across genes, thereby removing the representation component shared across genes within that cell. An anatomical reference direction was derived independently of SVC. Nuclear centroids were rasterized onto a 3-µm grid, and the resulting nuclear-density field was smoothed using a Gaussian kernel with an 18-µm bandwidth. The negative local density gradient was normalized to unit length and used as the anatomy-derived basal reference direction, pointing from the densely cellular region toward the surrounding lower-density tissue. To improve the stability of the gene-specific directional estimates used in the shared decomposition, genes were retained if at least five transcripts were detected in at least 100 cells, yielding 62 genes. For each gene, its centered 384-dimensional representations were regressed on the two-dimensional basal reference vectors across eligible cells using ordinary least squares, with both variables centered across cells before fitting, yielding a 2 × 384 coefficient matrix per gene. These matrices were stacked across genes, column-centered, and factorized by singular value decomposition. Each gene’s score along the dominant shared directional component was defined as its loading on the first left singular vector multiplied by the corresponding singular value. To test whether this shared component reflects tissue geometry rather than arbitrary structure in the representations, the same regression and decomposition procedure was repeated on 100 null datasets in which each cell’s reference vector was independently rotated, and the variance explained by the first component was compared with the resulting null distribution (Supplementary Fig. 68).

### Context-dependent analysis of *CXCL12* co-localization partners

To examine whether the subcellular co-localization partners of *CXCL12* varied across tissue microenvironments, we focused on endothelial cells located in nine major SVC-derived spatial domains spanning the epithelial-to-stromal axis. Domains 1, 5, and 16 were designated as epithelium-adjacent, domains 0, 9, 13, and 14 as intermediate, and domains 4 and 6 as deep stromal. After applying the gene-level filtering criteria, 132 candidate partner genes were retained for analysis. For each candidate partner, endothelial cells were matched across spatial contexts according to the joint distribution of log1p-transformed *CXCL12* and partner counts. Each epithelium-adjacent cell was greedily matched without replacement to the nearest intermediate and deep-stromal cells using Euclidean distance in the standardized two-dimensional log-count space, subject to a fixed caliper of 0.6. Incomplete triplets were discarded, yielding three equally sized, abundance-matched groups for group-level co-localization analysis. Co-localization scores were then computed separately for the matched epithelium-adjacent, intermediate and deep-stromal cells. For each candidate partner gene, the difference in *CXCL12* co-localization scores between the epithelium-adjacent and deep-stromal contexts was assessed using permutation tests based on 2,000 permutations of the spatial-context labels, followed by Benjamini-Hochberg correction across the 132 candidate genes. Significant partner genes were grouped according to whether *CXCL12* co-localization was stronger in epithelium-adjacent or deep-stromal cells.

### Perturbation effect prediction

We performed *in silico* perturbation experiments on the Xenium human breast cancer dataset using SVC trained on tissue section 1, and applied the trained model to both sections. Two sets of perturbations were designed. In the first, we depleted *LUM* in all stromal cells. In the second, we depleted *ERBB2* in all tumor-related cells (DCIS #1, DCIS #2, invasive tumor and proliferative invasive tumor cells). For every cell, the input of the depleted gene was clamped to its 5th percentile among the cells of that set. The same clamp was applied to that gene in the neighbor-expression context of every neighboring cell. For each analysis, we systematically quantified the predicted downstream responses of broadly expressed genes, defined as those detected in at least half of the corresponding cell population. This criterion retained 65 genes for *LUM* depletion and 60 for *ERBB2* depletion. Specifically, for each downstream gene, we masked the gene itself from the model input in a leave-one-out manner and quantified both the change in predicted cell-level expression and the change in subcellular localization after depletion of the given gene. The cell-specific size factor was computed from the unperturbed input and held fixed for both the unperturbed and perturbed predictions. For each of the two response measures, we summarized the perturbation effect for each gene across cells and assessed its significance relative to the distribution of effects across all analyzed genes using a median- and median absolute deviation-based robust z-score. Two-sided *P* values were derived from the standard normal distribution and corrected across genes using the Benjamini-Hochberg procedure. For the *LUM* perturbation we further asked whether the response depends on tissue context. We defined three stromal spatial contexts: epithelium-adjacent (domains 1, 5 and 16), intermediate (domains 0, 9, 11, 13 and 14), and deep stroma (domains 4, 6 and 8), based on the epithelial proximity of spatial domains. For each gene, we quantified the difference in mean localization change between epithelium-adjacent and deep-stromal cells and assessed its significance relative to the distribution of these differences across all analyzed genes using the same robust z-score procedure described above.

### Compared methods

A unique feature of SVC is its ability to model the subcellular spatial expression patterns of genes— an aspect not addressed by existing methods. As no comparable methods are available for direct evaluation of subcellular spatial pattern modeling, we instead assessed SVC’s performance through two relevant analytic tasks where benchmarking is feasible.

The first task focuses on the overall cell-specific gene expression prediction, evaluating expression prediction at the cellular level without subcellular spatial resolution. To achieve this, SVC generates single-cell gene expression prediction by summing the predicted pixel-level expression values across each cell image. In this analysis, we benchmarked SVC against four state-of-the-art spatial transcriptomics imputation methods: ENVI^16^, Tangram^17^, gimVI^18^, and uniPort^19^. All four methods leverage single-cell RNA-seq reference data to impute unmeasured genes at the single-cell level in spatial transcriptomic datasets. For these methods, we treated our training data as non-spatial scRNA-seq input and applied each method to impute gene expression in the test data. To ensure a fair comparison, we tuned the key hyperparameters of each baseline method using the validation data, as detailed in Supplementary Note 5. Additional implementation settings are described below; the remaining parameters followed the default settings of the original implementations.

- ENVI: we used the reference implementation (https://github.com/dpeerlab/ENVI) with negative-binomial likelihoods for both modalities (spatial_dist=“nb”, sc_dist=“nb”), and trained the model for 16,000 steps. Gene imputation was performed using the impute_genes function. The spatial neighborhood size (k_nearest) and the weight of the latent KL term (kl_coeff) were tuned for each dataset.
- Tangram: we used the reference implementation (https://github.com/broadinstitute/Tangram), with library-size normalization, and trained the model for 1,000 epochs with a uniform density prior (density_prior=“uniform”). For the two cultured-cell datasets we mapped individual reference cells (mode=“cells”); for the tissue-level Xenium datasets, we mapped cell-type clusters instead (mode=“clusters”, cluster_label=“cell_type”). For gene imputation at single-cell resolution, we normalized the mapping weights within each spatial cell and computed the weighted average of the mapped reference expression profiles. The weights of the density and gene-expression terms (lambda_d and lambda_g1) were tuned for each dataset.
- gimVI: we used the reference implementation (https://docs.scvi-tools.org/en/stable/), with negative-binomial generative distributions for both modalities (generative_distributions=[“nb”, “nb”]), and trained the model for 200 epochs. Raw count predictions were generated using the get_imputed_values function with the normalized parameter set to False. The latent dimension (n_latent) and library-size modeling (model_library_size) were tuned for each dataset.
- uniPort: we preprocessed data following the authors’ instructions (https://uniport.readthedocs.io/) using library-size normalization, log1p transformation, and the batch_scale function. We then trained the model for a maximum of 30,000 iterations and generated imputed expression with out=”predict”. Postprocessing included inverse batch scaling, expm1 transformation, and library size denormalization to recover raw counts. The weights of the KL and optimal-transport terms (lambda_kl and lambda_ot) were tuned for each dataset.

To evaluate performance, we calculated the root mean squared error (RMSE) and Pearson correlation coefficient (PCC) between the imputed and ground-truth expression values for every held-out gene across cells, using a K-fold cross-validation scheme over genes for each method.

The second task focuses on gene co-localization analysis, aiming to identify gene pairs that exhibit spatial co-localization patterns within cells. For this analysis, we compared SVC with InSTAnT^15^, a method that uses customized statistical tests to detect co-localized gene pairs. Given that there is no ground truth, we reasoned that gene pairs showing subcellular co-localization detected by SVC are likely, although not necessarily, also detected by InSTAnT. To quantify this agreement, we first computed the number of overlapping gene pairs among the top 50, 100, 200, 500, and 1,000 ranked gene pairs from each method. However, SVC’s latent representations are designed to capture gene-specific within-cell overall spatial pattern, whereas InSTAnT, which relies on statistical testing of local mRNA distances, primarily reflects short-range physical co-localization of individual transcripts. We therefore also examined differences between the top gene pairs detected by SVC and those detected by InSTAnT. Among gene pairs that were not shared between SVC and InSTAnT, we quantified the numbers of SVC-specific and InSTAnT-specific pairs that were supported by the 49 genes with annotated subcellular localizations from the original study as belonging to the same localization type, defined as one of three categories: nuclear/perinuclear, cytoplasmic, or protrusion. For InSTAnT, we followed its repository instructions (https://github.com/bhavaygg/InSTAnT) to set the ‘distance_threshold’ parameter to 4 and the ‘min_genecount’ parameter to 20. We used the ‘run_GlobalColocalization’ function to identify globally significant co-localized gene pairs, with a *P* value significance threshold of 0.05. We ranked gene pairs by their conditional *P* values ‘p_val_cond’ in ascending order and selected the top-ranking pairs accordingly.

### Implementation details of the SVC model

We implemented SVC using a Vision Transformer backbone with Performer encoder^65^. The encoder comprises 12 Performer layers with 12 attention heads per layer. For the benchmark analyses, SVC was trained from scratch independently on each of the four datasets (seqFISH+ mouse embryonic fibroblast, MERFISH U2-OS, Xenium mouse brain, and Xenium human breast cancer) and evaluated on the corresponding held-out cells or tissue sections. No externally pretrained model weights were used for model initialization. For all datasets, SVC was optimized using AdamW with a maximum learning rate of 1 × 10^−4^ . We used a single-cycle cosine-annealing learning-rate schedule, with the learning rate gradually decayed from 1 × 10^−4^ to 1 × 10^−6^ during training. The weight λ was set to zero for the first 50 epochs and increased linearly to 10 between epochs 50 and 100. Sensitivity analysis across a broad range of alternative λ settings showed that SVC performance was generally stable (Supplementary Fig. 79). Batch size was set to 32 for seqFISH+ and MERFISH U2-OS datasets. For the two Xenium datasets, we used a per-step batch size of 128 with two-step gradient accumulation, resulting in an effective batch size of 256.

Dataset-specific settings were used to account for differences between cultured-cell and tissue datasets. For seqFISH+ and MERFISH U2-OS, because each field of view contains only on the order of ten cells, a single neighborhood scale with k = 4 was used. As these cultured cells have no fixed anatomical orientation, we further applied rotation and reflection augmentation during training by randomly applying dihedral transformations to the subcellular gene images and the corresponding cell and nuclear morphology images. For the two Xenium tissue datasets, we used multi-scale neighbors with k = 4, 16 and 64, and concatenated the profiles pooled at the three scales before mapping them to a cell neighbor embedding.

For each of the four datasets, we held out a validation set from the training data for model selection. The test data were used only for the final performance evaluation. During training, we recorded both training and validation losses, used a maximum of 400 epochs, and applied early stopping if the validation loss did not improve for 50 consecutive epochs. The checkpoint with the lowest validation loss was used for evaluation. The corresponding training and validation loss curves for each dataset are provided in Supplementary Fig. 80.

### Analyzed datasets

We applied our method to eight publicly available subcellular-resolution spatial transcriptomics datasets generated using different technologies. For all datasets, cell segmentations were provided by the original studies, whereas nuclear segmentations were provided for all datasets except the MERFISH mouse brain section, for which we generated nuclear masks from the DAPI channel. Nuclear centers were then determined as the centroid of the nuclear segmentation coordinates. Binary cell and nuclear morphology masks used as model input were rasterized from these segmentations.

### SeqFISH+ mouse embryonic fibroblast data

The processed NIH/3T3 mouse embryonic fibroblast dataset profiled by seqFISH+^5^ was sourced from Bento^13^ following preprocessing via ELLA^14^. The dataset contains 2,747 genes and 171 cells across 17 FOV batches, and 2,659 genes are retained within defined gene co-expression networks from Gene2vec^60^. As Gene2vec was primarily pretrained on human genes, we first mapped the 16,906-gene set covered by Gene2vec to mouse orthologs and used the resulting ortholog-matched vectors. For our analysis, we focused on the top 1,000 highly expressed genes. We used batch 1 for validation, batch 3 as the holdout test set, and the remaining batches for training. We also obtained the corresponding cell cycle annotations from ELLA as cell identity labels for model input.

### MERFISH human osteosarcoma data

We obtained the processed human osteosarcoma data^20^ profiled by MERFISH from Bento^13^. The dataset contains 130 genes and 1,153 U2-OS cells across 70 imaging FOV batches. For our analysis, we retained 1,022 cells with complete nuclear morphology information and focused on 119 genes within defined gene co-expression networks from Gene2vec. We used batches 1-5 for validation, batches 61-68 as the holdout test set, and the remaining batches for training. As this dataset lacks sufficient cell-cycle-related marker genes, we did not provide additional cell identity labels to the model.

### Xenium mouse brain data

We obtained the tissue sections 1-2 from the mouse brain data profiled by Xenium. We removed negative control transcripts, blanks, and antisense transcripts, and focused on 228 genes within defined gene co-expression networks from Gene2vec, applying the same ortholog-mapping procedure to the mouse dataset as for the seqFISH+ data. For our analysis, we retained cells with total transcript counts greater than 200, resulting in 93,447 and 88,620 cells in the two tissue sections. Section 1 served as the training section, with 10% of its spatial grid regions held out for validation, while section 2 served as the independent test set. We obtained cell-type annotations for model input via SingleR^70^, with reference profiles from the Mouse Whole Cortex and Hippocampus SMART-seq dataset.

### Xenium Prime 5K mouse brain hemisphere data

We obtained the mouse brain hemisphere data profiled with Xenium using the 5K mouse pan tissue and pathways panel. We retained 144 genes overlapping with the Xenium mouse brain data listed above. Cells with total transcript counts greater than 50 and complete nuclear morphology information were retained, resulting in 18,783 cells. We obtained cell-type annotations for model input via SingleR, with reference profiles from the Mouse Whole Cortex and Hippocampus SMART-seq dataset.

### Xenium Prime 5K P301S tauopathy mouse brain data

We obtained mouse brain data from a P301S tau transgenic mouse and a wild-type control (GEO accession GSE330686). Both sections were obtained from 12-month-old female mice and profiled using the same Xenium Prime 5K Mouse Pan Tissue and Pathways panel^69^. We retained 144 genes shared with the Xenium mouse brain dataset described above. We obtained cell-type annotations for model input via SingleR, with reference profiles from the Mouse Whole Cortex and Hippocampus SMART-seq dataset. For our analysis, we retained cells with more than 20 transcripts and complete nuclear morphology information and focused on two cell populations involved in the neuroinflammatory response. This resulted in 568 microglia and perivascular macrophages (Micro-PVM; 175 wild-type and 393 tau transgenic) and 9,472 astrocytes (Astro; 4,382 wild-type and 5,090 tau transgenic).

### MERFISH mouse brain data

We obtained a coronal adult mouse brain section profiled by MERFISH with a 1,147-gene panel from the Brain Image Library (accession 293cc39ceea87f6d; mouse 2, section co2_14). We retained the 130 genes shared with the Xenium mouse brain panel. Transcripts were assigned to provider-defined cells within each z-plane and collapsed across five z-planes to generate 2D gene maps. Cells with more than 50 assigned transcripts across the 130 genes were retained, resulting in 24,957 cells. Nuclear masks were derived from the DAPI channel by intensity thresholding within each provider-defined cell contour in each z-plane. Gene maps and cell morphology were centered on the nuclear center. This dataset was used to evaluate cross-platform transfer of the model trained on the Xenium mouse brain data (Supplementary Note 12). We obtained cell-type annotations for model input via SingleR, with reference profiles from the Mouse Whole Cortex and Hippocampus SMART-seq dataset.

### Xenium human breast cancer data

We obtained the two tissue sections from breast cancer data profiled by Xenium^6^. We removed negative control transcripts, blanks, and antisense transcripts, and retained 294 genes with Gene2vec embeddings for analysis. Cells with total transcript counts greater than 200 were retained, resulting in 60,914 and 43,252 cells in the two tissue sections. Section 1 served as the training section, with 10% of its spatial grid regions held out for validation, while section 2 served as the independent test set. We also obtained the corresponding cell-type annotations as cell identity labels from the original study^6^ for model input.

### Xenium human lung cancer data

We obtained the human lung cancer section profiled by Xenium with the 377-gene panel from 10x Genomics. We removed negative control transcripts, blanks, antisense transcripts and transcripts not assigned to a cell, retained transcripts with a quality score of at least 20, and focused on the 141 genes shared with the Xenium human breast cancer panel described above. For our analysis, we retained cells with more than 50 transcripts within these 141 shared genes and complete nuclear morphology information, resulting in 5,385 cells. This dataset was used only to evaluate cross-tissue transfer of the model trained on the Xenium human breast cancer data (Supplementary Note 12). As no cell-type annotation compatible with the breast cancer labels is available for this section, the graph-based cell clusters provided with the dataset were mapped to breast cancer cell types based on cosine similarity between their mean expression profiles across the shared genes, and the resulting labels were used as cell identity input.

## Data availability

The original public data used in this study can be accessed through the following links:

1. SeqFISH+ mouse embryonic fibroblast data and (2) MERFISH human osteosarcoma data at https://doi.org/10.6084/m9.figshare.15109236;
2. Mouse brain data by 10x Xenium at https://www.10xgenomics.com/datasets/fresh-frozen-mouse-brain-replicates-1-standard;
3. Prime 5K mouse brain hemisphere data by 10x Xenium at https://www.10xgenomics.com/datasets/xenium-prime-fresh-frozen-mouse-brain;

1. Prime 5K P301S tauopathy mouse brain data by 10x Xenium at https://www.ncbi.nlm.nih.gov/geo/query/acc.cgi?acc=GSE330686;
4. Mouse Whole Cortex and Hippocampus SMART-seq data used as reference data for mouse brain cell-type annotation are available at https://portal.brain-map.org/atlases-and-data/rnaseq/mouse-whole-cortex-and-hippocampus-smart-seq;
5. Human breast cancer by 10x Xenium at https://www.10xgenomics.com/products/xenium-in-situ/preview-dataset-human-breast;
6. Mouse brain data by MERFISH at https://download.brainimagelibrary.org/29/3c/293cc39ceea87f6d/;
7. Human lung cancer by 10x Xenium at https://www.10xgenomics.com/datasets/preview-data-ffpe-human-lung-cancer-with-xenium-multimodal-cell-segmentation-1-standard.

## Code availability

The SVC software program and source code have been deposited at https://xiangzhou.github.io/software/ and https://github.com/aster-ww/SVC. Code for reproducing the figures in the manuscript is available at https://github.com/aster-ww/SVC-reproducibility.

## Acknowledgements

This study was supported by the National Institutes of Health (NIH) grants R01GM126553 and R01HG009124.

## Author contributions

X.Z. and H.W. conceived the idea and designed the project. H.W. developed the method and implemented the software. H.W., P.Y. and S.H. performed data analysis. P.Y. contributed to method evaluation and control analyses. S.H. contributed to validation analyses. J.W. contributed to data collection and preprocessing. H.W. drafted the manuscript. X.Z. supervised the project, provided funding support, wrote and edited the manuscript. All authors reviewed and approved the final manuscript.

## Ethics declarations

