## Supplementary Information for "SVC: A Vision Transformer-based Spatially and Virtually Embedded Cell Model for Deciphering Subcellular Spatial Transcriptomic Heterogeneity"

### Supplementary Figures

**Supplementary Figure 1. Ablation study of SVC model components.** Each ablated variant was retrained from scratch with five random seeds using the same settings as the full model. The four variants each remove a single embedding (cell morphology, cell neighbor, cell identity, or gene functional) while keeping all other components unchanged. In the cell-level-only setting, the subcellular gene localization embedding was ablated and replaced by cell-level gene expression counts, so that the model no longer received subcellular spatial distribution information; in the subcellular-only setting, only the subcellular gene localization embedding was retained, with the morphology, neighbor, cell identity, and gene-level functional embeddings all removed. Performance is reported as median PCC (top row; higher is better) and median RMSE (bottom row; lower is better) computed across cells, across genes, and within subcellular spatial maps. Cross-cell metrics compare total transcript counts across cells for each gene; cross-gene metrics compare expression profiles across genes within each cell; and subcellular metrics compare 12 × 12 spatial expression maps for each cell-gene pair. Bars show the mean across five seeds and error bars indicate the standard deviation. **a.** seqFISH+. **b.** MERFISH U2-OS.

a

b

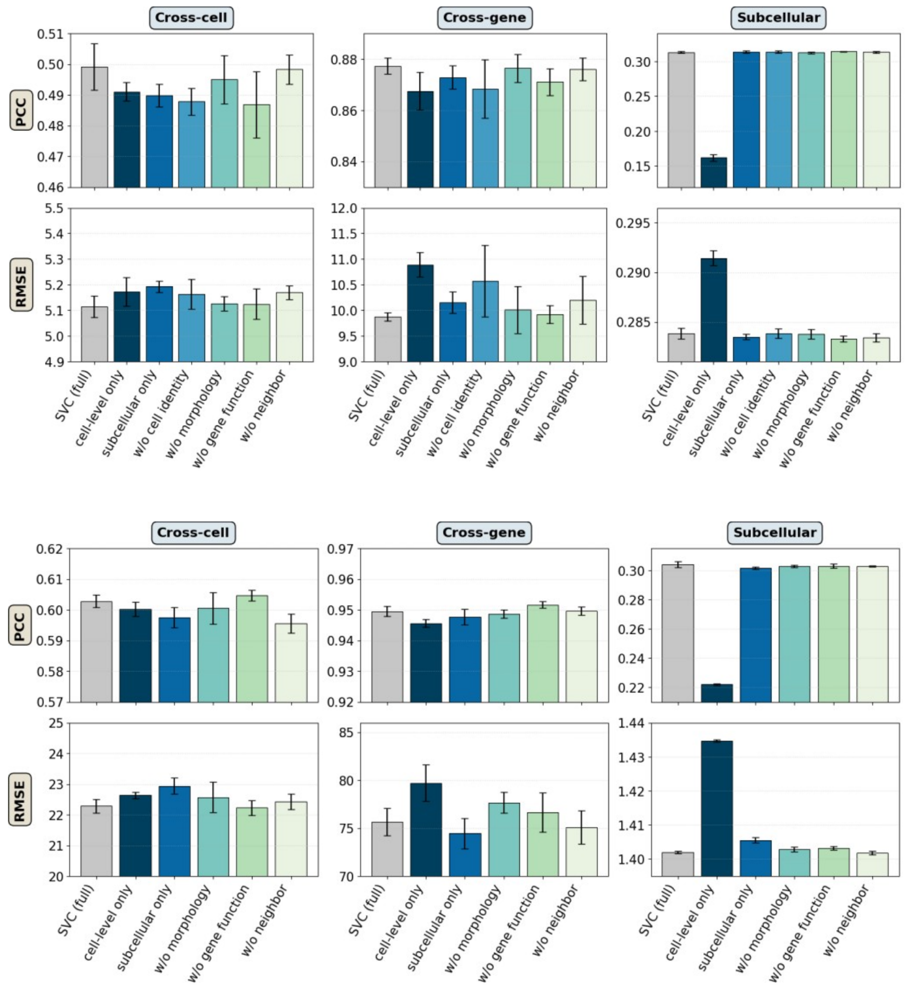

**Supplementary Figure 2. Computational scalability of SVC.** SVC was trained on random subsamples of increasing size from the Xenium mouse brain dataset. Left, total training time to convergence (left axis) and average time per epoch (right axis) across different numbers of training cells. Right, peak GPU memory per device across different numbers of training cells, measured over the entire run or during the training step alone, excluding cached validation data.

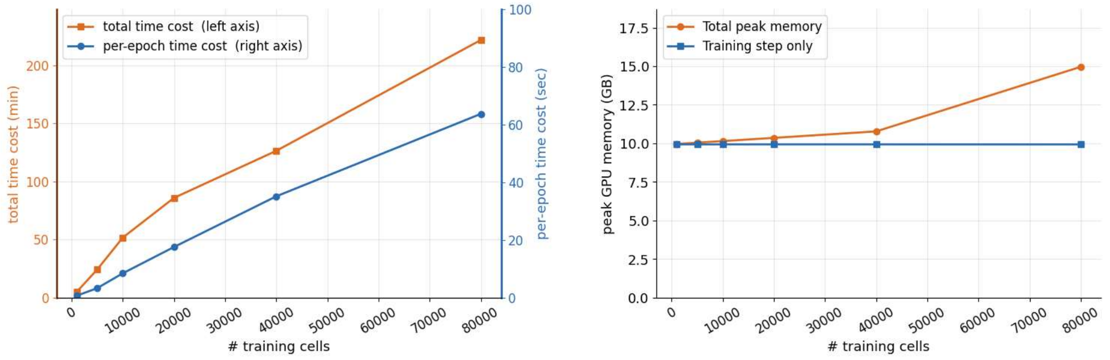

**Supplementary Figure 3. Gene subcellular pattern prediction performance comparison between SVC and four baseline models in the seqFISH+ mouse embryonic fibroblast data.** Box plots summarize Pearson correlation (PCC), cosine similarity, and RMSE across genes on the held-out test data. For PCC and cosine similarity, higher values indicate better agreement; for RMSE, lower is better.

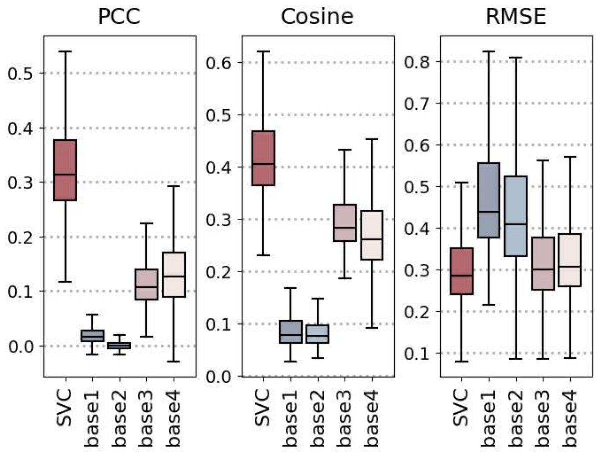

**Supplementary Figure 4. Gene subcellular pattern prediction performance of SVC and four baseline models in the seqFISH+ mouse embryonic fibroblast data, evaluated using distribution-based metrics. a.** Box plots show the earth mover's distance (EMD), Kullback-Leibler divergence (KL), and Jensen-Shannon divergence (JS) across genes in the held-out test set, computed between the predicted and measured subcellular distributions. For all three metrics, lower values indicate better agreement. **b.** Box plots show the per-gene performance gains of SVC over the four baseline models for EMD, KL, and JS, with positive values indicating better performance by SVC.

a b

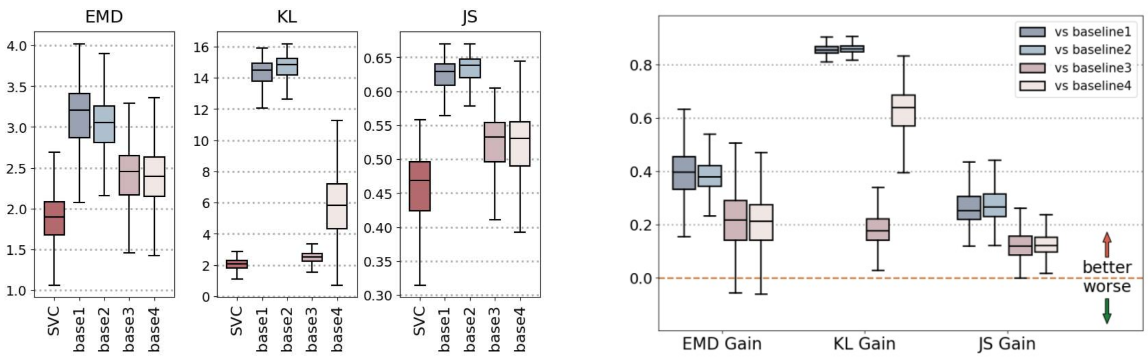

**Supplementary Figure 5. Subcellular localization prediction performance across transcript abundance levels in the seqFISH+ mouse embryonic fibroblast dataset.** **a.** Stochastic variation and systematic bias in the difference between predicted and measured localization, across gene-expression levels. For each gene in each test cell, the predicted and measured transcript maps were separately normalized, and their difference was projected onto three spatial coordinates: radial distance from the nuclear center and signed horizontal and vertical pixel coordinates (x and y axes). For each gene, stochasticity was quantified as the standard deviation of the signed displacement across cells, whereas bias was quantified as its mean. Box plots show the distributions of these gene-level statistics within five equal-sized expression quintiles. **b.** Subcellular localization accuracy of SVC and four baseline models across the same expression quintiles. **c.** Per-gene subcellular prediction accuracy across expression level for SVC and Baseline 4.

**a**

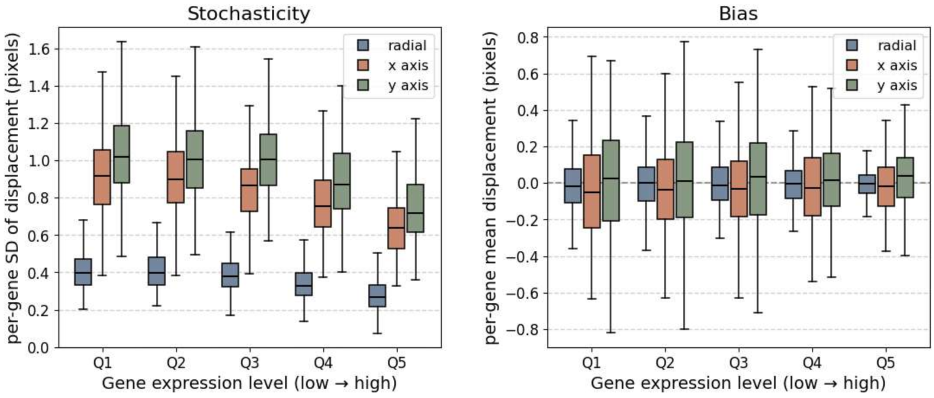

**b**

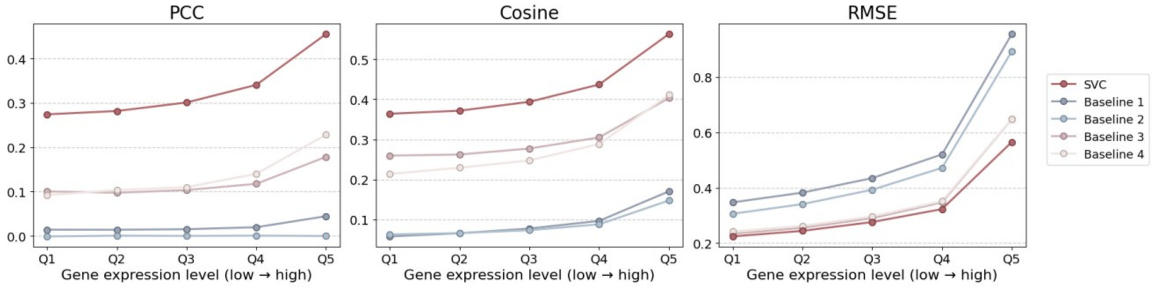

**c**

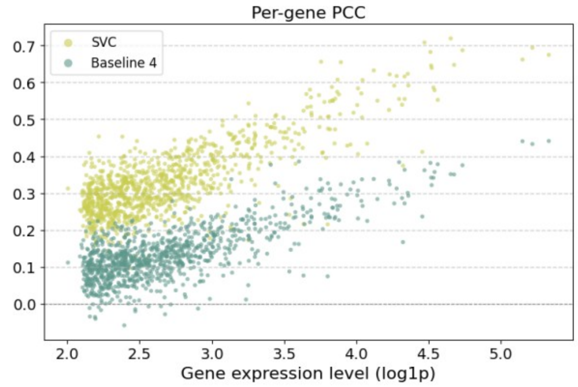

**Supplementary Figure 6. Comparison of SVC with mean-distance baselines in the seqFISH+ mouse embryonic fibroblast data.** Two mean-distance predictors were evaluated: baseline (train), which uses the mean distance of each gene across training cells, and baseline (test), which uses the leave-one-out mean distance of each gene across the other test cells. **a.** Scatter plots compare predicted and measured mean distances to the nuclear center for 49 genes with annotated subcellular localization patterns. Colors indicate different subcellular localization categories, and Pearson correlation coefficients are shown in each panel. For this gene-level comparison, baseline (test) reduces to the measured test-set mean and therefore serves as an oracle reference. The corresponding SVC predictions are shown in Fig. 2b. **b.** Relative reduction in RMSE achieved by SVC over each baseline across individual cells for each gene. Positive values indicate better performance by SVC.

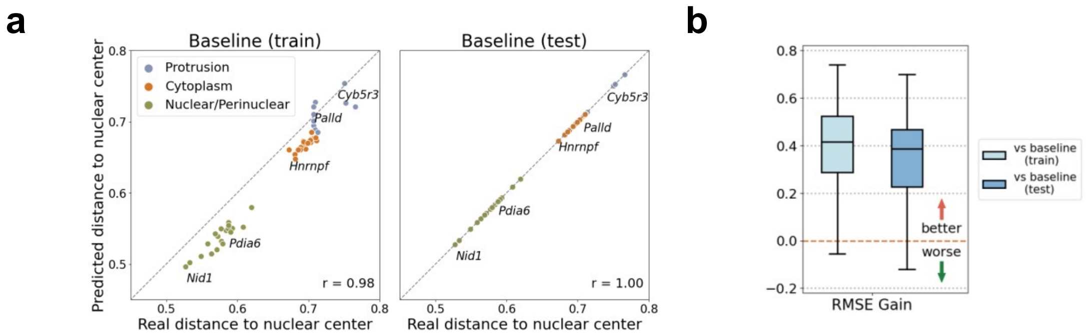

**Supplementary Figure 7. Hyperparameter tuning for competing methods.** For each competing method, we performed a grid search over two key hyperparameters separately for each dataset, using the same train/test split and gene cross-validation protocol as in the SVC evaluation. Hyperparameter settings were evaluated on a held-out validation set that was not used for final testing. Heatmap colors indicate the validation score, which combines per-gene RMSE and PCC after normalization, with higher values indicating better performance. In each panel, the asterisk denotes the best-performing setting selected for subsequent benchmark comparisons, whereas the triangle indicates the method's default setting. Rows correspond to competing methods and columns to datasets.

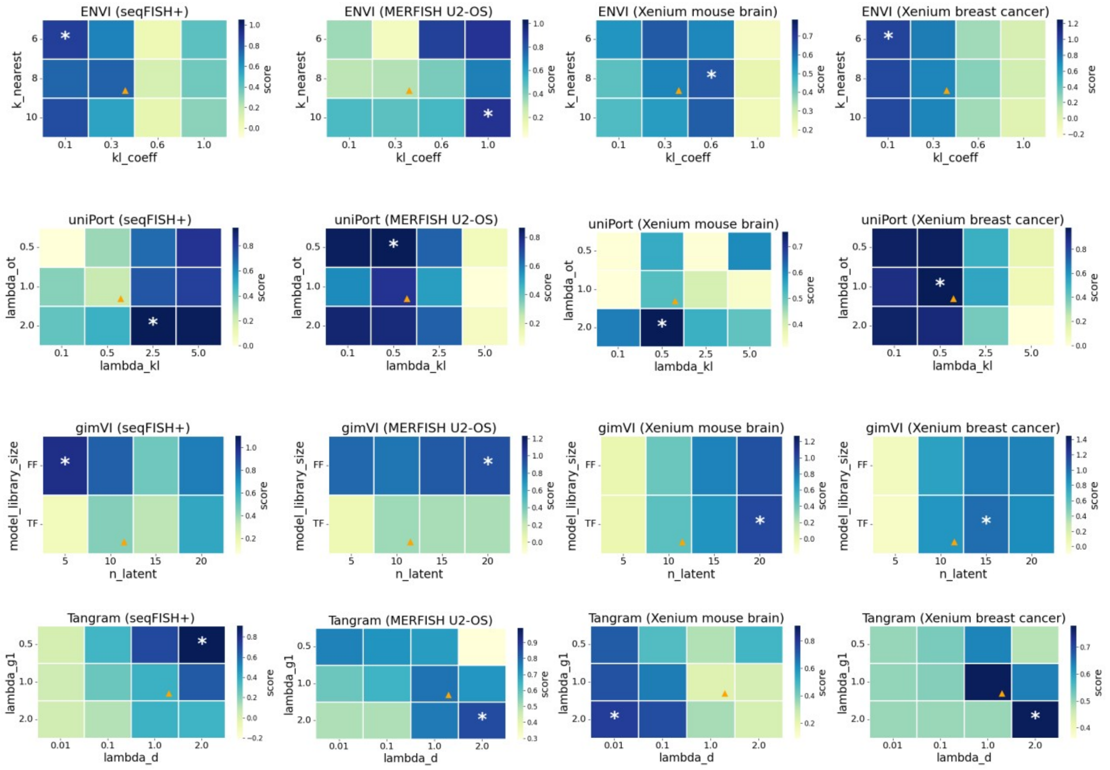

**Supplementary Figure 8. Cell-level gene expression prediction accuracy of compared methods in the seqFISH+ mouse embryonic fibroblast data.** Scatter plots compare predicted versus observed expression levels on a log-transformed scale, for uniPort, ENVI, Tangram, and gimVI on the test data, with each dot representing a gene. Pearson correlations are calculated on the original scale across all genes.

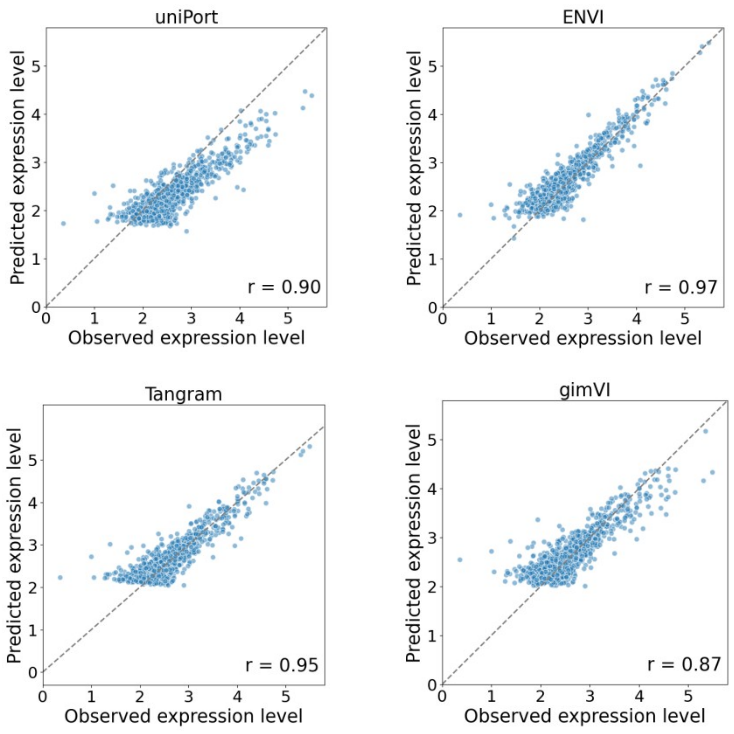

**Supplementary Figure 9. Subcellular gene co-localization analysis in the seqFISH+ mouse embryonic fibroblast data.** Observed transcript profiles within original cells for three additional representative co-localization gene pairs.

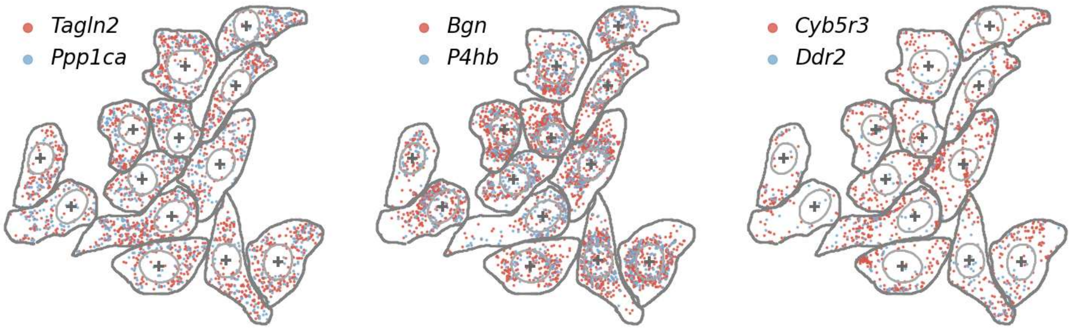

**Supplementary Figure 10. Visualization of three nuclear-associated subclusters in the seqFISH+ mouse embryonic fibroblast data.** Observed transcript profiles within original cells for three nuclear-associated subclusters provided by the original study[1], using 11 genes outside the 49 well-annotated gene set.

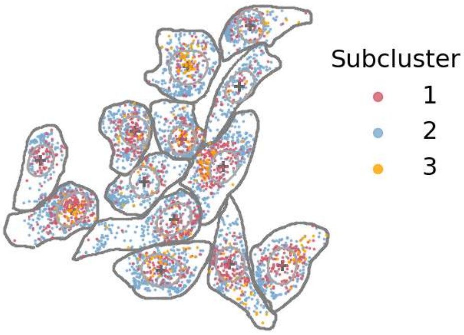

**Supplementary Figure 11. Subcellular gene co-localization analysis of three nuclear-associated subclusters in the seqFISH+ mouse embryonic fibroblast data.** Heatmap visualizes subcellular co-localization scores for 11 genes belonging to the aforementioned three subclusters.

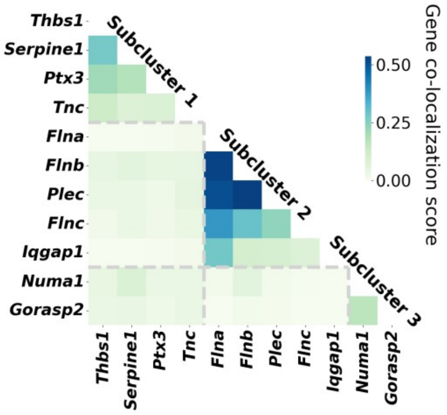

**Supplementary Figure 12. UMAP visualization of nuclear-associated subclusters in the seqFISH+ mouse embryonic fibroblast data.** The UMAP embedding was generated from the mean latent representation of each gene across training cells.

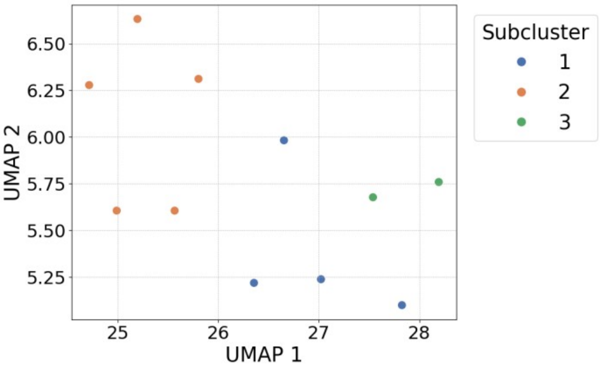

**Supplementary Figure 13. Cell area and eccentricity across cell-cycle phases in the seqFISH+ mouse embryonic fibroblast data.** Each point represents a cell, positioned according to its cell-mask area and eccentricity and colored by its annotated cell-cycle phase.

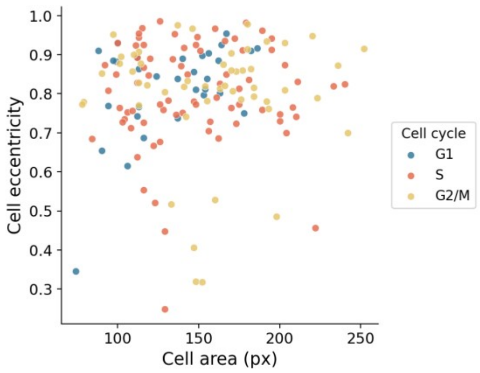

**Supplementary Figure 14. Gene co-localization score by cell-cycle phase under two morphology controls, in the seqFISH+ mouse embryonic fibroblast data. a.** Co-localization scores computed from an SVC variant trained without morphology input. **b.** Co-localization scores computed after residualizing each gene's latent representation on morphology features, taken as the first 20 principal components of the raw cell and nuclear mask images. In both panels, box plots show the distributions of co-localization scores for cytoplasmic-localized, nuclear/perinuclear-localized, and protrusion-related genes averaged across cells in G1, S, and G2/M phase. Statistical significance for pairwise comparisons (n.s. ≥ 0.05; * < 0.05; ** < 0.01; *** < 0.001) was assessed using one-sided Wilcoxon signed-rank tests.

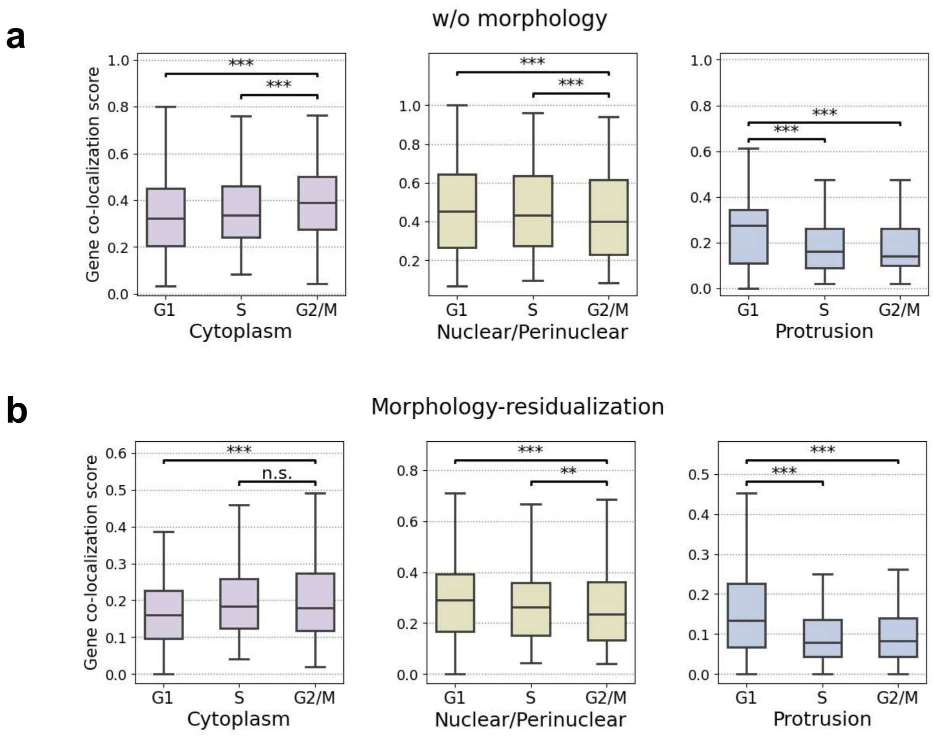

**Supplementary Figure 15. Measured *Ddr2* transcripts in representative cells from each cell-cycle phase in the seqFISH+ mouse embryonic fibroblast data.**

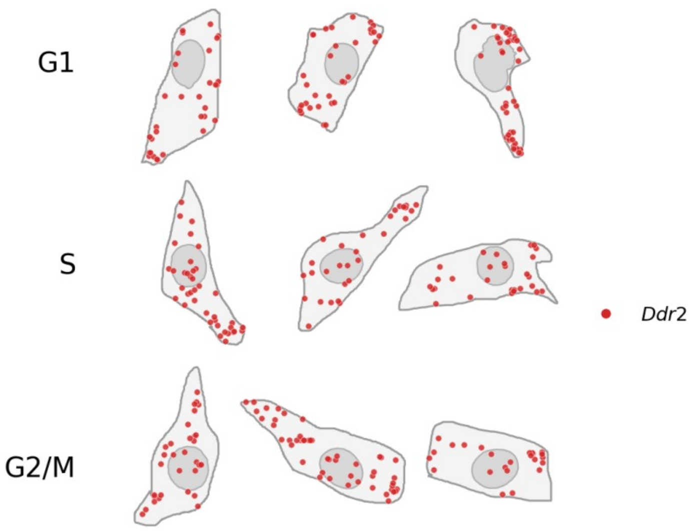

**Supplementary Figure 16. Gene subcellular pattern prediction performance of SVC and four baseline models in the MERFISH human osteosarcoma data.** Box plots summarize Pearson correlation (PCC), cosine similarity, and RMSE across genes on the held-out test data. For PCC and cosine similarity, higher values indicate better agreement; for RMSE, lower is better.

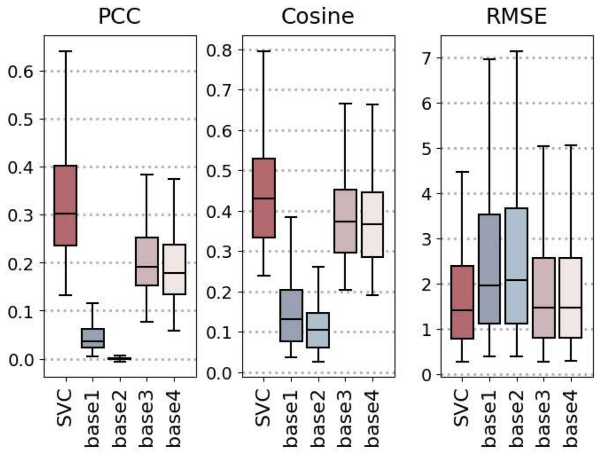

**Supplementary Figure 17. Gene subcellular pattern prediction performance of SVC and four baseline models in the MERFISH human osteosarcoma data, evaluated using distribution-based metrics. a.** Box plots show the earth mover's distance (EMD), Kullback-Leibler divergence (KL), and Jensen-Shannon divergence (JS) across genes in the held-out test set, computed between the predicted and measured subcellular distributions. For all three metrics, lower values indicate better agreement. **b.** Box plots show the per-gene performance gains of SVC over the four baseline models for EMD, KL, and JS, with positive values indicating better performance by SVC.

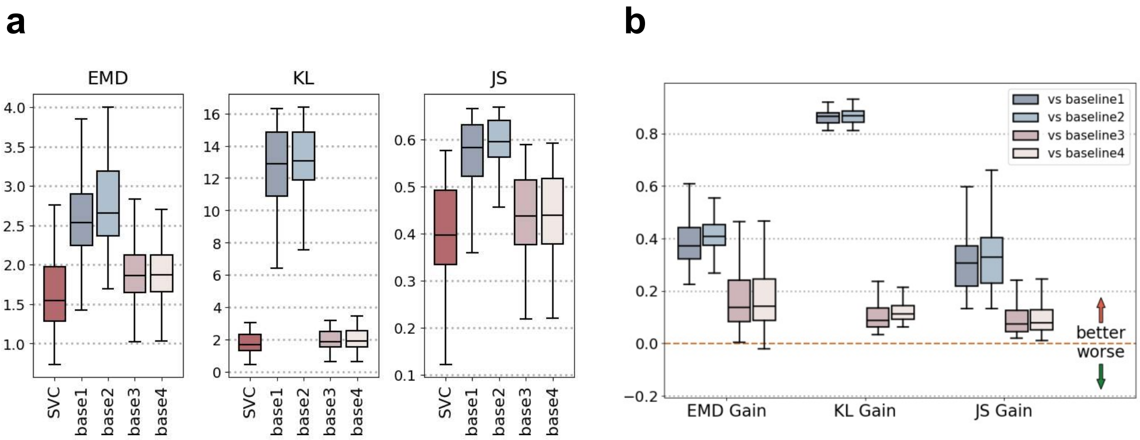

**Supplementary Figure 18. Agreement between predicted and observed subcellular localization in the MERFISH human osteosarcoma data.** Scatter plot compares the SVC-predicted relative distance to the nuclear center against the truth values across genes (Pearson correlation = 0.99).

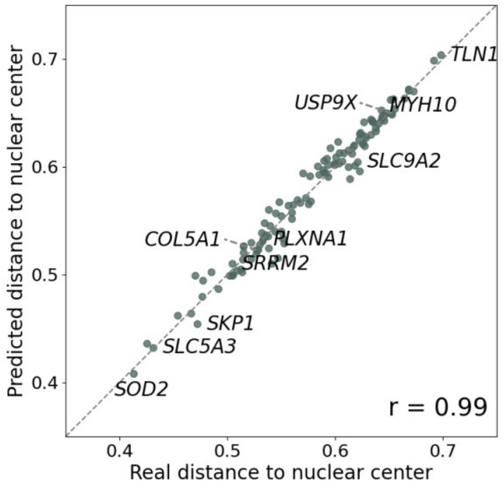

**Supplementary Figure 19. Comparison of SVC with mean-distance baselines in the MERFISH human osteosarcoma data.** Two mean-distance predictors were evaluated: baseline (train), which uses the mean distance of each gene across training cells, and baseline (test), which uses the leave-one-out mean distance of each gene across the other test cells. **a.** Scatter plots compare predicted and measured mean distances to the nuclear center across all measured genes, with Pearson correlation coefficients shown in each panel. Because this comparison uses gene-level averages, baseline (test) reduces to the measured test-set mean and therefore serves as an oracle reference. **b.** Relative reduction in RMSE achieved by SVC over each baseline across individual cells for each gene. Positive values indicate better performance by SVC.

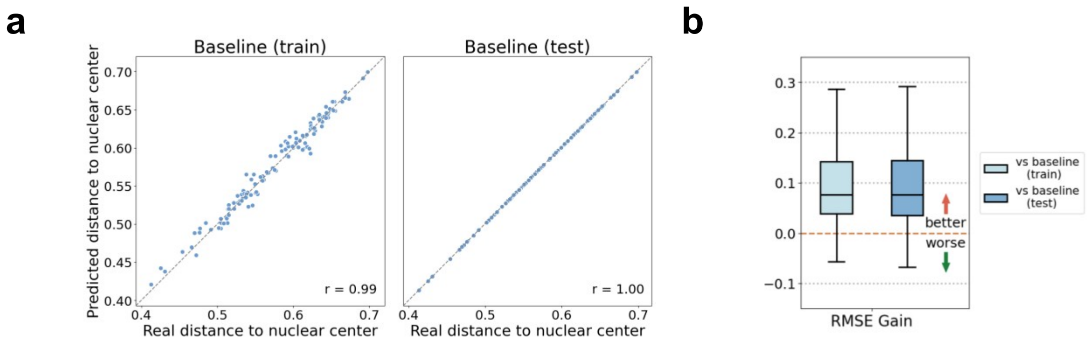

**Supplementary Figure 20. Visualization of SVC subcellular gene pattern predictions in the MERFISH human osteosarcoma data.** SVC-predicted transcript profiles are visualized within the original cells alongside the ground truth for five example genes in the held-out batch 64 test data. To improve visibility under point overlap, we added zero-mean Gaussian jitter with standard deviation 10 to each transcript's coordinates.

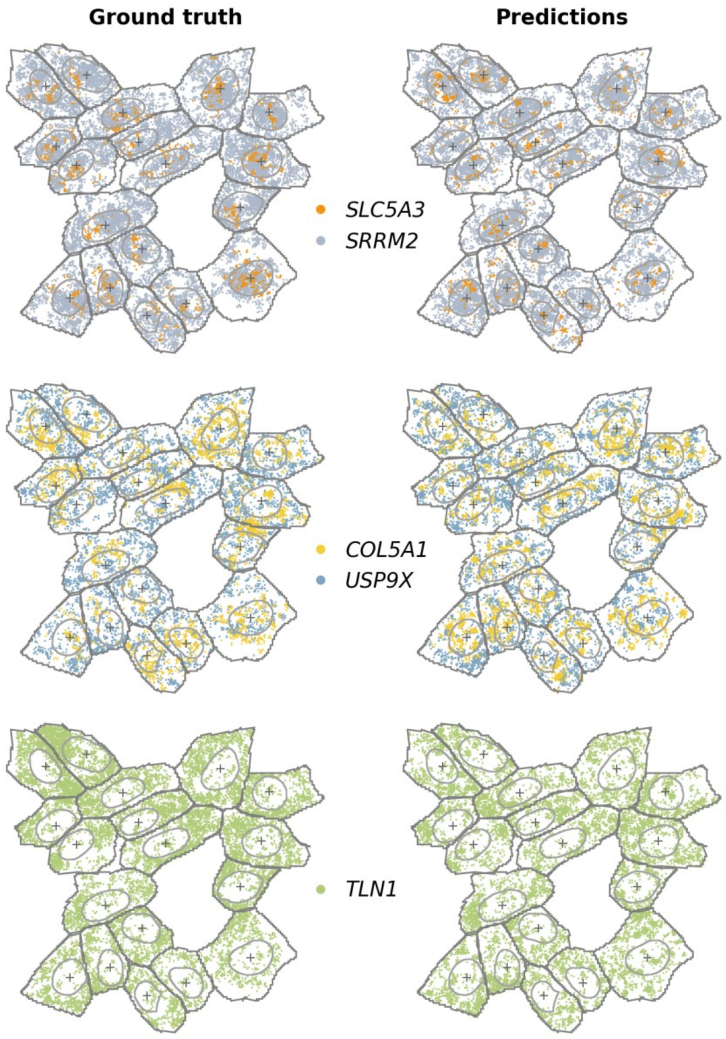

**Supplementary Figure 21. Cell-level gene expression prediction accuracy comparisons in the MERFISH human osteosarcoma data.** For the cell-level gene expression prediction task, box plots show the per-gene RMSE (log1p-transformed) and PCC scores between observed and predicted expression counts using 10 held-out gene folds. SVC is compared with uniPort, ENVI, Tangram, and gimVI.

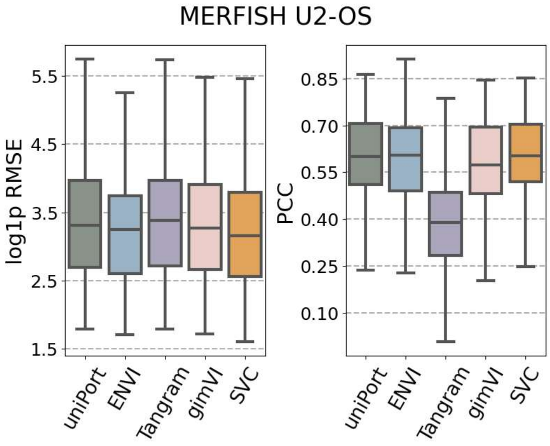

**Supplementary Figure 22. GO enrichment analysis of the ten lowest-entropy genes in the MERFISH human osteosarcoma data.** Bar plot shows the -log10(*P*) values of GO terms with *P* value < 0.01 among the ten lowest-entropy genes.

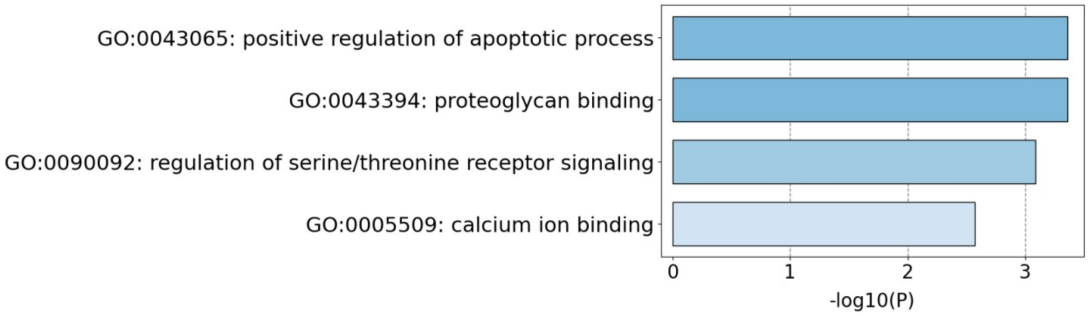

**Supplementary Figure 23. Robustness of SVC to spatial noise in the seqFISH+ mouse embryonic fibroblast data. a.** Example measured subcellular maps of three genes at increasing noise level. **b.** Relative change in PCC (top) and RMSE (bottom) with respect to the unperturbed input as a function of noise level, with 50% of images jittered. **c.** Relative change in PCC (top) and RMSE (bottom) with respect to the unperturbed input, as a function of the fraction of jittered images, at a noise level of 1.0. Cross-cell metrics compare total transcript counts across cells for each gene; cross-gene metrics compare expression profiles across genes within each cell; and subcellular metrics compare 12 × 12 spatial expression maps for each cell-gene pair. Lines show the mean across five random seeds and error bars indicate the standard deviation.

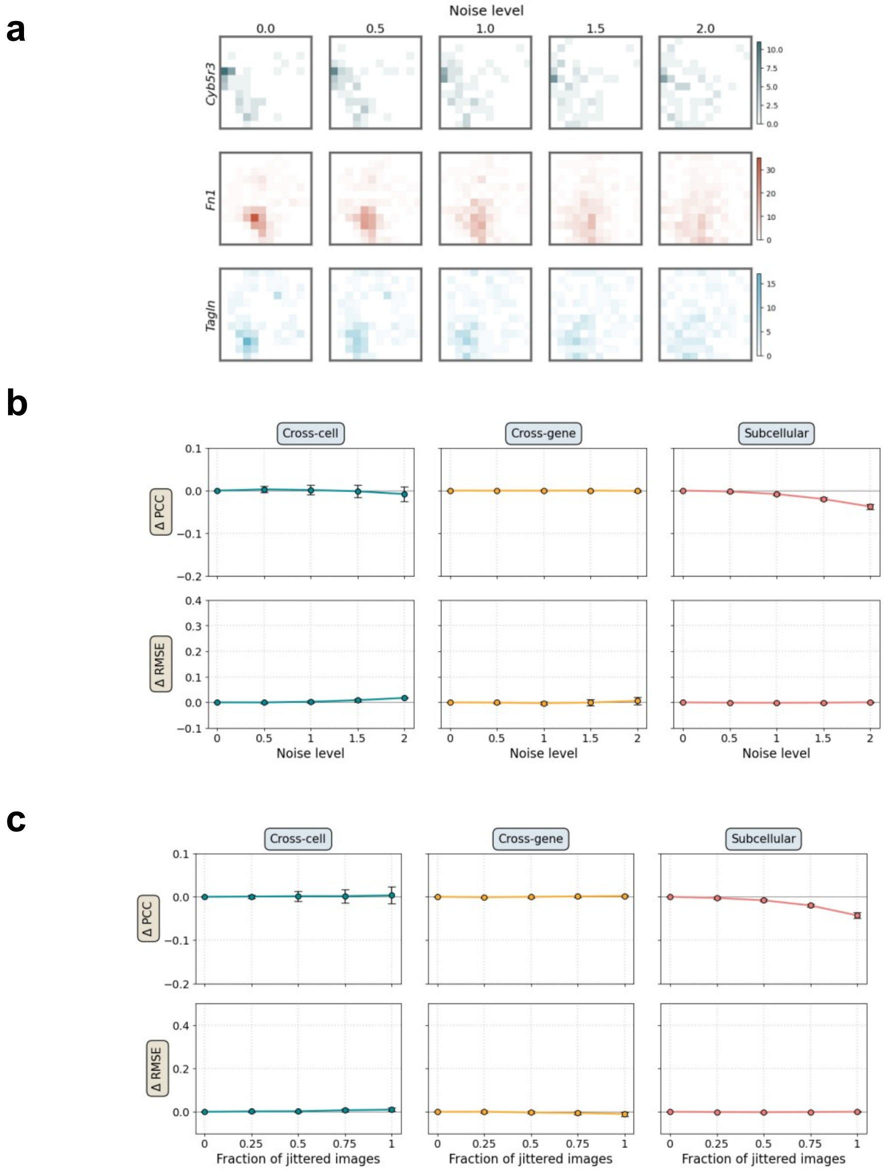

**Supplementary Figure 24. Robustness of SVC to morphology perturbation in the seqFISH+ mouse embryonic fibroblast data. a.** Example cell masks at increasing noise level, where the boundary of each cell and its nucleus is elastically deformed with a root-mean-square displacement given by the noise level, in pixels; the dashed outline marks the unperturbed boundary. **b.** Relative change in PCC (top) and RMSE (bottom) with respect to the unperturbed input, as a function of noise level, with 50% of cells perturbed. **c.** Relative change in PCC (top) and RMSE (bottom) with respect to the unperturbed input, as a function of the fraction of cells whose cell and nuclear masks are both replaced by those of a randomly selected training cell. Cross-cell metrics compare total transcript counts across cells for each gene; cross-gene metrics compare expression profiles across genes within each cell; and subcellular metrics compare 12 × 12 spatial expression maps for each cell-gene pair. Lines show the mean across five random seeds and error bars indicate the standard deviation.

**a**

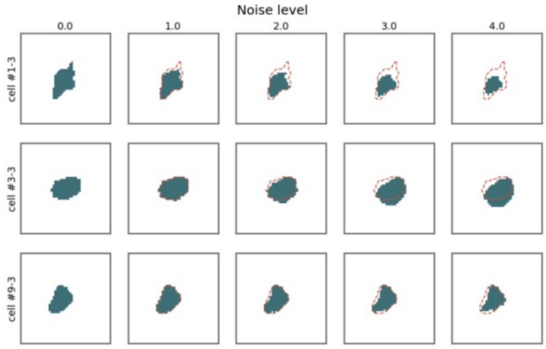

**b**

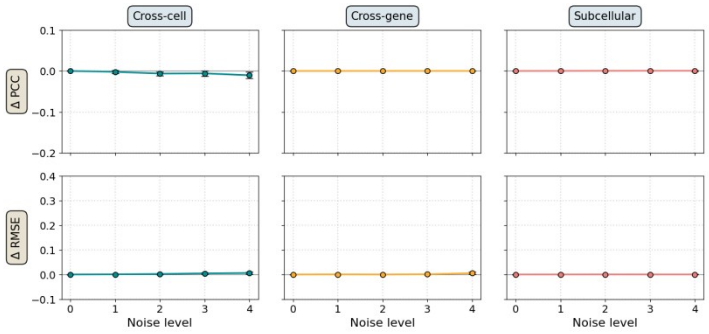

**c**

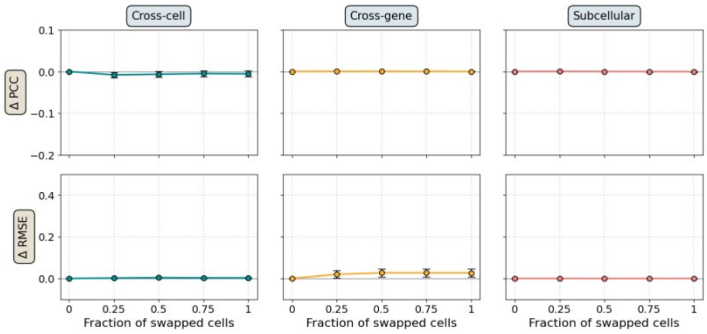

**Supplementary Figure 25. Robustness of SVC to spatial noise in the MERFISH human osteosarcoma data. a.** Example measured subcellular maps of three genes at increasing noise level. **b.** Relative change in PCC (top) and RMSE (bottom) with respect to the unperturbed input as a function of noise level, with 50% of images jittered. **c.** Relative change in PCC (top) and RMSE (bottom) with respect to the unperturbed input, as a function of the fraction of jittered images, at a noise level of 1.0. Cross-cell metrics compare total transcript counts across cells for each gene; cross-gene metrics compare expression profiles across genes within each cell; and subcellular metrics compare 12 × 12 spatial expression maps for each cell-gene pair. Lines show the mean across five random seeds and error bars indicate the standard deviation.

a

b

c

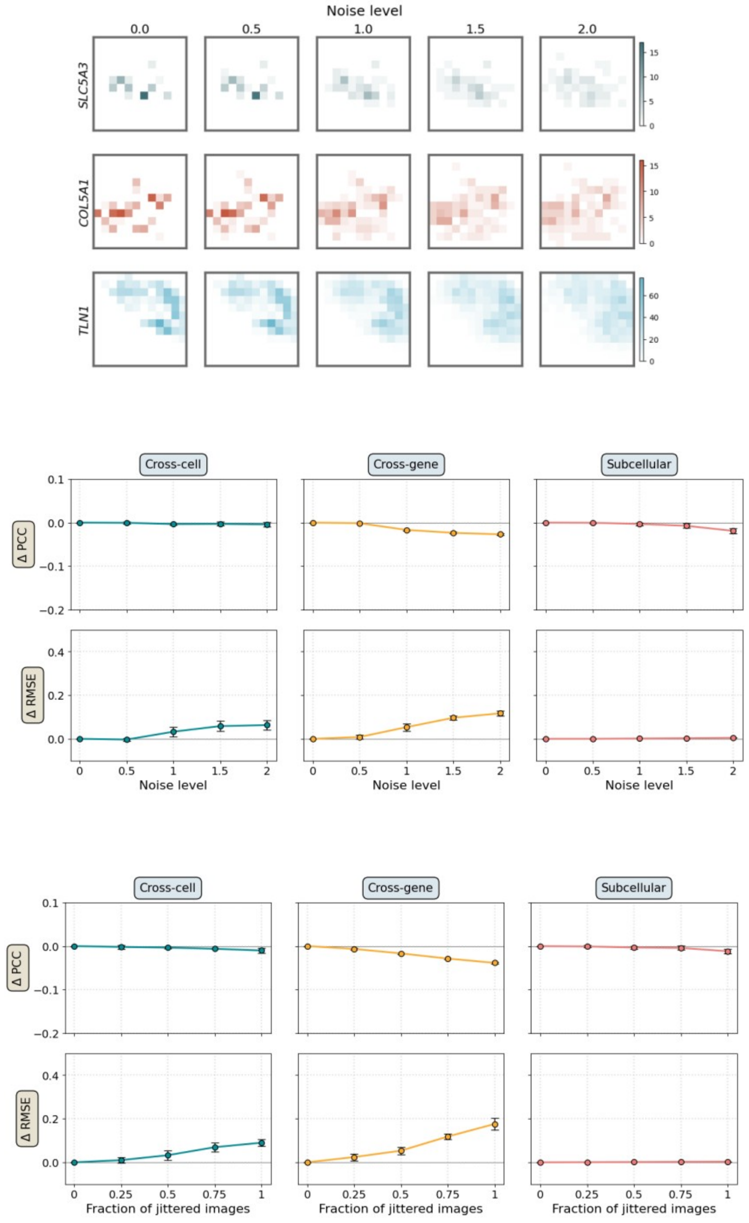

**Supplementary Figure 26. Robustness of SVC to morphology perturbation in the MERFISH human osteosarcoma data. a.** Example cell masks at increasing noise level, where the boundary of each cell and its nucleus is elastically deformed with a root-mean-square displacement given by the noise level, in pixels; the dashed outline marks the unperturbed boundary. **b.** Relative change in PCC (top) and RMSE (bottom) with respect to the unperturbed input, as a function of noise level, with 50% of cells perturbed. **c.** Relative change in PCC (top) and RMSE (bottom) with respect to the unperturbed input, as a function of the fraction of cells whose cell and nuclear masks are both replaced by those of a randomly selected training cell. Cross-cell metrics compare total transcript counts across cells for each gene; cross-gene metrics compare expression profiles across genes within each cell; and subcellular metrics compare 12 × 12 spatial expression maps for each cell-gene pair. Lines show the mean across five random seeds and error bars indicate the standard deviation.

**a**

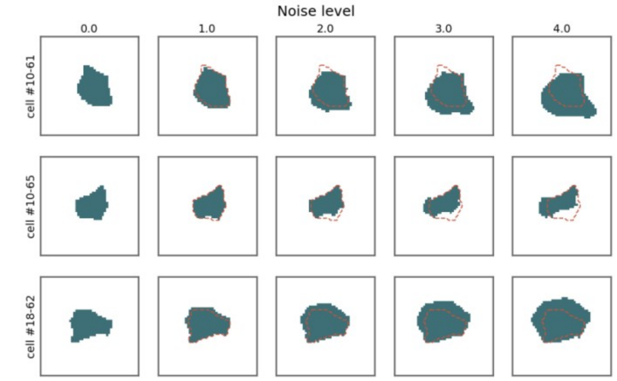

**b**

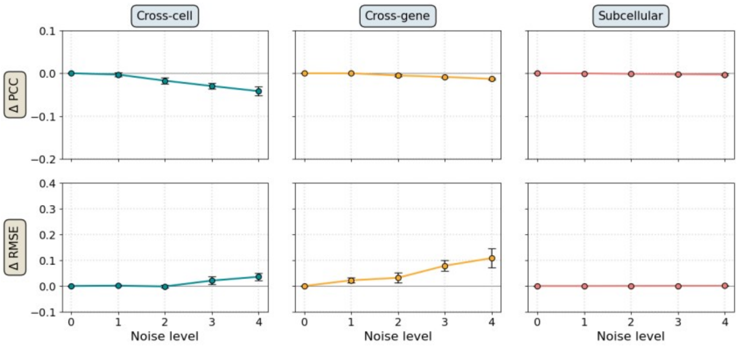

**c**

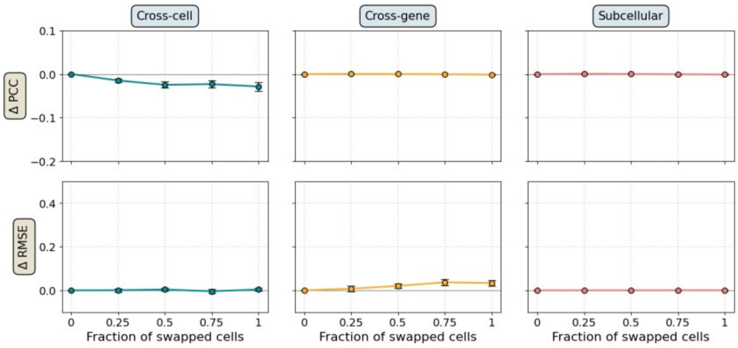

**Supplementary Figure 27. Expression levels of two cell-cycle markers in the MERFISH human osteosarcoma data.** Box plots show the expression distribution (log1p-transformed) of *CKAP5* (a G2/M phase marker) and *BRCA2* (an S/G2 phase marker) across clusters 0-3.

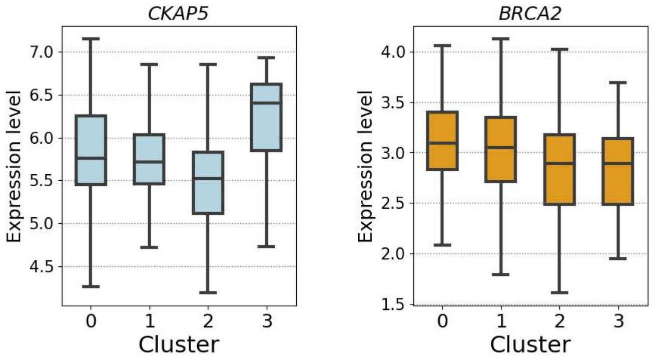

**Supplementary Figure 28. GO enrichment analysis of genes involved in cluster 3-specific co-localized gene pairs in the MERFISH human osteosarcoma dataset.** Bar plot shows the -log10(P) values of GO terms with *P* value < 0.01.

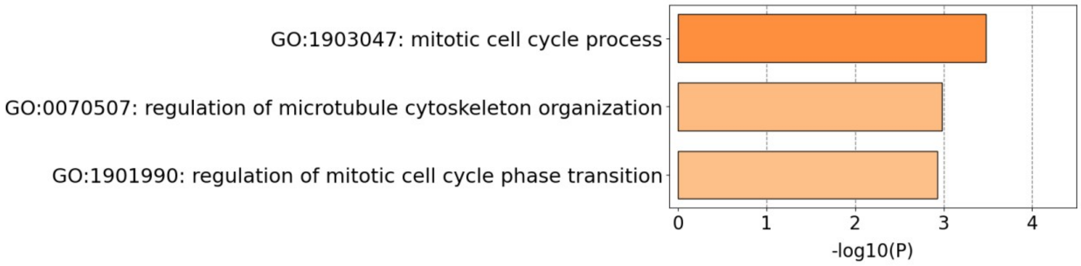

**Supplementary Figure 29. Comparison of the SVC-derived cell clusters with cell morphology in the MERFISH human osteosarcoma dataset. a.** Training cells projected onto a morphology-only feature space defined by the first two principal components of the concatenated cell and nuclear mask images, colored by their SVC-derived cluster labels. **b.** Contingency table comparing morphology-only clusters with SVC-derived cell clusters. Each entry indicates the number of cells assigned to the corresponding pair of clusters.

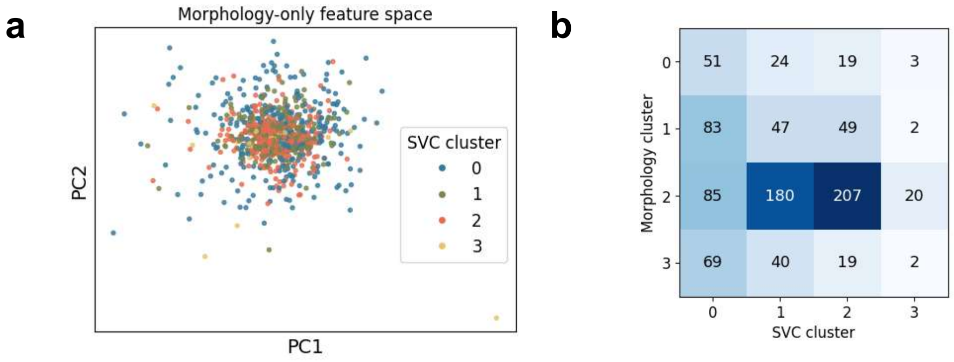

**Supplementary Figure 30. Contribution of cell morphology to the SVC-derived cell clusters in the MERFISH human osteosarcoma dataset. a.** Cells projected onto the first two linear discriminants of three representations, each fitted to separate the SVC-derived clusters: the SVC latent representation (left), the morphology-only representation (middle), and the morphology-residualized SVC latent representation (right). **b.** Cross-validated recovery of the SVC cluster labels from morphology alone and from the morphology-residualized latent. A class-balanced multinomial logistic-regression classifier was evaluated by five repeats of stratified five-fold cross-validation; bars give the mean and error bars the standard deviation across repeats. **c.** Cell area against cell eccentricity, one dot per cell, colored by SVC cluster.

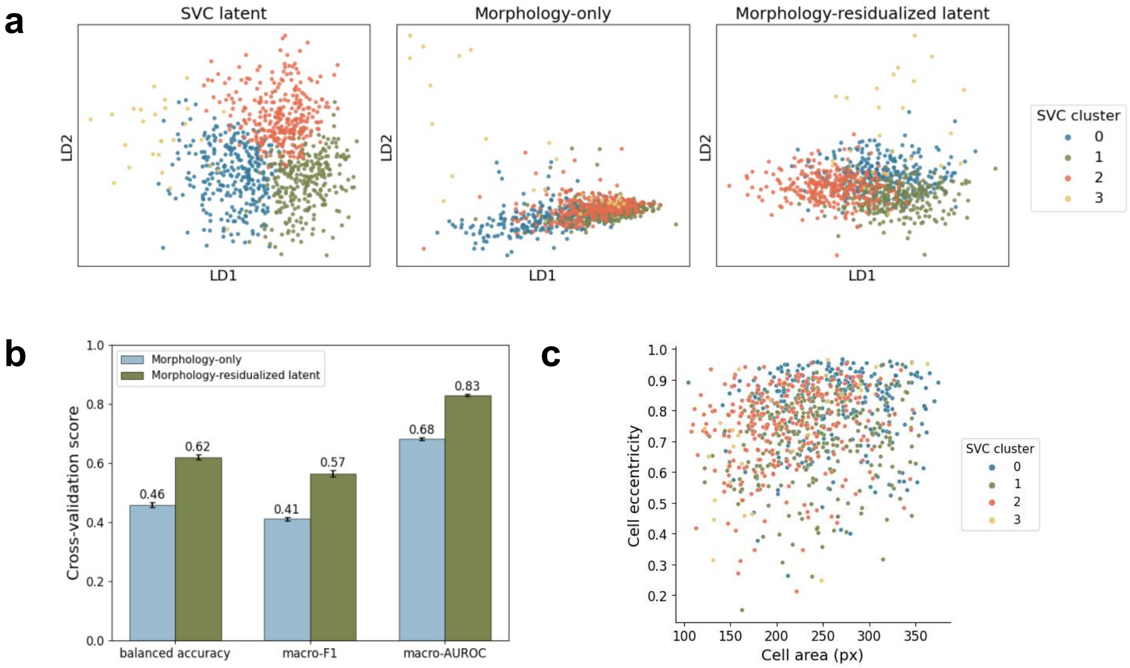

**Supplementary Figure 31. Spatial domains detected by SVC in Xenium mouse brain section 2.**

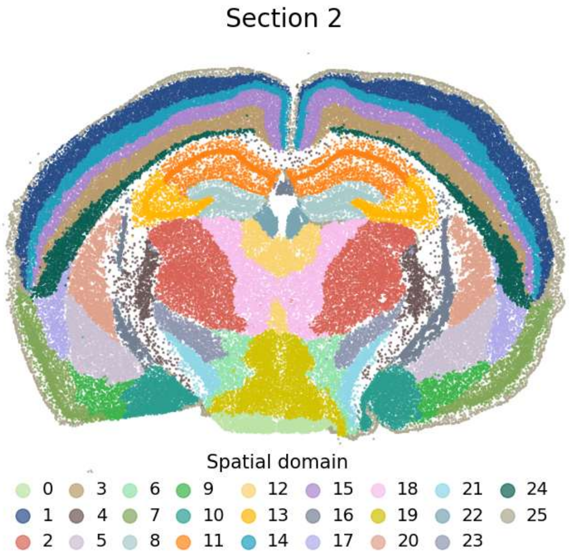

**Supplementary Figure 32. Gene subcellular pattern prediction performance of SVC and four baseline models in Xenium mouse brain. a.** Box plots summarize Pearson correlation (PCC), cosine similarity, and RMSE across genes on the held-out test data. For PCC and cosine similarity, higher values indicate better agreement; for RMSE, lower is better. **b.** Box plots show the per-gene performance gains of SVC over the four baseline models, evaluated using Pearson correlation coefficient (PCC), cosine similarity, and root mean squared error (RMSE), with positive values indicating better performance by SVC.

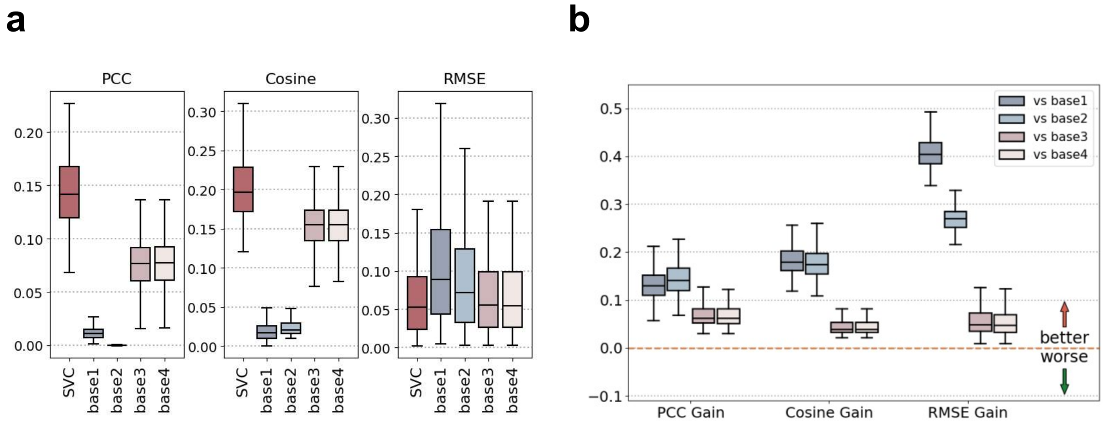

**Supplementary Figure 33. Cell-type-resolved prediction accuracy in the Xenium mouse brain data.** Predictions were compared with measurements separately within each major cell type, with each point representing one gene; Pearson correlation coefficients are shown in each panel. **a.** Scatter plots compare predicted versus observed expression levels (log1p-transformed) for each gene within the cell type; **b.** Predicted versus measured mean distance to the nuclear center for the same cell types.

**a**

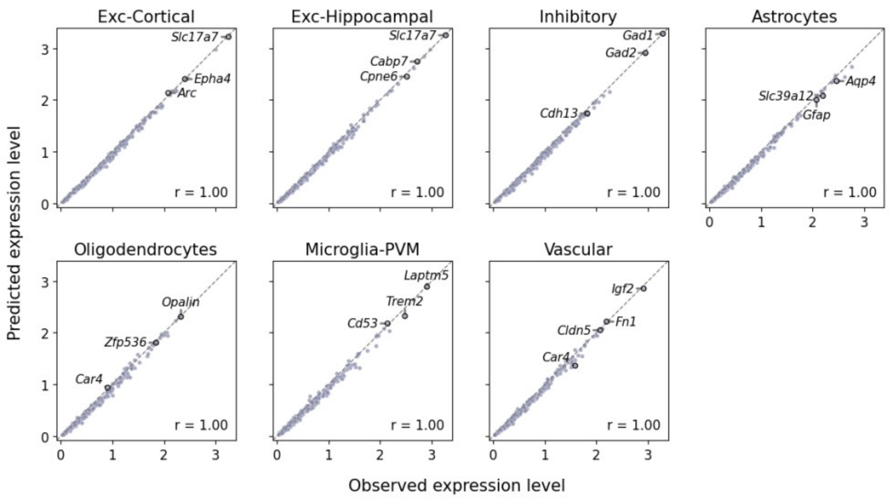

**b**

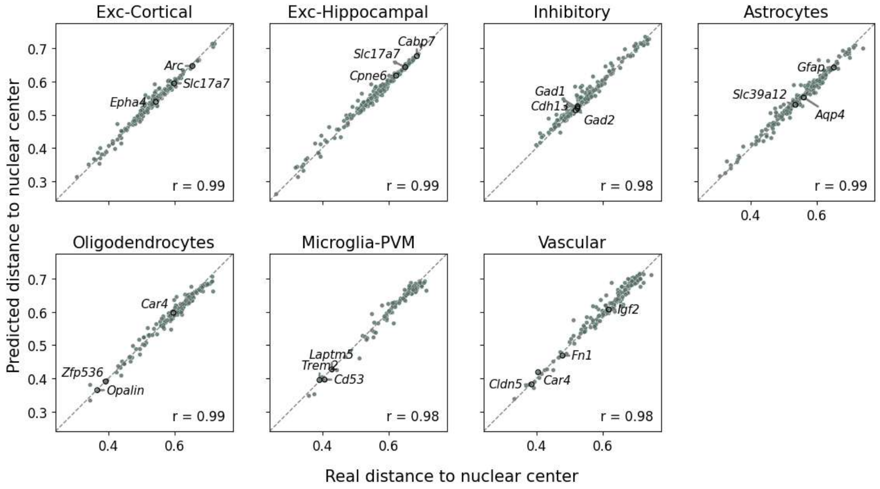

**Supplementary Figure 34. Subcellular transcript localization predictions in the Xenium mouse brain dataset.** Representative tissue regions showing the measured transcript distributions (top) and corresponding SVC-predicted distributions (bottom). Cells are colored according to major cell-type annotations, and transcript dots are colored by gene as indicated.

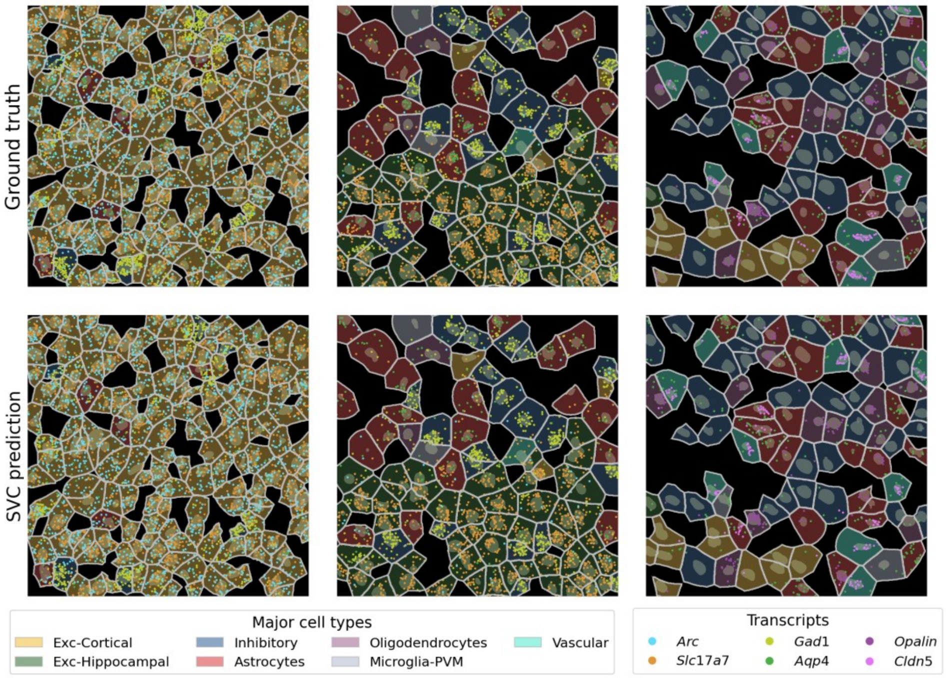

**Supplementary Figure 35. Cell-type-resolved comparison of SVC with the mean-distance baseline in the Xenium mouse brain data.** Predicted and measured mean distances to the nuclear center were compared separately within each of five cell types, with each point representing one gene; Pearson correlation coefficients are shown in each panel. Top row, predictions from SVC. Bottom row, predictions from baseline (train).

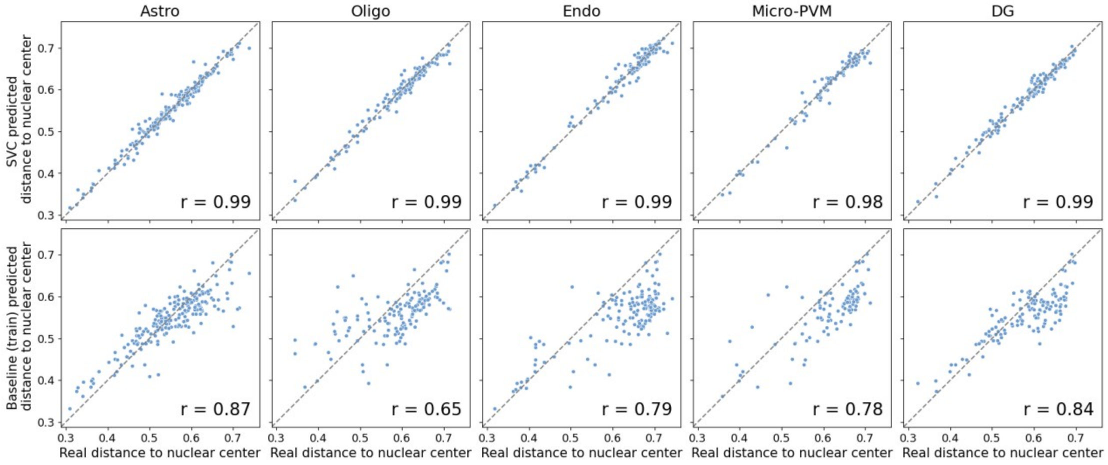

**Supplementary Figure 36. Cell-level gene expression prediction accuracy comparisons in the Xenium mouse brain data.** For the cell-level gene expression prediction task, box plots show the per-gene RMSE (log1p-transformed) and PCC scores between observed and predicted expression counts using 10 held-out gene folds. SVC is compared with uniPort, ENVI, Tangram, and gimVI.

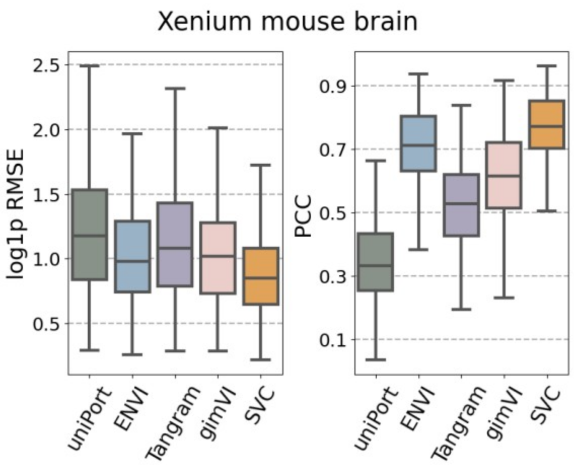

**Supplementary Figure 37. Retrieval of subcellular gene patterns from cell morphology in the Xenium mouse brain data. a.** ROC curves for retrieval, averaged over queries. A retrieved cell was counted as correct either when it was the query cell itself (self), or when it was among the five cells whose measured transcript patterns are most similar to that of the query (top 5 similar); the dashed line indicates chance. **b.** Representative retrieval examples for one query cell of each type. Each pair of rows shows the cell morphology and the corresponding transcript pattern summed over the genes used, for the query cell and for the five highest-ranked retrieved cells.

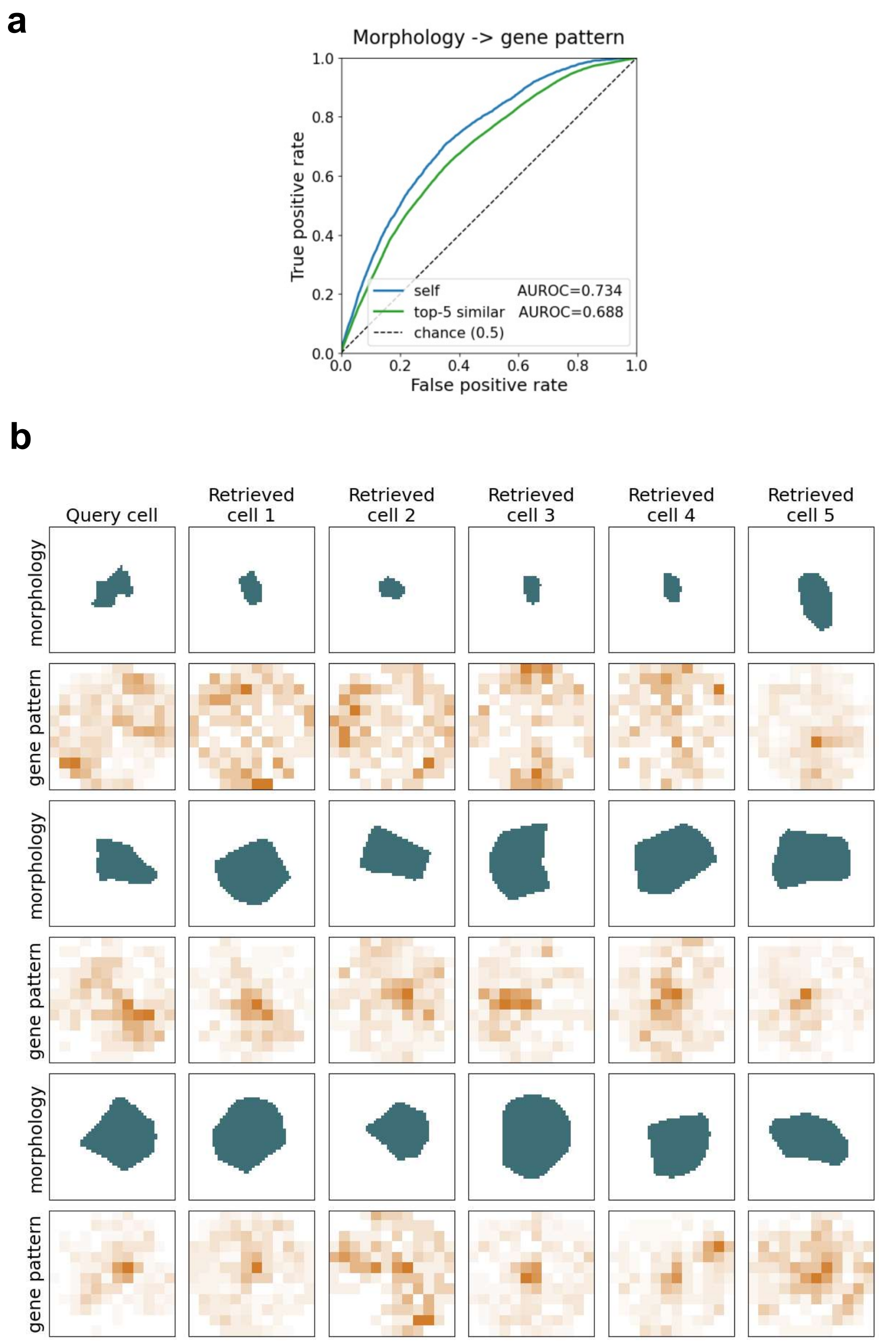

**Supplementary Figure 38. Retrieval of cell morphology and spatial context from subcellular gene patterns in the Xenium mouse brain data. a.** ROC curves for retrieval, averaged over queries, for retrieving a cell's morphology (left) and its spatial context, defined as the cell-type composition of its nearest neighbors (right). A retrieved cell was counted as correct either when it was the query cell itself (self), or when it was among the five cells with the most similar measured target, which includes the query cell itself (top 5 similar); the dashed line indicates chance. **b.** Representative retrieval examples for one query cell of each type, for the retrieval of morphology. Each pair of rows shows the transcript pattern summed over the genes used and the corresponding cell morphology, for the query cell and for the five highest-ranked retrieved cells.

a

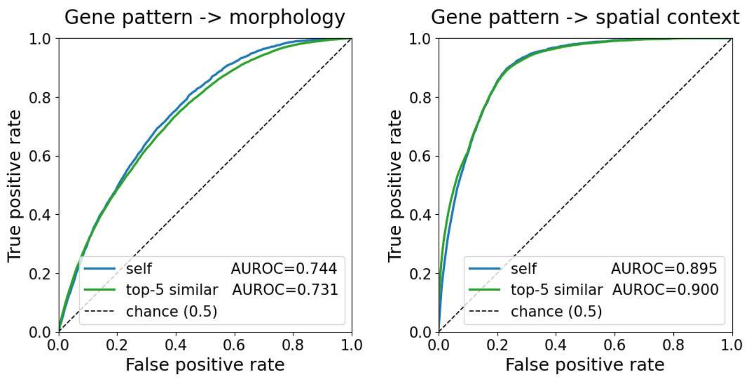

b

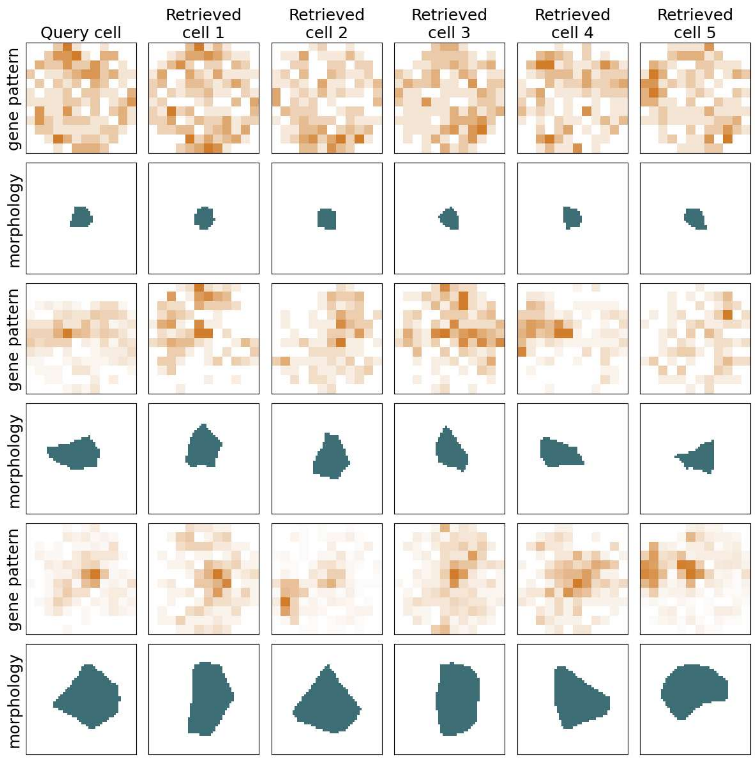

**Supplementary Figure 39. Spatial-shuffle permutation test in the Xenium mouse brain data.** Histograms showing the null distribution of the mean per-cell Earth Mover's Distance between the two genes' measured subcellular maps, over 1,000 rounds of a within-cell spatial shuffle, with the observed mean marked in orange; with one panel for the most type-specific co-localizing gene pair in each cell type, corresponding to the pairs shown in Fig. 4d.

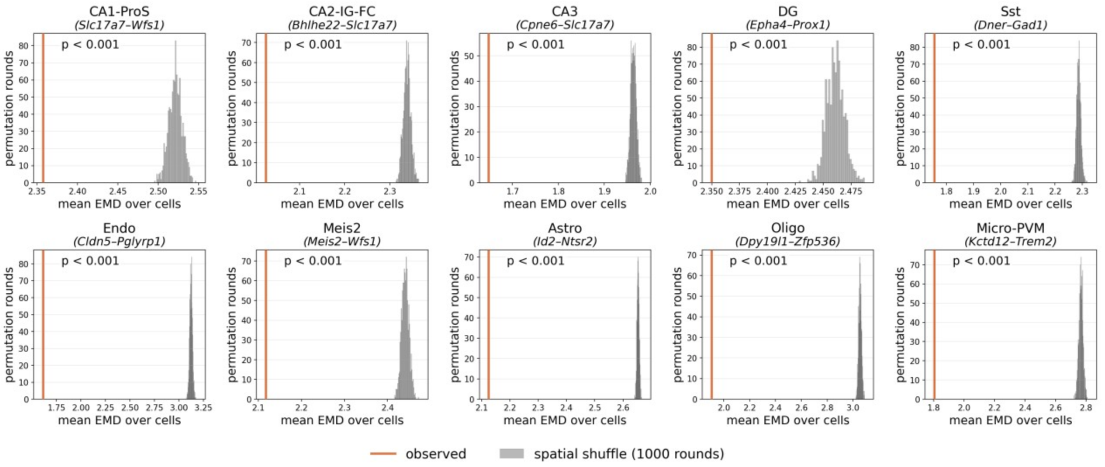

**Supplementary Figure 40. Relationship between subcellular co-localization and cell-level co-expression across ten cell types in the Xenium mouse brain data.** Each panel represents one cell type, with each point corresponding to one gene pair. Cell-level co-expression on the x axis was calculated as the Pearson correlation between the two genes' size-normalized, log1p-transformed expression across cells, whereas the subcellular co-localization score is shown on the y axis. Gene pairs are colored according to whether they rank in the top 1% for co-localization only, co-expression only, both, or neither within each cell type. Stars indicate the most type-specific gene pair of each cell type, shown in Fig. 4d. The Pearson correlation between co-localization and co-expression scores is indicated in each panel.

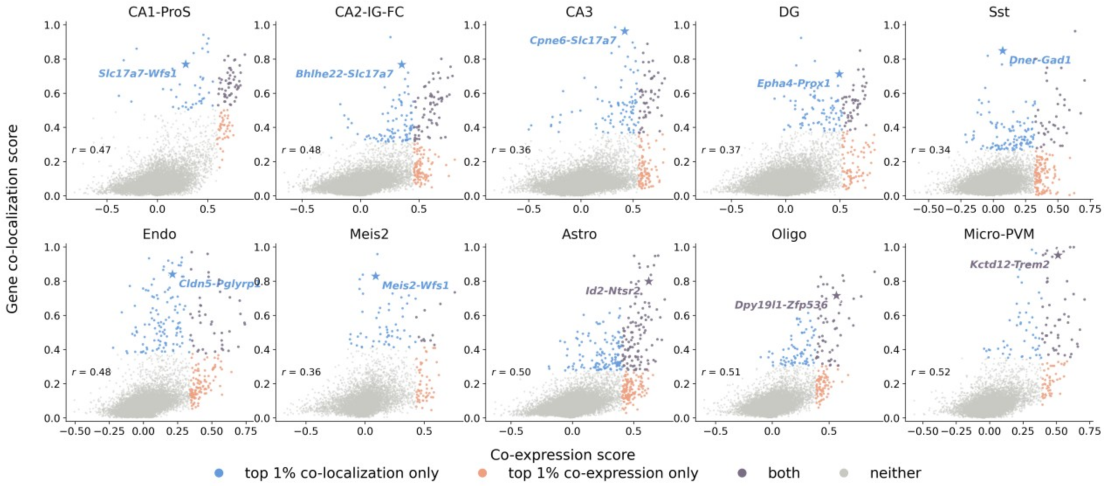

**Supplementary Figure 41. Gene-level principal components of SVC latent representations in VLMCs from the Xenium mouse brain dataset.** Within each cell, gene tokens were centered by subtracting their mean across the full gene panel, thereby removing the cell-level component shared across genes. Each gene was then represented by its mean centered token across cells. Principal component analysis was then performed on the resulting gene-level representations after residualizing them with respect to the logarithms of each gene's median and total transcript counts across cells.

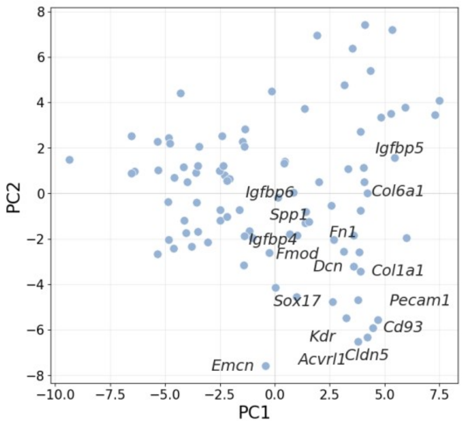

**Supplementary Figure 42. Spatial distribution of the difference in module distance to the nuclear center and of *Opalin* expression in oligodendrocytes across the two Xenium mouse brain sections.** Top: each oligodendrocyte detecting all six genes of the two modules (OPC: *Pdgfra*, *Cspg4*, *Gpr17*; maturation-associated: *Clmn*, *Zfp536*, *Prox1*), colored by the per-cell difference in distance to the nuclear center between the two modules (maturation-associated minus OPC). Bottom: the same cells colored by their *Opalin* transcript count on a log scale. Gray marks all remaining cells of each section.

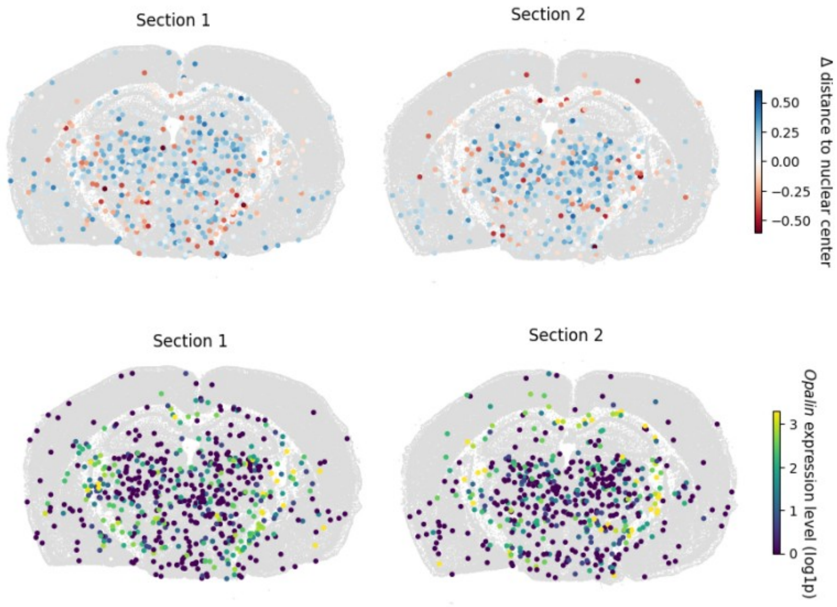

**Supplementary Figure 43. Expression of the two oligodendrocyte modules along the *Opalin* maturation axis in Xenium mouse brain data.** Average expression level of each module, summed over its three genes in all oligodendrocytes, binned by *Opalin* expression level.

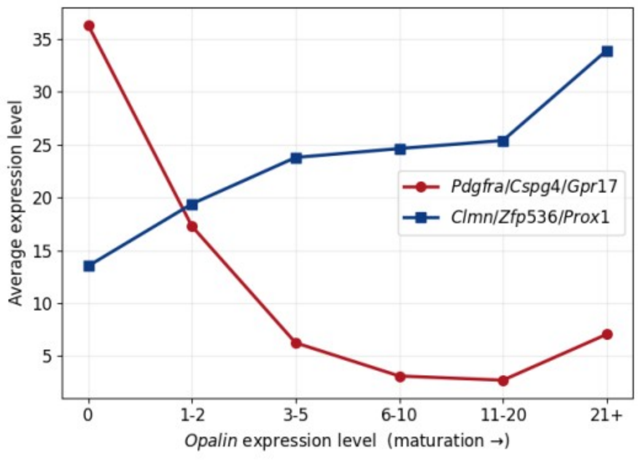

**Supplementary Figure 44. Evaluation of the cross-dataset transfer ability of SVC trained on Xenium mouse brain data section 1.** Gene subcellular pattern prediction performance comparison between SVC and four baseline models. **a.** Distribution-based performance metrics (EMD, KL and JS) for within-dataset evaluation from Xenium mouse brain section 1 to section 2. **b.** Pearson correlation, cosine similarity and RMSE for the same within-dataset evaluation. **c.** Distribution-based performance metrics for cross-dataset transfer from Xenium mouse brain data to MERFISH mouse brain data without retraining. **d.** Pearson correlation, cosine similarity and RMSE for the same cross-dataset transfer. Positive values indicate better performance.

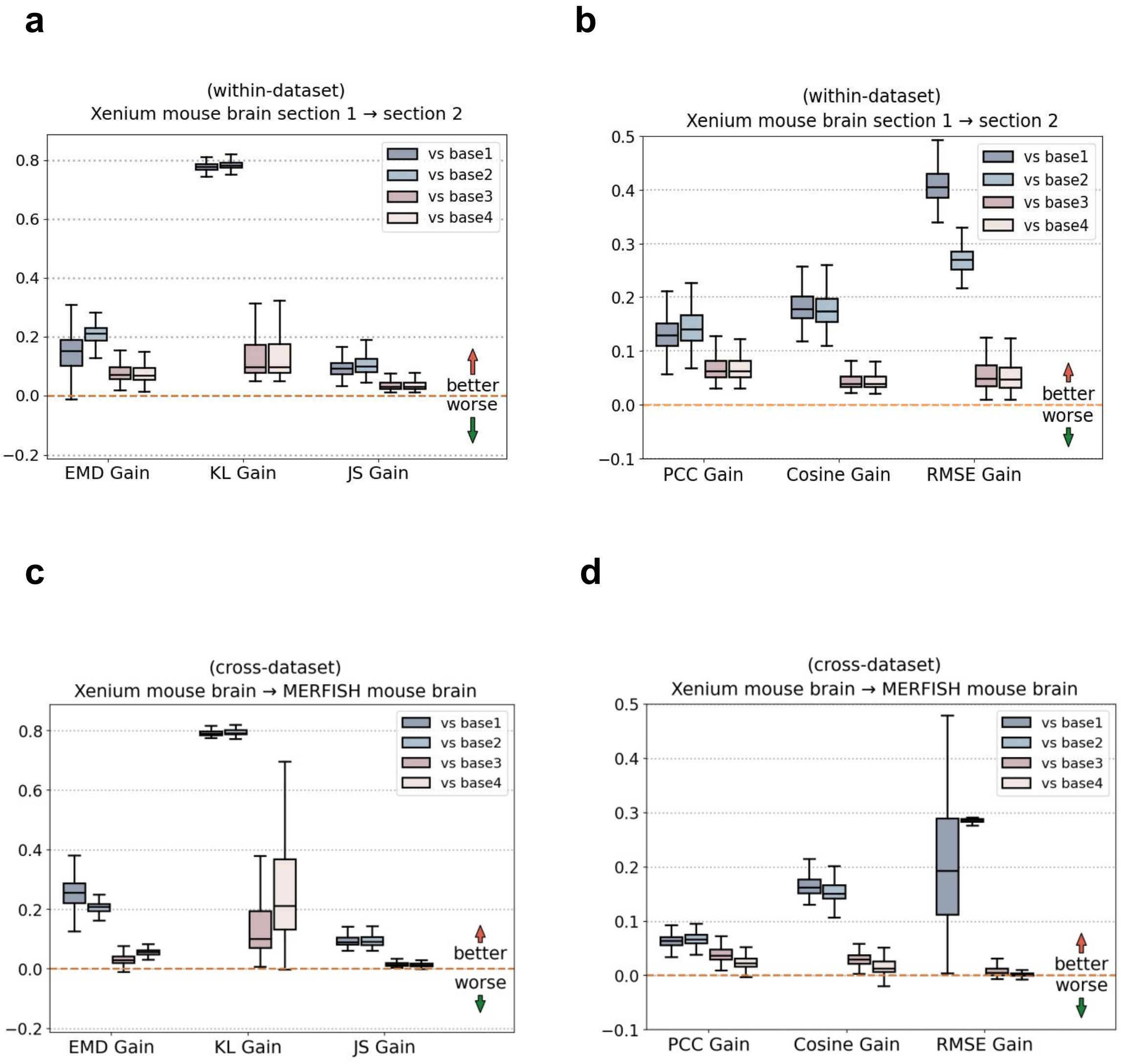

**Supplementary Figure 45. Subcellular transcript localization predictions in MERFISH mouse brain data under cross-dataset transfer.** Top: the whole section with cells colored by major cell type and the three regions outlined. Below: for each region, the measured transcript distributions (upper row) and the corresponding SVC-predicted distributions (lower row), from the model trained on Xenium mouse brain data and applied without retraining. Cells are colored according to major cell-type annotations, and transcript dots are colored by gene as indicated.

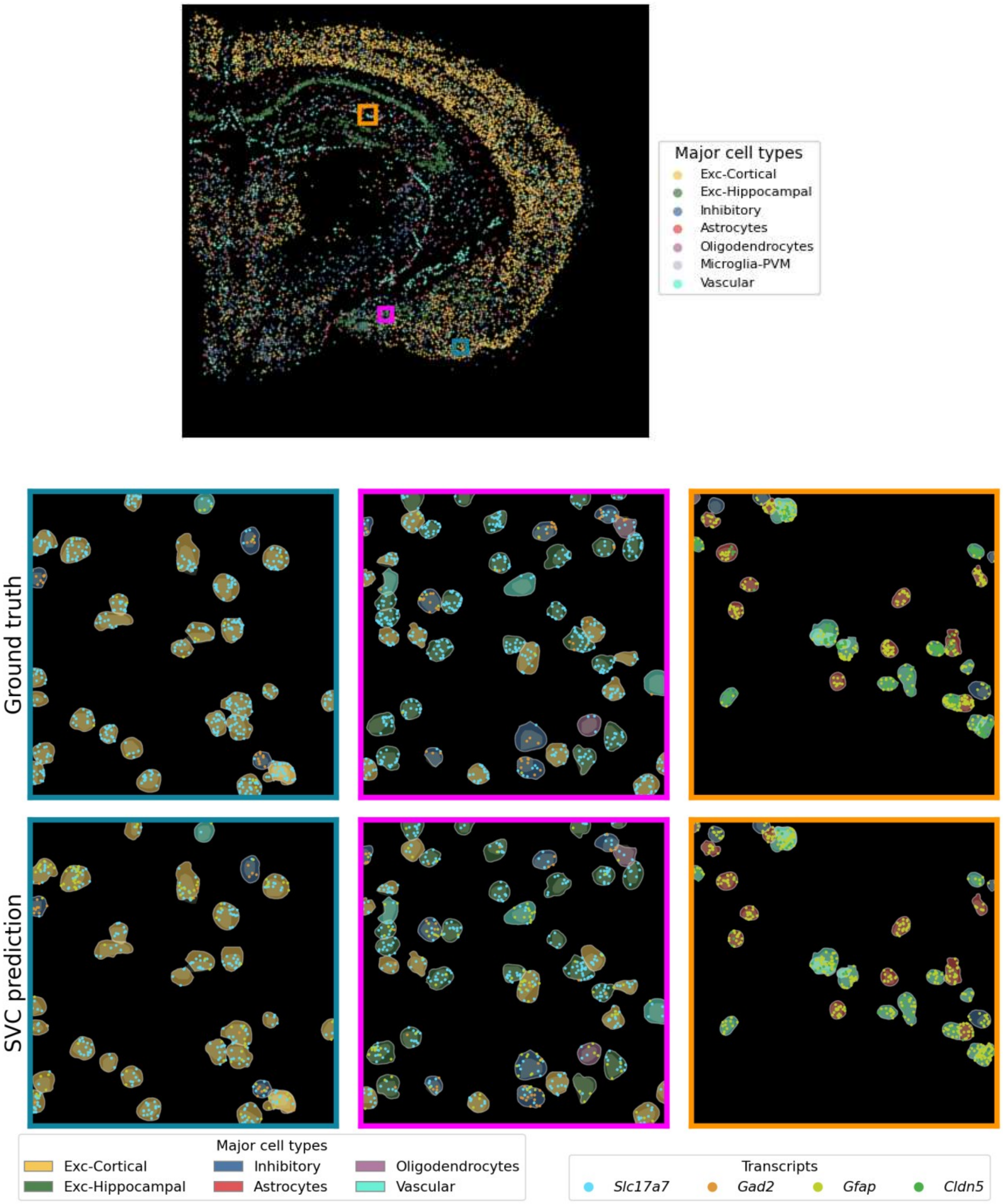

**Supplementary Figure 46. Evaluation of the cross-tissue transfer ability of SVC trained on Xenium breast cancer data section 1.** Gene subcellular pattern prediction performance comparison between SVC and four baseline models. **a.** Distribution-based performance metrics (EMD, KL and JS) for within-tissue evaluation from Xenium breast cancer section 1 to section 2. **b.** Pearson correlation, cosine similarity and RMSE for the same within-tissue evaluation. **c.** Distribution-based performance metrics for cross-tissue transfer from Xenium breast cancer data to Xenium lung cancer data without retraining. **d.** Pearson correlation, cosine similarity and RMSE for the same cross-tissue transfer. Positive values indicate better performance.

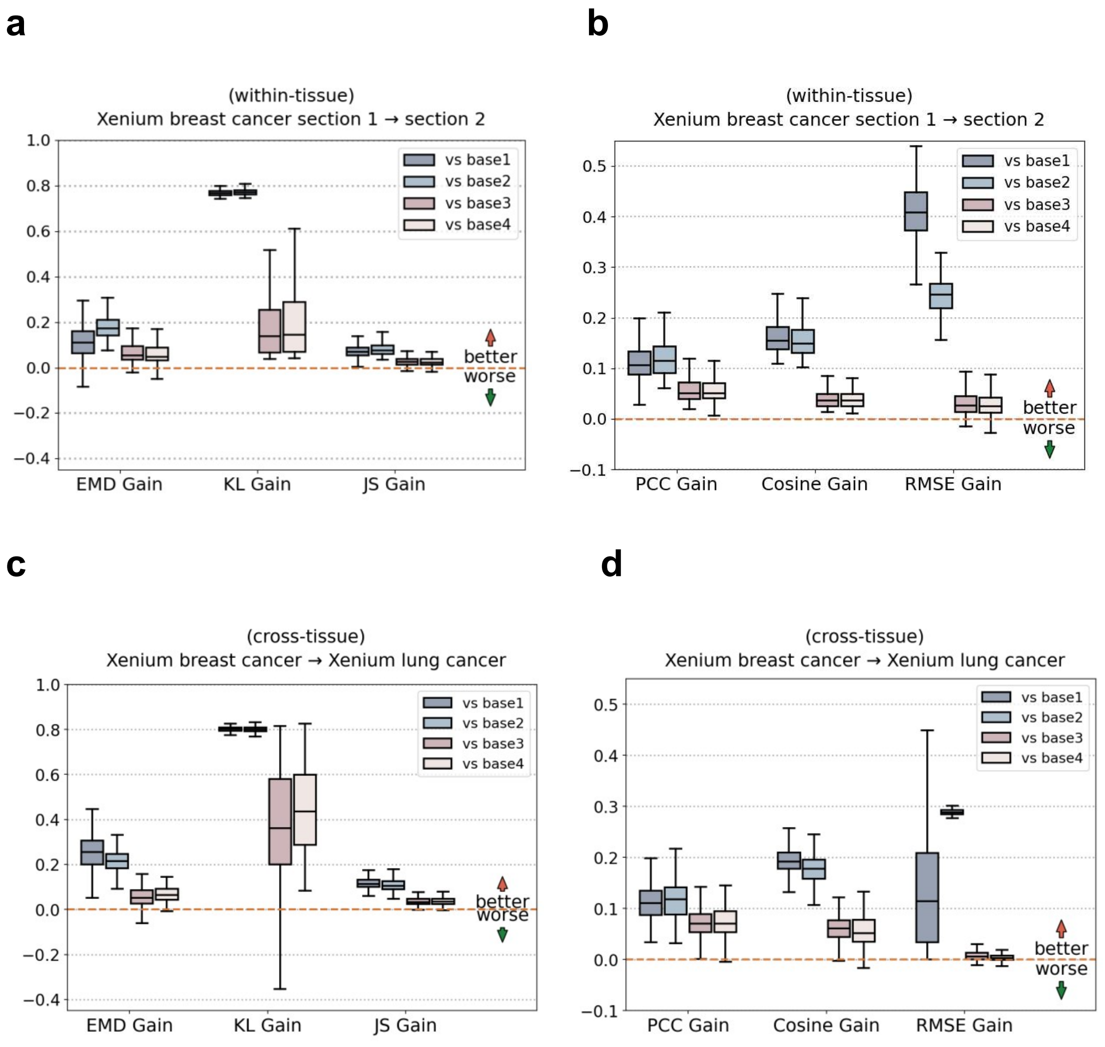

**Supplementary Figure 47. Subcellular transcript localization predictions in Xenium lung cancer data under cross-tissue transfer.** Top: the whole section with cells colored by cluster and the three regions outlined. Below: for each region, the measured transcript distributions (upper row) and the corresponding SVC-predicted distributions (lower row), from the model trained on Xenium breast cancer data and applied without retraining. Cells are colored by the lung sample's graph clusters, and transcript dots are colored by gene as indicated.

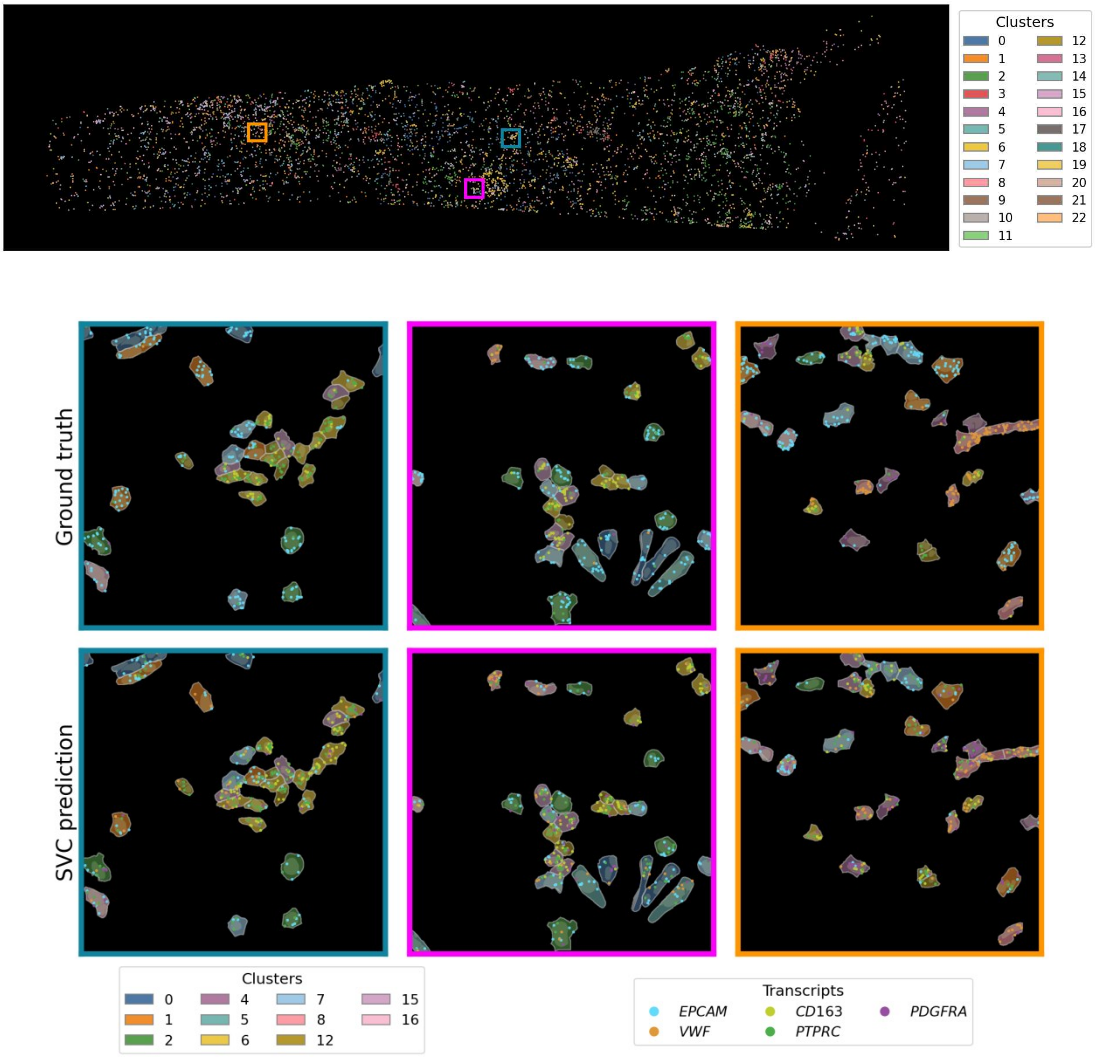

**Supplementary Figure 48. Imputation of unmeasured marker genes in the Xenium Prime 5K mouse brain hemisphere dataset.** Spatial maps show SVC-predicted cell-level expression for unmeasured marker genes *Rasgrf2*, *Rprm*, *Wfs1*, and *Prss35*.

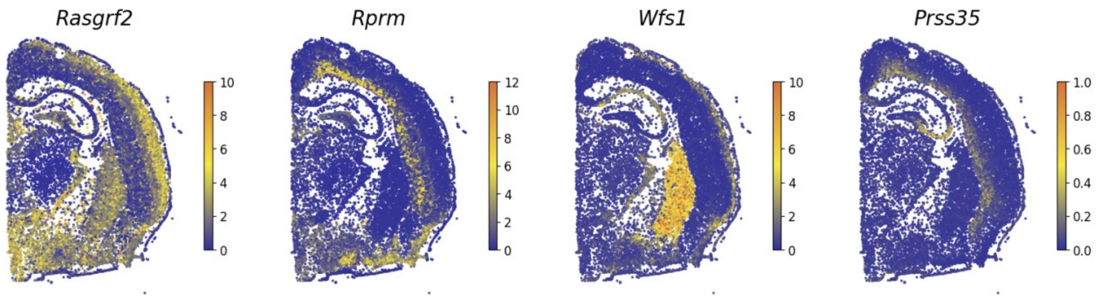

**Supplementary Figure 49. Measured per-cell polarity of three secreted genes in VLMCs of the Xenium mouse brain dataset.** Box plots showing the per-cell polarity of *Igfbp6*, *Spp1* and *Igfbp5*, the three genes in VLMCs of the Xenium mouse brain sections. Each gene was compared with a count-matched background constructed from the other genes of the panel by resampling to match the target gene's per-cell transcript-count distribution. Statistical significance (n.s. ≥ 0.05; * < 0.05; ** < 0.01; *** < 0.001) was assessed using one-sided Mann-Whitney U tests, with Benjamini-Hochberg correction across genes.

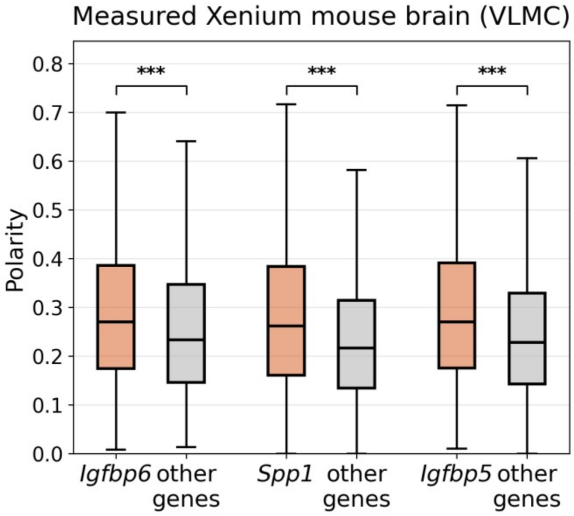

**Supplementary Figure 50. Predicted *Spp1* transcripts in representative VLMCs of the Xenium Prime 5K mouse brain hemisphere dataset.** Green dots represent predicted *Spp1* transcripts, and purple dots represent the pooled predicted transcripts of the other imputed genes. For each transcript set, the shaded ellipse represents the 1-standard-deviation contour of a fitted bivariate Gaussian distribution, the cross marks the transcript center, and the connecting segment shows its displacement from the nuclear center.

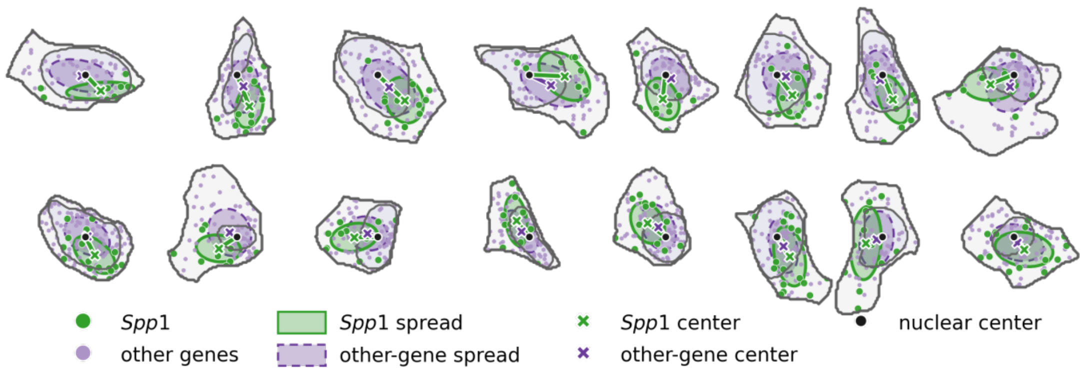

**Supplementary Figure 51. Co-localization of the two gene pairs measured in the Xenium mouse brain dataset.** Box plots showing the per-cell Earth Mover's Distance between the measured subcellular distributions of each gene pair in astrocytes (Astro, left) and endothelial cells (Endo, right) of the Xenium mouse brain sections. Each pair was compared with a count-matched background of the same gene paired with every other gene, resampled to match the pair's per-cell total transcript count. Statistical significance (n.s. ≥ 0.05; * < 0.05; ** < 0.01; *** < 0.001) was assessed using one-sided Mann-Whitney U tests, with Benjamini-Hochberg correction across gene pairs.

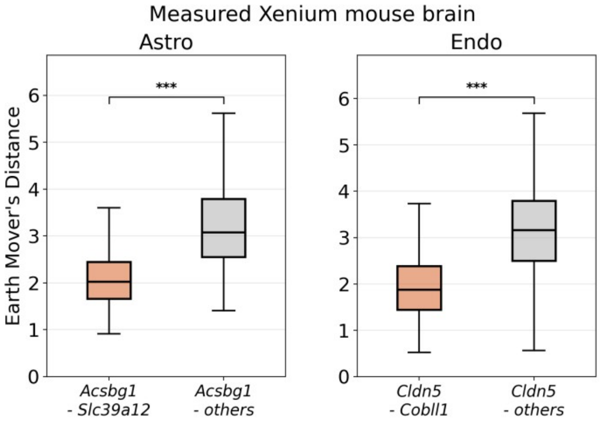

**Supplementary Figure 52. Spatial-shuffle permutation test for the two gene pairs in the Xenium 5K imputation and in measured Xenium mouse brain data.** Histograms showing the null distribution of the mean per-cell Earth Mover's Distance between the two genes' subcellular maps, over 1,000 rounds of a within-cell spatial shuffle, with the observed mean marked in orange; one panel per cell type for each of the two pairs shown in Fig. 4j, on the Xenium 5K imputation (left) and the measured Xenium mouse brain sections (right).

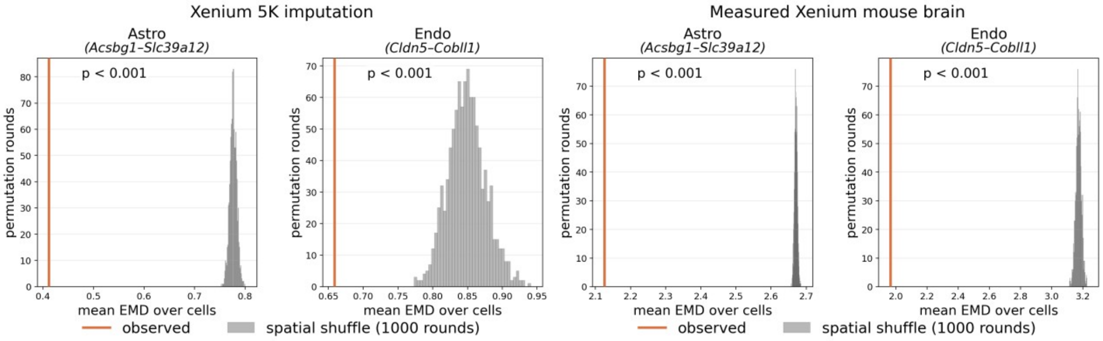

**Supplementary Figure 53. Spatial distributions of astrocytes and Micro-PVM cells in wild-type and P301S tau-transgenic sections in Xenium Prime 5K P301S tauopathy mouse brain data.** Cell positions are colored by cell type. Dashed boxes indicate the hippocampal regions, which are enlarged below.

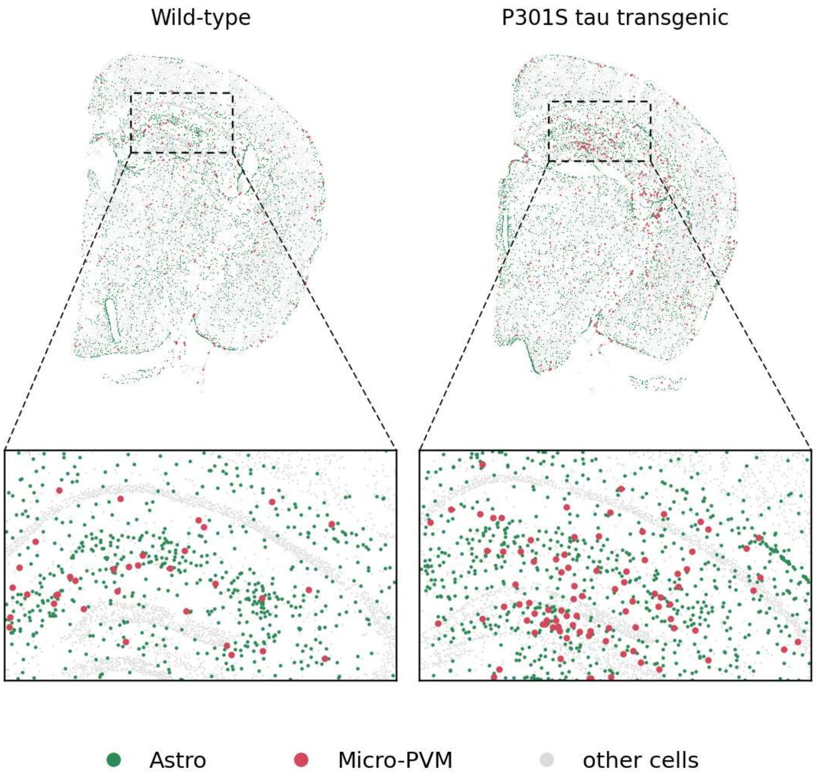

**Supplementary Figure 54. Relationship between predicted expression and subcellular localization changes for imputed genes in Xenium Prime 5K P301S tauopathy mouse brain data.** Each point represents one imputed gene in one cell type that met the detection threshold (5 genes in Micro-PVM cells and 11 genes in astrocytes). The x axis shows the log2 fold change in mean predicted transcript count between tau Tg and WT cells, and the y axis shows the corresponding change in median distance to the nuclear center (tau Tg - WT). Genes with significant localization shifts (FDR < 0.05) are outlined in black and labeled. The Pearson correlation between expression and localization changes is indicated in the plot.

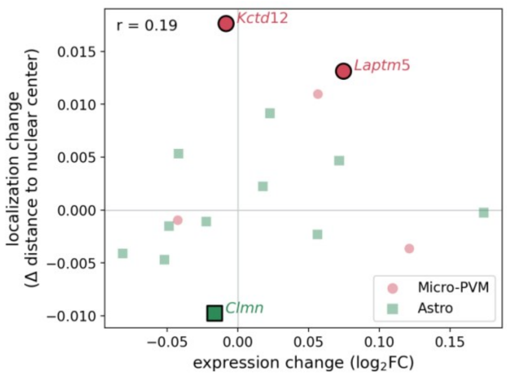

**Supplementary Figure 55. Gene subcellular pattern prediction performance of SVC and four baseline models in Xenium breast cancer. a.** Box plots summarize Pearson correlation (PCC), cosine similarity, and RMSE across genes on the held-out test data. For PCC and cosine similarity, higher values indicate better agreement; for RMSE, lower is better. **b.** Box plots show the per-gene performance gains of SVC over the four baseline models, evaluated using Pearson correlation coefficient (PCC), cosine similarity, and root mean squared error (RMSE), with positive values indicating better performance by SVC.

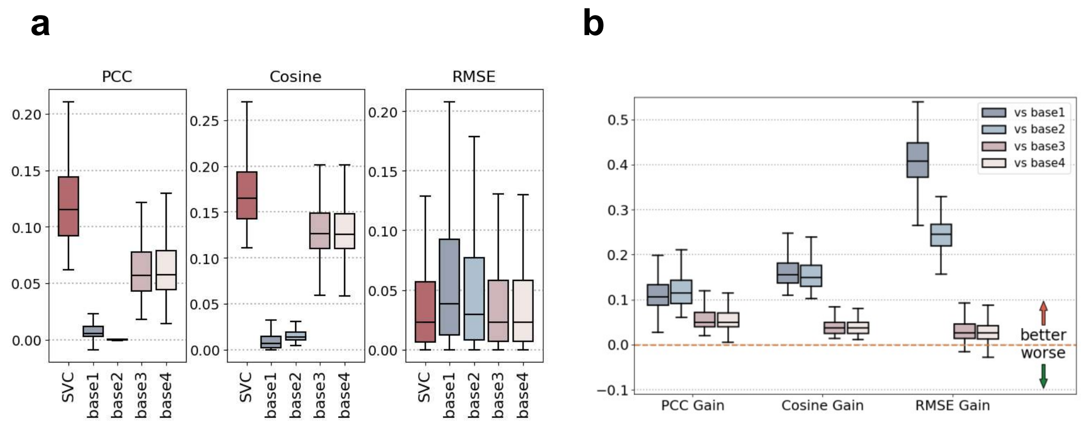

**Supplementary Figure 56. Cell-type-resolved prediction accuracy in the Xenium breast cancer data.** Predictions were compared with measurements separately within each major cell type, with each point representing one gene; Pearson correlation coefficients are shown in each panel. **a.** Scatter plots compare predicted versus observed expression levels (log1p-transformed) for each gene within the cell type; **b.** Predicted versus measured mean distance to the nuclear center for the same cell types.

**a**

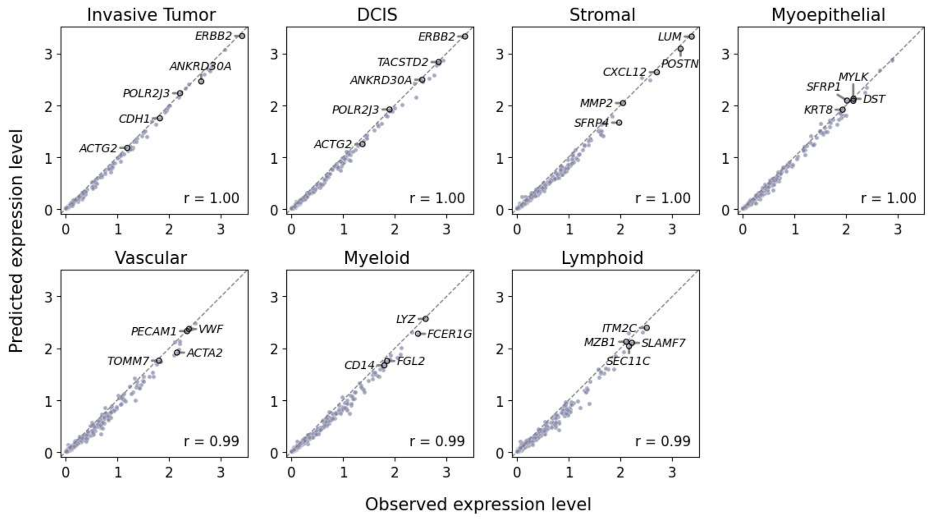

**b**

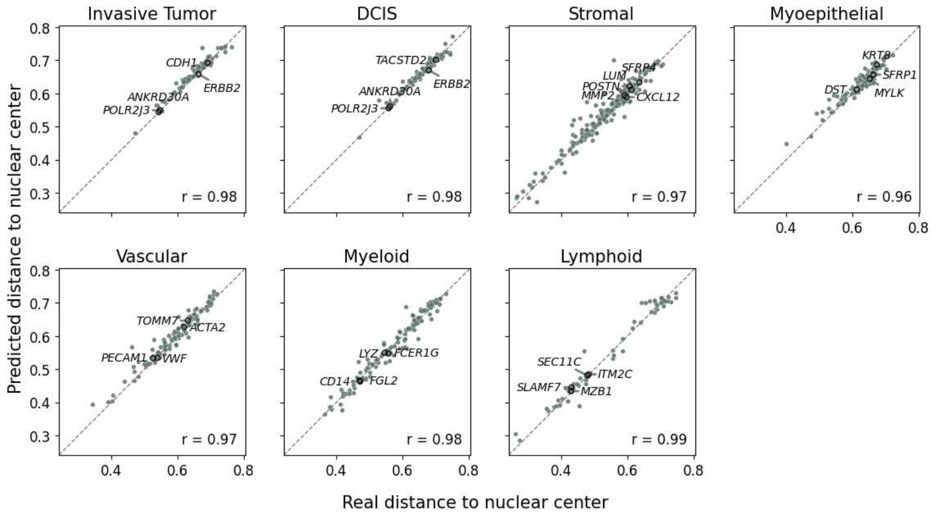

**Supplementary Figure 57. Subcellular transcript localization predictions in the Xenium breast cancer dataset.** Representative tissue regions showing the measured transcript distributions (top) and corresponding SVC-predicted distributions (bottom). Cells are colored according to major cell-type annotations, and transcript dots are colored by gene as indicated.

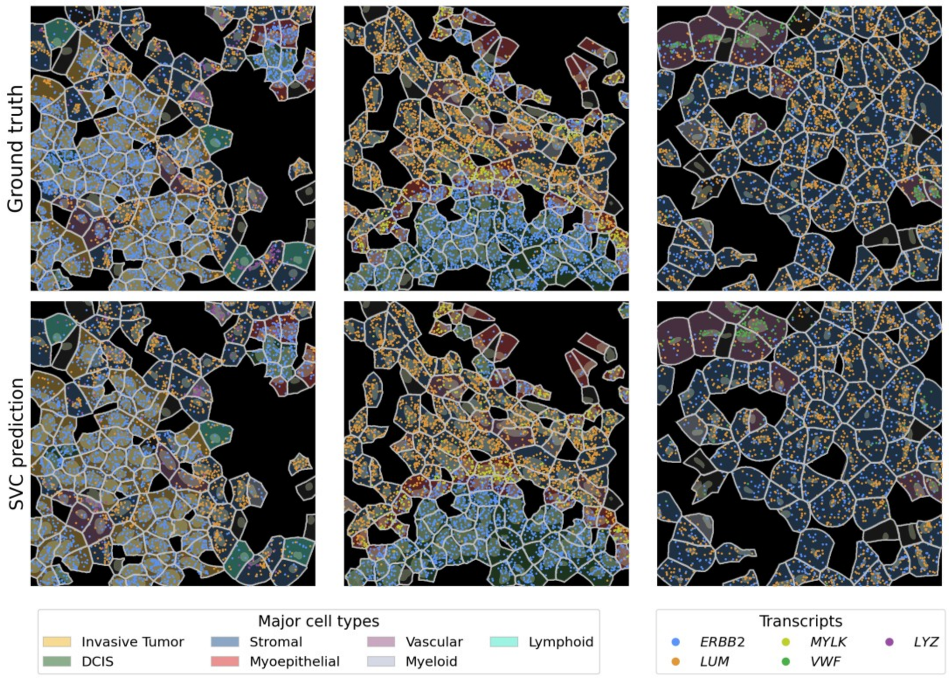

**Supplementary Figure 58. Cell-level gene expression prediction accuracy comparisons in Xenium breast cancer.** For the cell-level gene expression prediction task, box plots show the per-gene RMSE (log1p-transformed) and PCC scores between observed and predicted expression counts using 10 held-out gene folds. SVC is compared with uniPort, ENVI, Tangram, and gimVI.

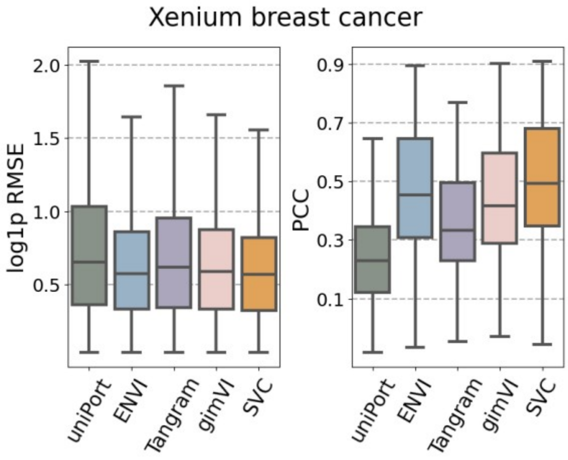

**Supplementary Figure 59. Generalization of SVC across anatomically distinct tissue regions in the Xenium breast cancer dataset.** SVC was retrained from scratch using only the cells on one side of a horizontal cut through the section, with the cells on the other side held out entirely. The trained model was then applied to all cells of the section, and spatial domains were detected on the whole section. **a.** Training on the upper region, with the lower region held out. **b.** The complementary split, training on the lower region.

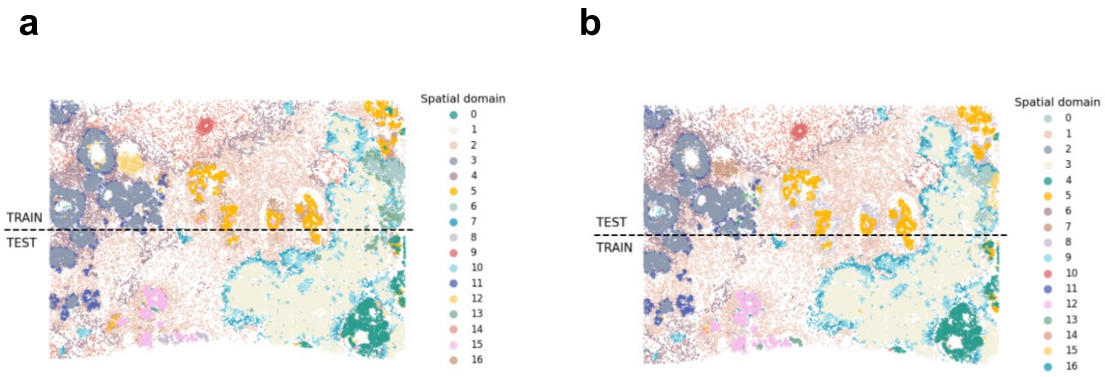

**Supplementary Figure 60. Generalization of SVC subcellular prediction across anatomically distinct regions in the Xenium breast cancer dataset. a.** Subcellular prediction performance of SVC and the four baselines when SVC was trained on the upper 50% of the tissue section and evaluated on the held-out lower 50%, assessed using PCC, cosine similarity, and RMSE. **b.** Corresponding performance gains of SVC over each baseline. **c-d.** Same analyses with the training and test regions reversed, using the lower 50% for training and the upper 50% for evaluation. Positive gain values indicate better performance by SVC.

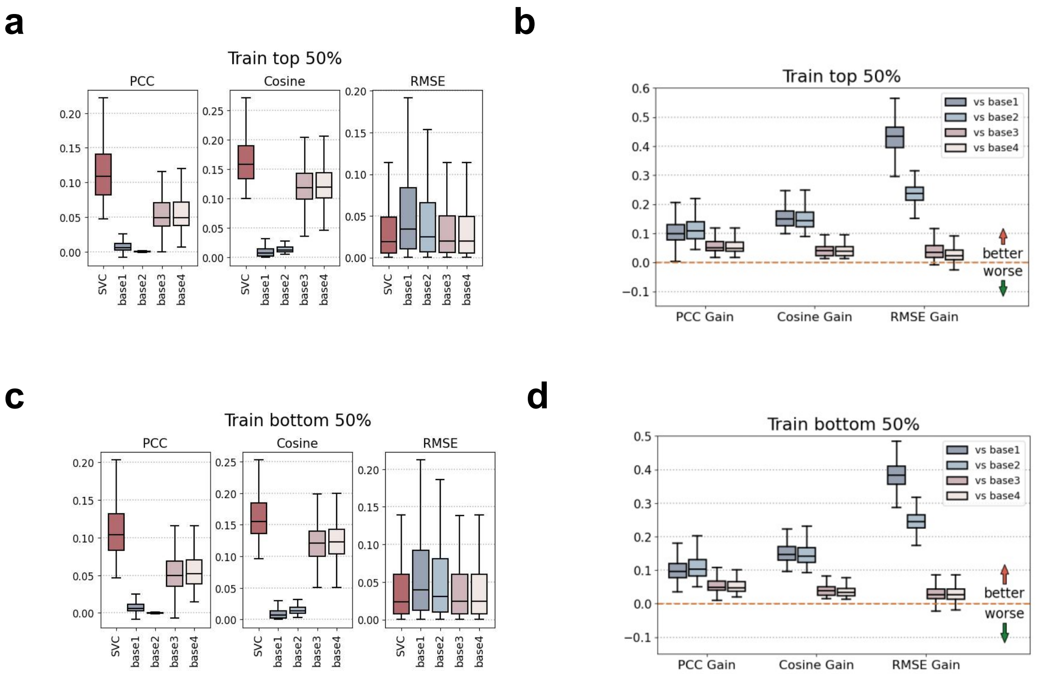

**Supplementary Figure 61. Cell-type-specific out-of-distribution prediction of nuclear-proximity patterns in the Xenium breast cancer dataset.** SVC was retrained from scratch with B cells removed entirely from the training data. **a.** Genes selected for OOD evaluation, whose mean distance to the nuclear center in held-out B cells falls outside the range observed across all other expressing cell types. Stars indicate B cells and gray dots indicate the other cell types. **b.** Predicted versus measured mean distances to the nuclear center for the selected genes in held-out B cells, comparing SVC with Baseline 3, which uses the corresponding gene-specific mean distance from the training cells. **c.** Accuracy of the predicted subcellular maps of these genes in the held-out B cells, measured by PCC, cosine similarity and RMSE against the four baselines; bars show the mean across genes and dots the individual genes.

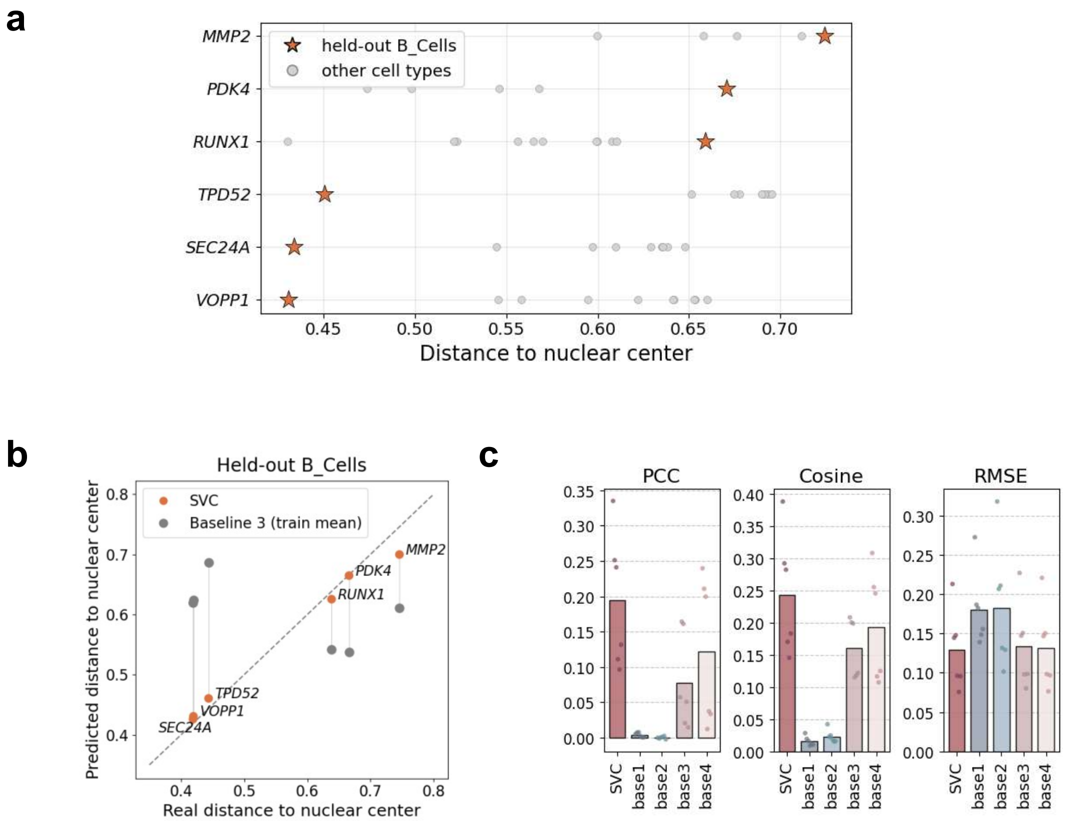

**Supplementary Figure 62. Cell-type-specific out-of-distribution prediction of transcript-polarity patterns in the Xenium breast cancer dataset.** SVC was retrained from scratch with ACTA2+ myoepithelial cells entirely excluded from the training data. **a.** Genes selected for OOD evaluation, whose mean polarity in held-out ACTA2+ myoepithelial cells falls outside the range observed across all other expressing cell types. Stars indicate held-out ACTA2+ myoepithelial cells and gray dots indicate the other cell types. **b.** Measured and predicted mean polarity of the selected genes in held-out ACTA2+ myoepithelial cells, comparing SVC with Baseline 3, using the polarity of the gene's mean map over the training cells. **c.** Subcellular localization prediction performance for these genes in held-out ACTA2+ myoepithelial cells, evaluated using PCC, cosine similarity, and RMSE against the four baselines. Bars show the mean across genes and dots indicate individual genes.

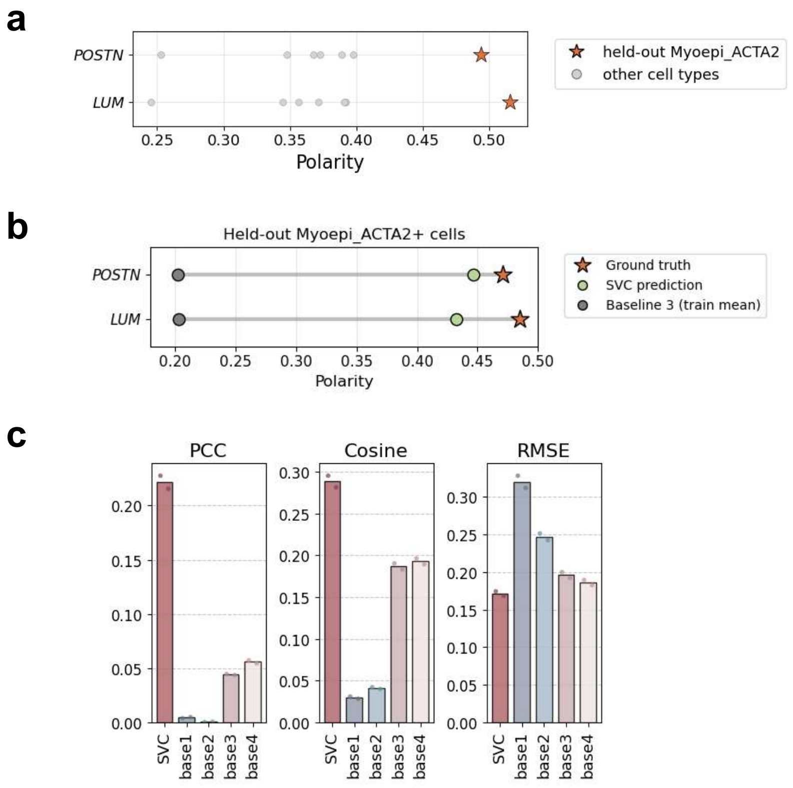

**Supplementary Figure 63. Subcellular localization predictions for cell-type-specific OOD polarity genes in the Xenium breast cancer dataset.** A representative tissue region showing the measured transcript distributions (left) and corresponding SVC predictions (right) for *LUM* and *POSTN* in held-out ACTA2+ myoepithelial cells. Cells are colored according to cell-type annotations, and transcript dots are colored by gene as indicated.

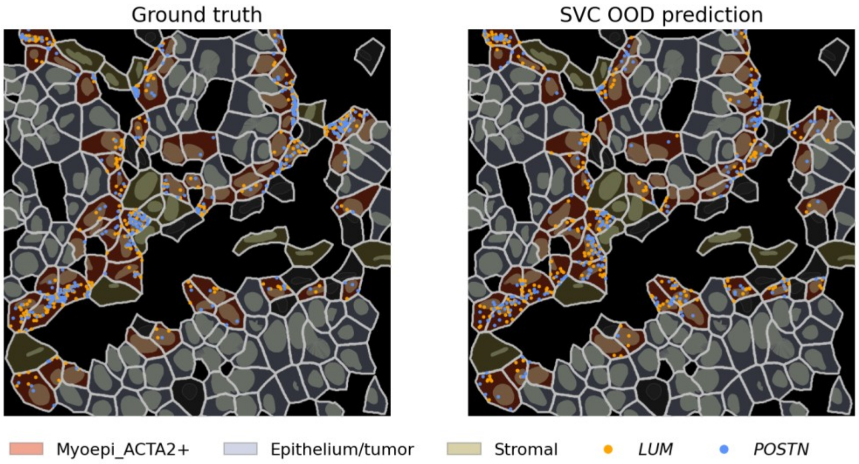

**Supplementary Figure 64. Spatial-shuffle permutation test in Xenium breast cancer data.** Histograms showing the null distribution of the mean per-cell Earth Mover's Distance between the two genes' measured subcellular maps, over 1,000 rounds of a within-cell spatial shuffle, with the observed mean marked in orange; with one panel for the most type-specific co-localizing gene pair in each cell type, corresponding to the pairs shown in Fig. 5d.

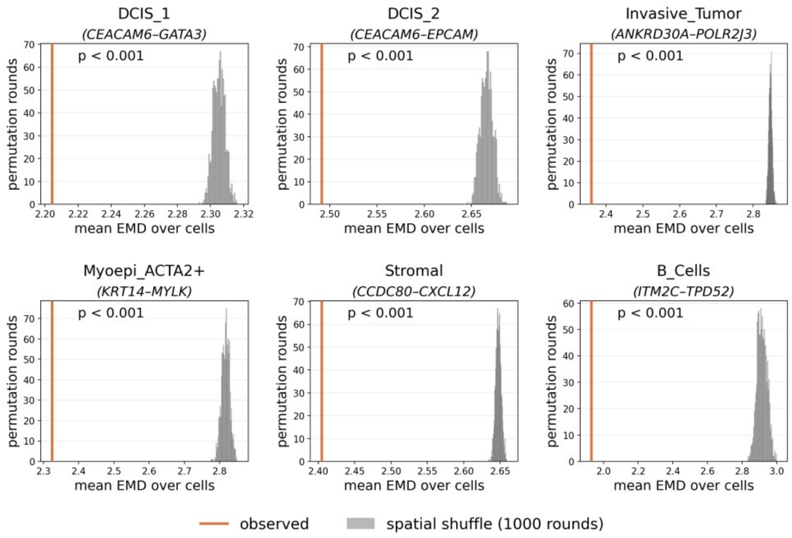

**Supplementary Figure 65. Relationship between subcellular co-localization and cell-level co-expression across six cell types in Xenium breast cancer data.** Each panel represents one cell type, with each point corresponding to one gene pair. Cell-level co-expression on the x axis was calculated as the Pearson correlation between the two genes' size-normalized, log1p-transformed expression across cells, whereas the subcellular co-localization score is shown on the y axis. Gene pairs are colored according to whether they rank in the top 1% for co-localization only, co-expression only, both, or neither within each cell type. Stars indicate the most type-specific gene pair of each cell type, shown in Fig. 5d. The Pearson correlation between co-localization and co-expression scores is indicated in each panel.

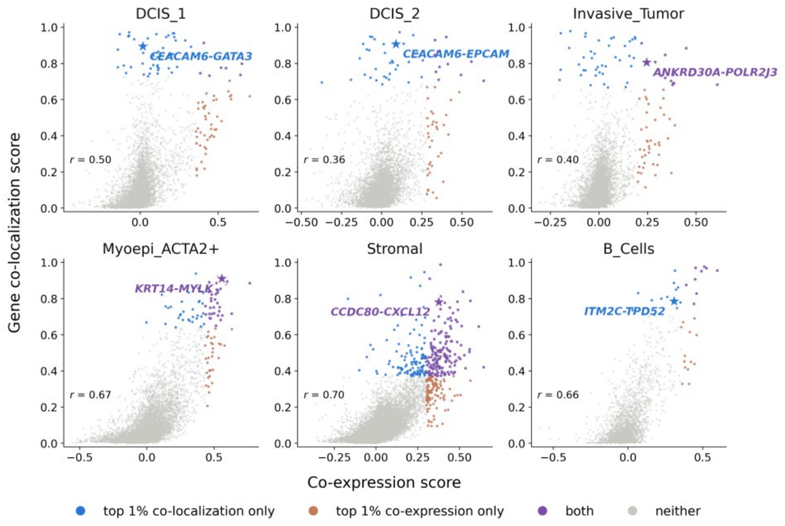

**Supplementary Figure 66. Spatial distribution of ACTA2+ myoepithelial cells relative to DCIS regions in the two sections of the Xenium human breast cancer dataset.**

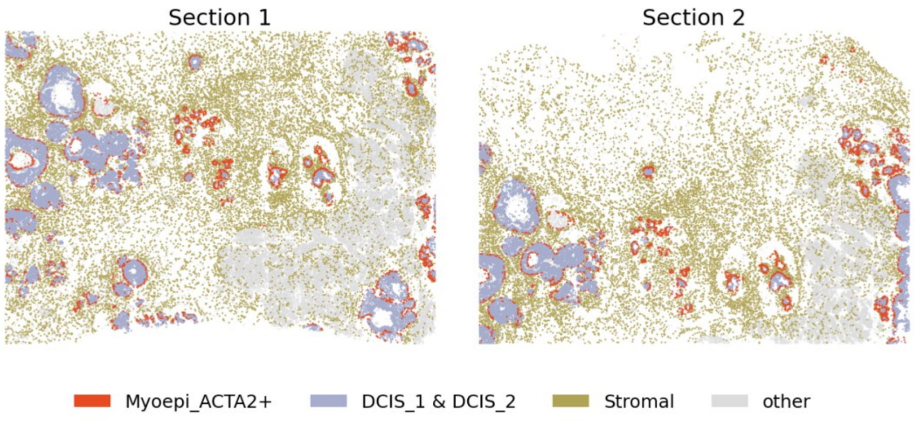

**Supplementary Figure 67. The nuclear-density field and the anatomy-derived basal reference direction in the Xenium human breast cancer dataset.** Left: the smoothed nuclear-density field, built from nucleus centroids on a 3-µm grid with an 18-µm Gaussian kernel; contour lines mark three density levels. Right: magnified view of a representative duct showing the anatomy-derived basal reference directions for ACTA2+ myoepithelial cells. Each arrow represents the normalized negative gradient of the local nuclear-density field and points outward from the densely cellular ductal region toward the surrounding lower-density tissue, consistent with the expected apicobasal orientation of myoepithelial cells.

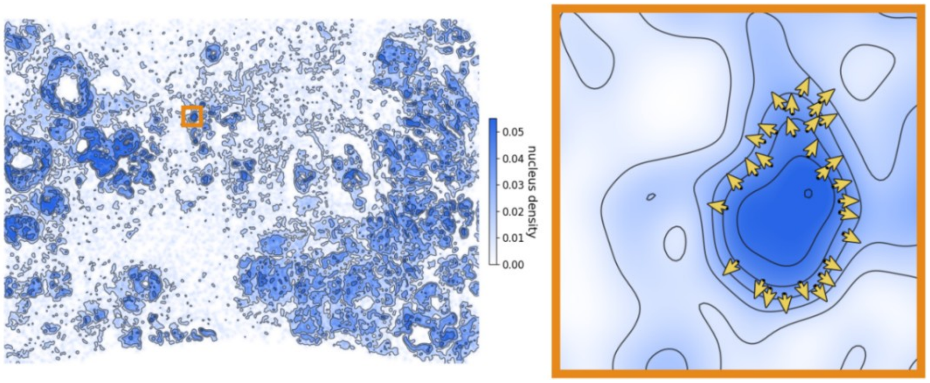

**Supplementary Figure 68. The shared directional component compared with a rotated-direction null in ACTA2+ myoepithelial cells from the Xenium human breast cancer dataset.** The histogram shows the proportion of variance explained by the first shared directional component across 100 rotated-direction nulls. For each control, every cell's anatomical reference direction was independently rotated by a random angle, after which the per-gene regression and singular value decomposition were repeated. The vertical line marks the value obtained with the observed anatomy-derived direction. No null reached the observed value, yielding an empirical *P* value at the floor set by 100 permutations (*P* value = 1/101 < 0.01).

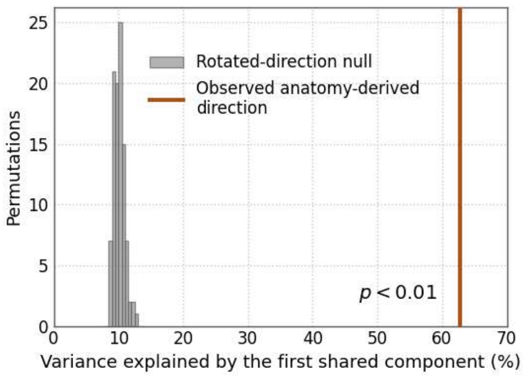

**Supplementary Figure 69. Co-localization change versus co-expression change for the partner genes of *CXCL12* in endothelial cells of the Xenium human breast cancer dataset.** Scatter plot compares, for the 132 partner genes of *CXCL12*, the change in subcellular co-localization between epithelium-adjacent and deep-stromal endothelial cells (x axis) with the corresponding change in cell-level co-expression (y axis), defined as the difference in Pearson correlation between the two genes' size-normalized, log1p-transformed counts. Point color marks the partners with significantly stronger co-localization in epithelium-adjacent (red, 22 genes) or deep-stromal (blue, 29 genes) cells at FDR < 0.05; gray points are the remaining 81 partners. The Pearson correlation between the two changes is annotated in the plot.

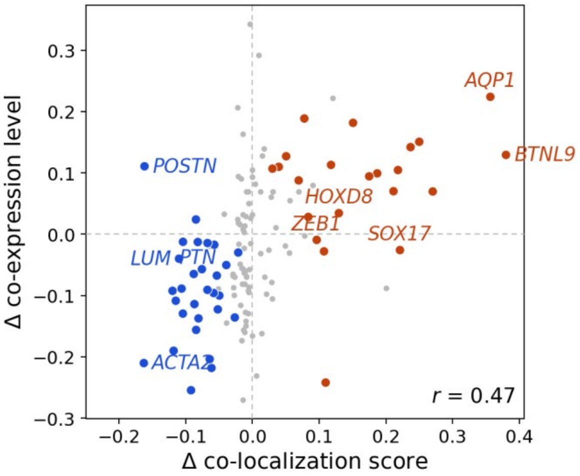

**Supplementary Figure 70. Functional enrichment of the two *CXCL12* partner sets in endothelial cells of the Xenium human breast cancer dataset.** Bar plot shows enriched terms among partners with significantly stronger *CXCL12* co-localization in epithelium-adjacent cells (orange, 22 genes) or deep-stromal cells (blue, 29 genes). Enrichment was computed with Enrichr, querying GO (biological process, cellular component and molecular function), MSigDB Hallmark, Reactome and WikiPathways. *P* values were adjusted separately within each gene-set library using the Benjamini-Hochberg procedure. The x-axis is -log10 of the adjusted *P* value.

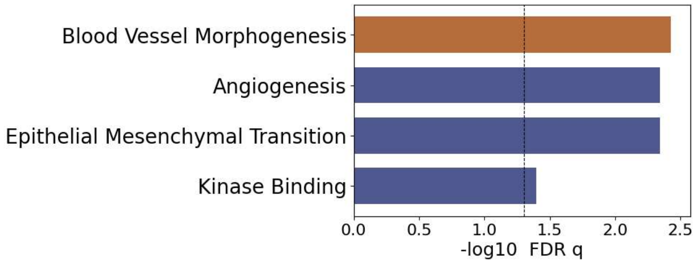

**Supplementary Figure 71. Distribution of the per-cell subcellular change for each downstream gene under *in silico LUM* depletion in stromal cells of the Xenium human breast cancer dataset.** The ridgeline plot shows kernel density estimates of the per-cell change in each analyzed gene's relative transcript distance from the nuclear center. Negative values indicate redistribution toward the nuclear center, whereas positive values indicate redistribution toward the cell periphery. Genes are ordered by their mean predicted localization change, with color indicating the direction and magnitude of the mean change. The dashed line marks no change. Bold gene labels identify the six genes with significant localization changes after Benjamini-Hochberg correction across genes (FDR < 0.05).

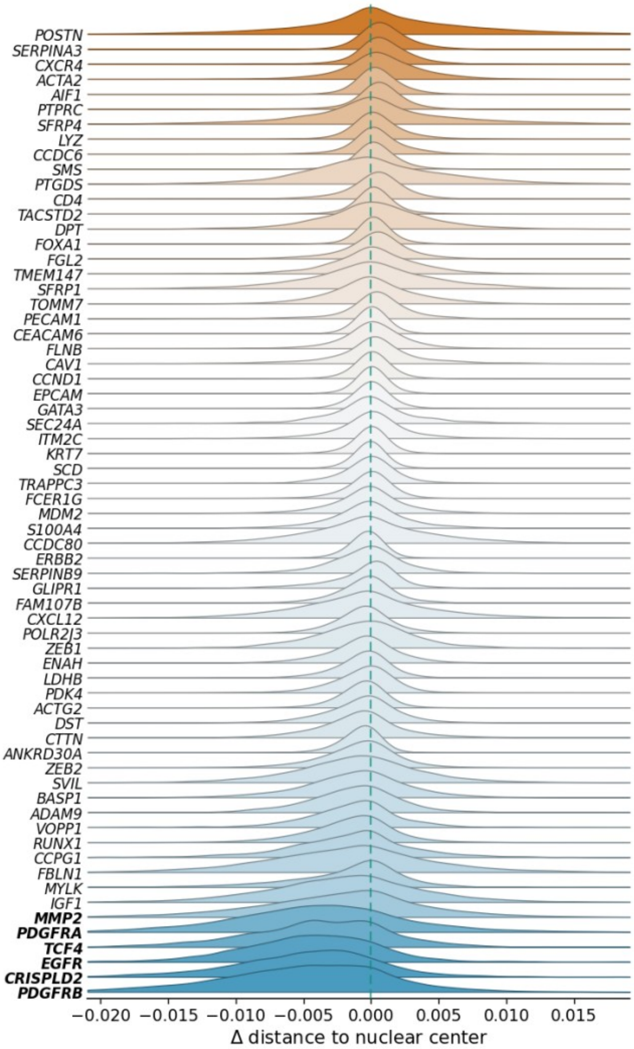

**Supplementary Figure 72. Radial profile of the predicted subcellular change of *POSTN* under *in silico LUM* depletion across spatial contexts in stromal cells of the Xenium human breast cancer dataset.** Bars show the change in predicted mass fraction summed within six concentric rings running from the nuclear center to the cell boundary, computed separately for stromal cells located in epithelium-adjacent, intermediate and deep-stromal regions. Each cell's predicted map was normalized to sum to one before averaging within a class.

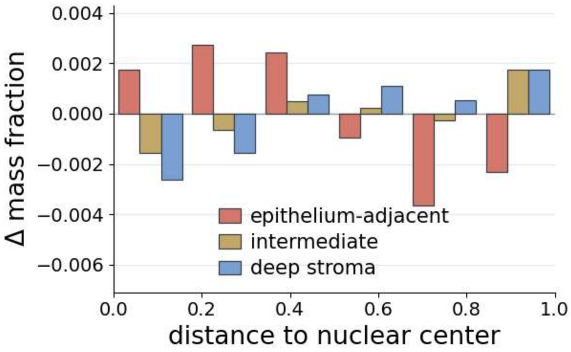

**Supplementary Figure 73. A scatter plot compares the cell-level and the subcellular responses of 60 genes to an *in silico* depletion of *ERBB2* in tumor-related cells of the Xenium human breast cancer dataset.** The x-axis shows the log2 fold change of predicted counts and the y-axis the change in relative distance to the nuclear center (inward, negative; outward, positive); point color and shape indicate which of the two readouts is significant at FDR < 0.05.

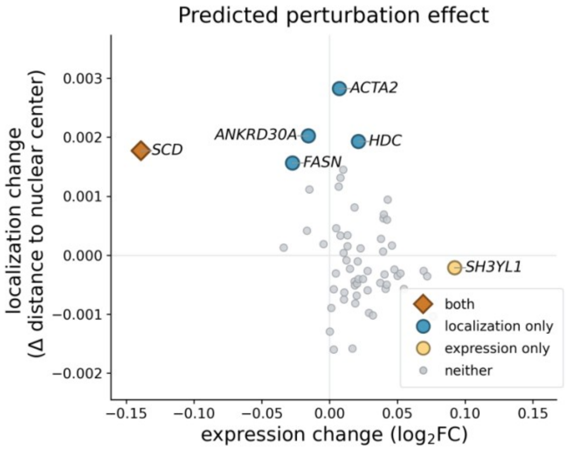

**Supplementary Figure 74. Distribution of the per-cell subcellular change for each downstream gene under *in silico ERBB2* depletion in tumor-related cells of the Xenium human breast cancer dataset.** The ridgeline plot shows kernel density estimates of the per-cell change in each analyzed gene's relative transcript distance from the nuclear center. Negative values indicate redistribution toward the nuclear center, whereas positive values indicate redistribution toward the cell periphery. Genes are ordered by their mean predicted localization change, with color indicating the direction and magnitude of the mean change. The dashed line marks no change. Bold gene labels identify the five genes with significant localization changes after Benjamini-Hochberg correction across genes (FDR < 0.05).

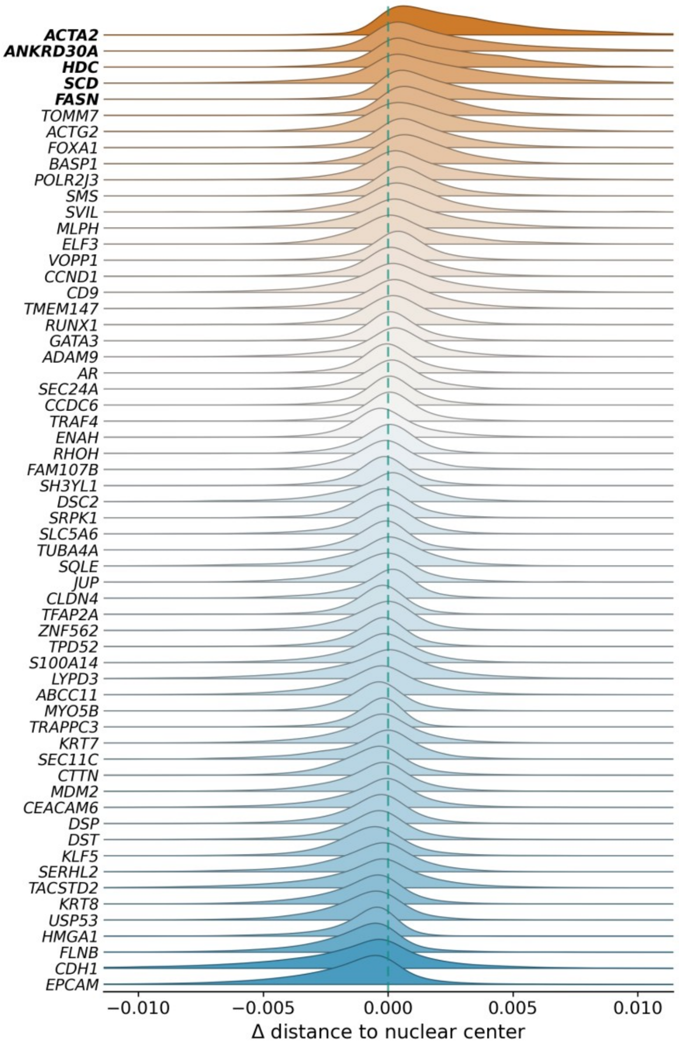

**Supplementary Figure 75. Predicted subcellular localization redistribution of *SCD* following *in silico ERBB2* depletion in tumor-related cells of the Xenium human breast cancer dataset.** Left panel: change in the SVC-predicted subcellular distribution of *SCD* upon *ERBB2* depletion, averaged over cells and shown as the per-pixel change in mass fraction on the $12 \times 12$ image; each cell's map was normalized to sum to one before averaging. Right panel: the same change summed within six concentric rings from the nuclear center to the cell boundary.

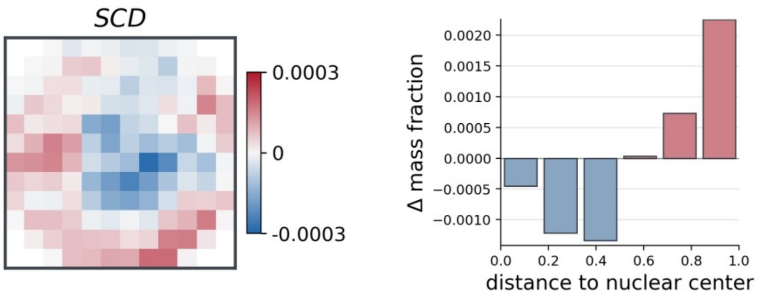

**Supplementary Figure 76. Comparison of the original and Cellpose cell segmentations in the Xenium human breast cancer dataset.** Each row shows a representative 110 × 110 µm field from a 1,275 × 1,275 µm region of section 2. The left column shows the DAPI image; the middle column shows the original 10x Xenium cell segmentation provided with the dataset; and the right column shows an alternative Cellpose segmentation generated from the same DAPI image using the Cellpose nuclei model (diameter = 25 px, flow threshold = 0.9, cell-probability threshold = -2.0), followed by a 10 µm outward expansion of each nuclear mask to define cell boundaries. In both segmentation overlays, cells are filled with random colors and outlined in yellow.

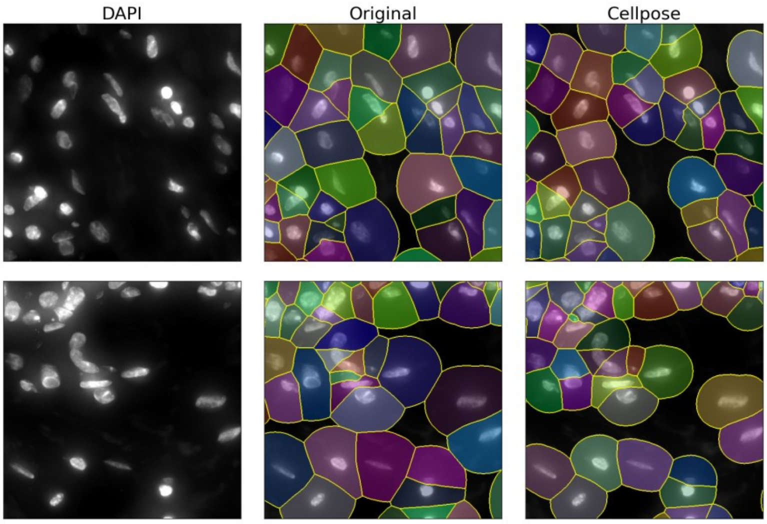

**Supplementary Figure 77. Robustness of SVC prediction and gene co-localization to the choice of cell segmentation. a.** Mean Pearson correlation (PCC) between predicted and measured expression under the original 10x Xenium and Cellpose segmentations, evaluated at the cell, gene, and subcellular levels using a 10-fold masked-gene evaluation in stromal cells from a 1,275 × 1,275 µm region of section 2. Bars show the mean and error bars indicate ±1 standard deviation. **b.** Comparison of pairwise gene co-localization scores between the two segmentations for the 50 most highly expressed stromal genes. Spearman and Pearson correlations are indicated. **c.** Co-localization scores under both segmentations for the ten highest-scoring gene pairs identified using the original segmentation.

**a**

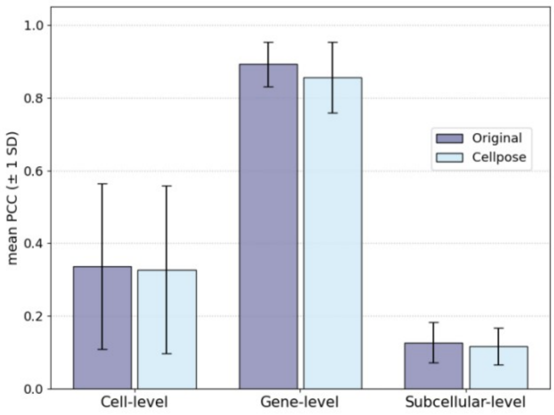

**b**

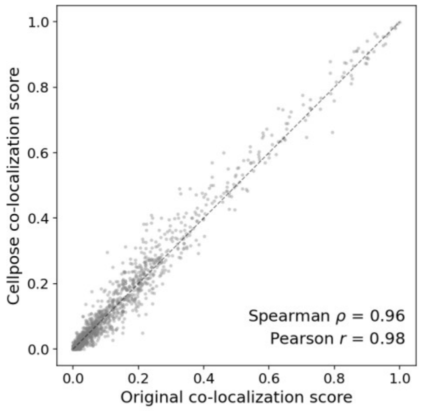

**c**

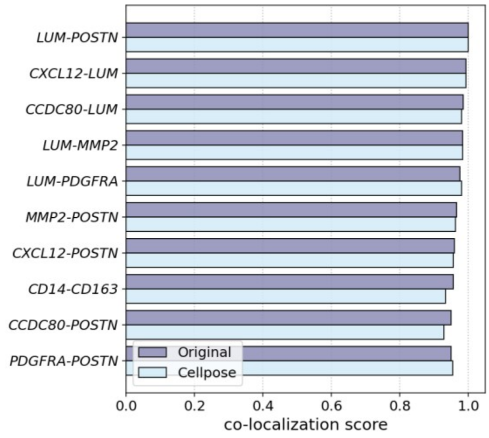

**Supplementary Figure 78. Illustration of the three-zone subcellular partition used in SVC for the MERFISH U2-OS data analysis.** The inner, middle, and outer zones approximate the nuclear, cytoplasmic, and membrane-associated regions, respectively.

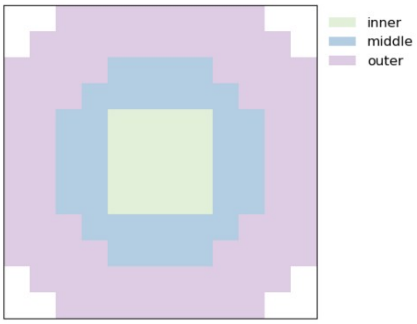

**Supplementary Figure 79. Sensitivity of SVC performance to hyperparameter λ.**

SVC was trained with $\lambda \in \{0, 1, 3, 10, 30, 100\}$ using five random seeds on **a.** seqFISH+ and **b.** MERFISH U2-OS datasets, and evaluated by PCC and RMSE across cells, across genes and within subcellular spatial maps. Lines show the mean across five seeds and error bars indicate the standard deviation; the dashed line marks the default $\lambda = 10$.

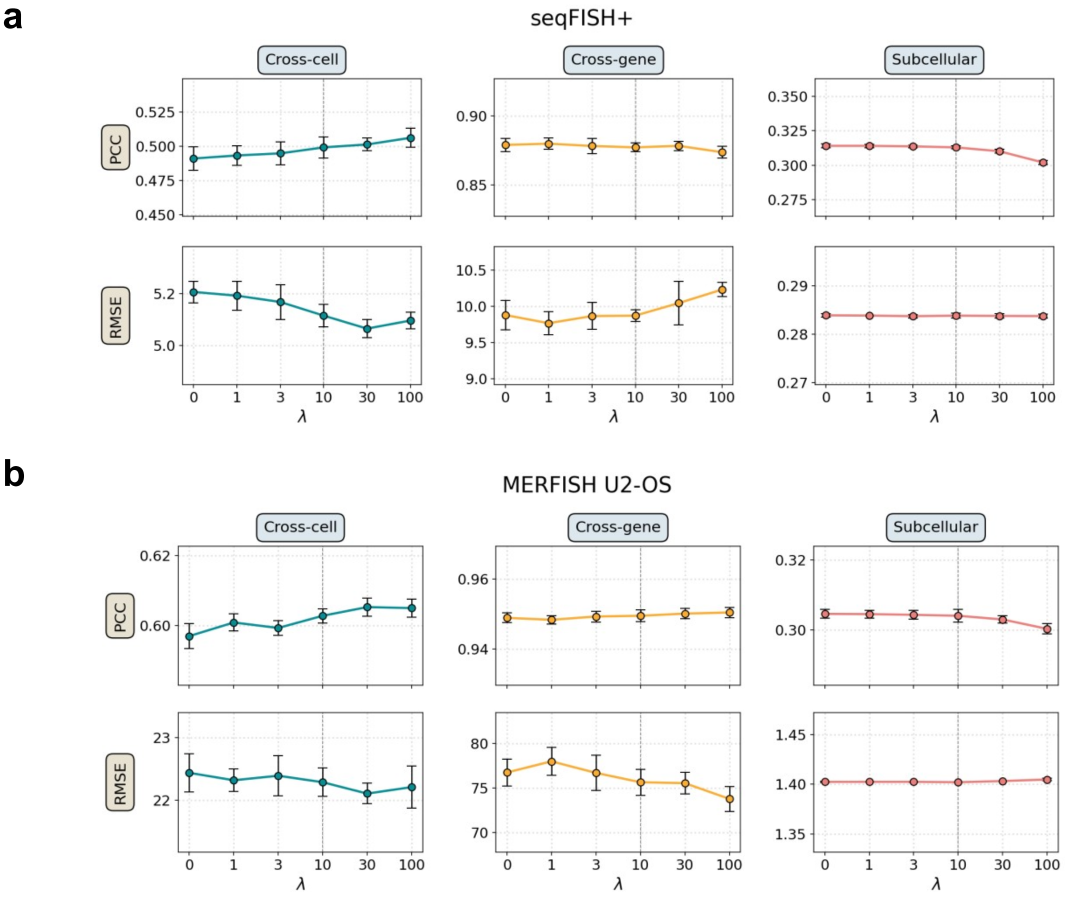

**Supplementary Figure 80. Training and validation loss curves of SVC across four datasets.** Training and validation losses are shown across training epochs for seqFISH+, MERFISH U2-OS, Xenium mouse brain, and Xenium breast cancer. The checkpoint selected based on the lowest validation loss and the end of training are indicated in each panel.

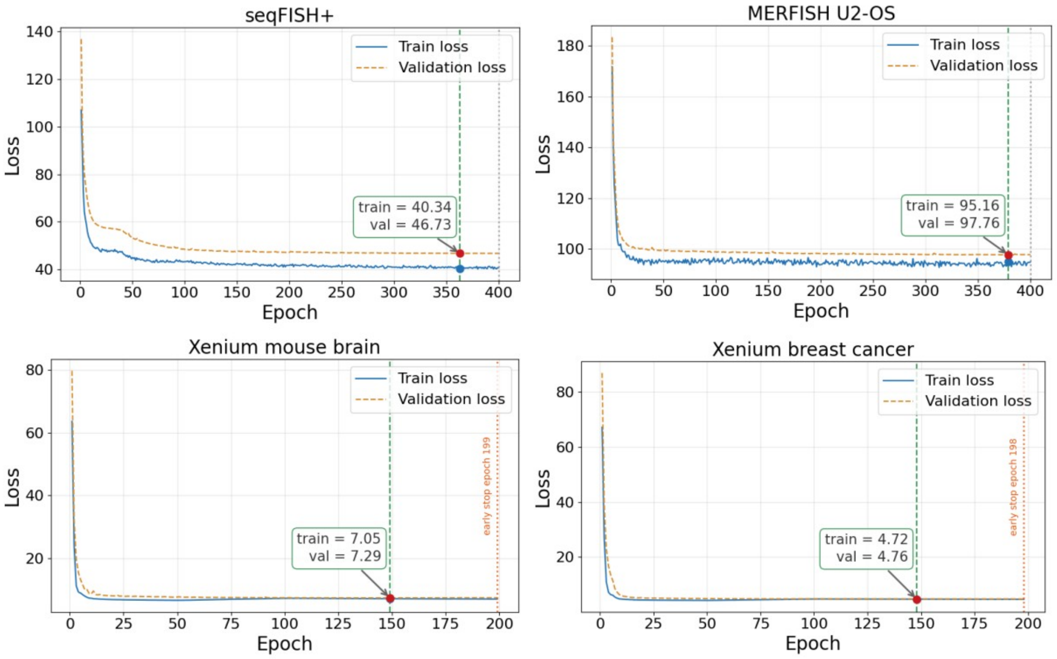

### Supplementary Tables

**Supplementary Table 1. Dataset-specific SVC training configurations and computational resource usage.**

| Dataset | Training/ validation cells | Gene panel | Best/ final epoch | Early stopped | N_gpu | Memory cost (GB) | Time cost (min) |
|---|---|---|---|---|---|---|---|
| seqFISH+ | 145/12 | 1,000 | 363/400 | No | 1 | 39.2 | 11.1 |
| MERFISH U2-OS | 828/72 | 119 | 379/400 | No | 1 | 5.5 | 12.8 |
| Xenium breast cancer | 54,258/6,656 | 294 | 148/198 | Yes | 4 | 17.2 | 176.8 |
| Xenium mouse brain | 81,541/11,906 | 228 | 149/199 | Yes | 4 | 23.8 | 217.0 |

### Supplementary Notes

#### Supplementary Note 1. Ablation study

We performed six different ablation analyses. These include four leave-one-input-out ablations that remove either cell-level morphology embedding, cell-level neighbor embedding, cell-level identity embedding, or gene-level functional embedding, as well as two additional settings to evaluate the contribution of subcellular localization information. These two additional settings include a cell-level-only setting, where the subcellular gene localization embedding was ablated and replaced by only cell-level gene expression counts, so that the model no longer received subcellular spatial distribution information; as well as a subcellular-only setting, where we retained only the subcellular gene localization embedding while removing the four morphology, neighbor, cell identity, and gene-level functional embeddings. These two settings were designed to test thoroughly whether cell-level inputs alone can achieve comparable performance, and whether subcellular localization information itself contributes to SVC performance. We evaluated each of the six ablated models across five random seeds using median PCC and RMSE computed across cells, across genes, and within subcellular spatial maps. For the seqFISH+ dataset, the results of these ablation analyses are shown in Supplementary Fig. 1a. For the MERFISH U2-OS dataset, the results of these ablation analyses are shown in Supplementary Fig. 1b. As cell identity labels are not available for the MERFISH U2-OS dataset, five ablation settings were evaluated.

The ablation results show that the full model achieves the best overall performance across a wide variety of tasks. Notably, the cell-level-only model performs substantially worse on subcellular localization prediction than the full SVC model (seqFISH+: $\Delta$PCC=-0.15, $\Delta$RMSE=+0.0076; MERFISH U2-OS: $\Delta$PCC=-0.082, $\Delta$RMSE=+0.033), underscoring the central role of the subcellular localization embedding in modeling subcellular transcript localization. The cell-level-only model also underperforms in the cross-cell and cross-gene evaluations, indicating that the subcellular localization embedding captures information beyond cell-level inputs alone and improves performance not only for subcellular prediction but also for cross-cell and cross-gene prediction. In contrast, the subcellular-only model achieved competitive performance on subcellular localization prediction, but showed a moderate decline in the cross-cell and cross-gene evaluations, suggesting that the cell-level and gene-level inputs provide complementary information that helps SVC perform more consistently across evaluation settings. More broadly, the ablation results indicate that different modalities contribute in a complementary, dataset-dependent manner, supporting the benefits of joint SVC modeling. Overall, the full SVC model provided the best and most stable performance across evaluation levels, highlighting the benefits of integrating multiple inputs to capture complementary multi-scale information.

#### Supplementary Note 2. Computational scalability of SVC

To rigorously evaluate the scalability of SVC, we first conducted a controlled subsampling experiment using the large Xenium mouse brain dataset. We generated datasets containing 1,000, 5,000, 10,000, 20,000, 40,000, and 80,000 cells, respectively, while keeping the other settings fixed. The validation set was consistently maintained at 10% of each dataset, with the remaining 90% used for training. All experiments were performed on four NVIDIA H100 GPUs using distributed data-parallel training, and each model was trained using the same early-stopping criterion. For each dataset size, we recorded the total training time, runtime per epoch, and peak GPU memory usage per device (Supplementary Fig. 2).

The total training time as well as the average runtime per epoch increased approximately linearly with the number of training cells. Peak GPU memory increased only moderately, from approximately 10 GB for the smallest subsets to 15 GB for the 80,000-cell subset. This increase was primarily attributable to the validation data cached during evaluation, because the validation set was maintained at 10% of the total dataset and therefore grew with dataset size. By contrast, the GPU memory required for each training step remained approximately constant.

We additionally report the end-to-end computational requirements for the four benchmark datasets, including the numbers of training and validation cells, gene-panel size, training epochs, number of GPUs, peak GPU memory usage per device, and total training time (Supplementary Table 1). Because the existing methods included in our gene-imputation benchmarks operate primarily on cell-level expression matrices and do not perform the same subcellular-resolution prediction task as SVC, their computational requirements are not directly comparable. We therefore focus on the absolute computational requirements of SVC and its scaling behavior.

Across these datasets, SVC completed training within approximately 11-217 min. The models for the two large Xenium datasets were trained on four NVIDIA H100 GPUs within approximately 3-4 h. Together with the controlled subsampling experiment, these results show that the runtime of SVC increases approximately linearly with the number of cells, whereas the GPU memory required for model training is determined primarily by the gene-panel size. Because SVC uses Performer-based linear attention, the memory requirement of the attention module scales approximately linearly with the number of gene tokens, rather than quadratically as in standard self-attention. Overall, these results support the scalability and practical applicability of SVC to large-scale subcellular spatial transcriptomic datasets.

#### Supplementary Note 3. Evaluation of transcript sparsity in SVC subcellular localization analyses

To evaluate whether transcript abundance affects subcellular localization inference, we examined whether SVC prediction residuals showed expression-dependent variability or directional bias, assessed prediction performance across the full expression range, and documented how sparse cell-gene observations were handled in downstream analyses.

We first performed an expression-stratified residual analysis on the seqFISH+ test data to directly quantify the variability and directional bias of SVC localization predictions across transcript-abundance levels. Specifically, we divided the 1,000 genes into five quintiles (Q1-Q5; 200 genes per group) based on their average expression levels. For each detected cell-gene pair, the predicted and measured maps were normalized over the foreground pixels, and their residual was projected onto the radial, x, and y coordinates. For each gene, the mean displacement across cells quantified systematic bias, whereas its standard deviation quantified cell-to-cell residual variability. Residual variability decreased by approximately 30% from the lowest to the highest expression quintile, with median values declining from 0.399 to 0.269 pixels radially, from 0.918 to 0.637 pixels along the x axis, and from 1.019 to 0.717 pixels along the y axis (Supplementary Fig. 5a). In contrast, the mean displacement remained close to zero across all quintiles and showed no consistent trend with expression. Across genes, the pooled mean residuals were -0.005, -0.030, and 0.021 pixels in the radial, x, and y directions, respectively, with the largest corresponding to only 0.25% of the cell-image width. Consistent with our expectations, these results show that transcript sparsity increases the stochasticity of localization estimates, but does not introduce detectable directional bias in the expected localization pattern.

We next examined whether expression-level differences impact the reliability of SVC's subcellular localization prediction. Using the same five expression quintiles, we compared SVC with the four baselines across the full expression range. Within each expression group, we quantified subcellular-level prediction accuracy using PCC, cosine similarity, and RMSE (Supplementary Fig. 5b). We also visualized the average subcellular prediction PCC at the per-gene level by plotting each gene's average subcellular PCC against its expression level (Supplementary Fig. 5c). As expected, prediction accuracy depended on gene expression level. From Q1 to Q5, where the median per-cell expression level in Q5 was 3.8-fold higher than that in Q1, both PCC and cosine similarity gradually increased, consistent with high-count genes providing less noisy observed subcellular patterns. RMSE is less informative in this setting because it is scale-dependent: highly expressed genes have larger absolute count ranges and are therefore expected to exhibit larger absolute prediction errors even when relative accuracy is similar. Consistent with this, RMSE increases across expression groups. Importantly, SVC consistently outperformed all four baselines across all five expression groups. The performance gap between SVC and the baselines remained broadly stable across the expression range and did not diminish in the low-expression groups. This trend was further supported by the dot plot of all 1,000 genes, where SVC achieved better subcellular prediction PCC than the most competitive baseline (Supplementary Fig. 5c; Baseline 4, which uses the per-gene population mean in the test data and is therefore not a deployable predictor; we include it only as a stringent reference point) across the full dynamic range of gene expression. Together, these results indicate that SVC's improvement over the baselines is not restricted to highly expressed genes and remains robust for sparse genes.

To carefully handle sparse genes in downstream analyses, we therefore first explicitly quantified transcript coverage across the four primary benchmark datasets used in these analyses and adopted filtering criteria designed to down-weight low-information observations while retaining broad coverage where appropriate. Among the four datasets, the two cell-culture ones have relatively high per-cell coverage: in seqFISH+, the mean cell-gene image contains 17 transcripts, and 88.5% of images contain at least 5 transcripts; in MERFISH U2-OS, the corresponding mean is 113 transcripts, with 95.9% containing at least 5. In contrast, the two Xenium tissue datasets have much lower per-cell coverage, with mean cell-gene counts of 1.7 transcripts in mouse brain and 1.1 transcripts in breast cancer. However, because these tissue studies profile many more cells, they still provide substantial gene-level support: even after requiring at least 5 transcripts for a cell-gene localization estimate, a median of 10,660 cells per gene in mouse brain and 915 cells per gene in breast cancer remain available. Thus, although individual cell-gene observations are sparser in the tissue datasets, there are typically sufficient numbers of cells available to support robust downstream analyses.

In terms of how sparse genes are handled, we used analysis-specific quality control criteria. For the localization prediction benchmark, we retained all genes in the evaluation in order to assess comprehensive model performance across the full expression range, including both sparse and highly expressed genes. For downstream analyses in which the results are interpreted in terms of subcellular localization or co-localization, including mean distance to the nuclear center and gene co-localization analysis, we applied gene-level filtering to reduce the influence of unreliable sparse genes. Specifically, within a cell population or a selected group of interest, such as a specific cell type, we retained only genes with at least 20 valid cells, where a valid cell was defined as having at least 5 transcripts for that gene. For subcellular localization analyses based on the mean distance to the nuclear center, cell-gene images with total subcellular transcript counts below 5 were further excluded. This filtering strategy ensures that the subcellular localization estimates are supported by sufficient transcript observations while still allowing comprehensive model evaluation. Any additional filtering applied in individual analyses is specified in the corresponding Methods sections.

#### Supplementary Note 4. Evaluation of SVC against gene-specific mean-distance baselines

To assess whether SVC captures subcellular localization variation beyond gene-specific population averages, we compared SVC with two gene-specific mean-distance baselines. The first baseline, denoted as "baseline (train)", uses the mean distance to the nucleus for each gene computed from the training cells as the prediction for the test data. The second baseline, denoted as "baseline (test)", uses the mean nuclear distance for each gene computed from other test cells. We note that the proposed "baseline (test)" becomes equivalent to the ground truth test-set mean and therefore serves as an oracle reference rather than a comparable prediction baseline for the particular task of predicting the mean distance. The scatter plots and PCC values for SVC and these two baselines on the seqFISH+ dataset are shown in Fig. 2b and Supplementary Fig. 6a. The results show that the baseline (test) serves as the oracle for the task of predicting the mean distance to the nuclear center and, as expected, achieves perfect agreement. SVC and the baseline (train) also perform extremely well. Compared with the baseline (train), SVC's predictions are less biased with respect to the true distance, whereas the baseline (train) tends to underestimate the distance to the nuclear center despite having a slightly higher PCC (0.98 vs 0.97). Importantly, the results above reflect performance in predicting the mean distance to the nuclear center, even though individual cells can exhibit substantial variation. To assess how well SVC and the two baselines capture cell-to-cell variation in each gene's nuclear distance, we further computed, for each gene, the RMSE between predicted and ground-truth distances to the nuclear center across test cells and compared SVC against both baselines. These results are shown in Supplementary Fig. 6b. The RMSE gain boxplot shows that SVC substantially outperformed both baselines, with positive gains in more than 87% of genes, suggesting that SVC better captures cell-specific variation beyond the gene-level average distance to the nuclear center. This cell-to-cell variation is not captured by either "baseline (train)" or "baseline (test)", explaining the performance gap between SVC and the two mean-distance baselines. Similarly, for the MERFISH U2-OS dataset, Supplementary Figs. 18 and 19 show the scatter plots and PCC values for SVC and the two mean-distance baselines, together with the corresponding RMSE gain boxplot for cell-specific nuclear-distance prediction. The results are largely consistent with those from seqFISH+. Specifically, SVC performs comparably to baseline (train) and baseline (test), the latter serving as an oracle reference, in predicting gene-level mean distances, while substantially outperforming both baselines in predicting cell-specific distances to the nuclear center, with positive gains in more than 91% of genes.

We note that the two datasets above consist of cultured cells: the training and test data come from the same cell line but different FOV batches, and the gene-specific mean distances to the nuclear center are therefore highly similar between training and test cells. To further assess performance in a more heterogeneous setting, we performed an additional analysis on the tissue-level Xenium mouse brain dataset. Using the same training data from section 1, we selected five distinct cell types from the test cells and, for each cell type, compared the predicted and ground-truth mean distances to the nuclear center. The results comparing SVC with “baseline (train)” on the Xenium mouse brain dataset are shown in Supplementary Fig. 35. We did not show “baseline (test)” because, at the population-average level, it serves as an oracle and is identical to the ground truth, as expected. The results show that “baseline (train)” became much less competitive in this heterogeneous tissue setting, even for predicting the mean distances. In contrast, SVC maintained strong agreement with the ground-truth gene-level nuclear-distance patterns across the selected test cell types. Together, these findings suggest that SVC can predict cell-type-specific distances to the nuclear center in the test data, rather than simply recapitulating the average nuclear-distance pattern learned from the training population. This improved generalization is particularly important in tissue contexts, where subcellular spatial organization can differ substantially across cells.

#### Supplementary Note 5. Hyperparameter selection of competing methods

We tuned the compared methods using a validation-based strategy. For each method, we selected key hyperparameters and performed grid searches around the default values, including *k_nearest* and *kl_coeff* for ENVI, *lambda_ot* and *lambda_kl* for uniPort, *model_library_size* and *n_latent* for gimVI, and *lambda_d* and *lambda_g1* for Tangram.

For each dataset, hyperparameters were selected using a validation set held out from the original training data, separate from the final test set. For each candidate hyperparameter setting, we computed the median per-gene RMSE and PCC on the validation set, applied min-max normalization to each metric across candidate settings, and used their average as the validation score for model selection. The dataset-specific hyperparameter selections for each compared method are summarized in Supplementary Fig. 7, where asterisks indicate the selected optimal hyperparameter combinations and triangles indicate the default settings.

#### Supplementary Note 6. Morphology control analyses of cell-cycle-associated subcellular co-localization in the seqFISH+ dataset

Cell and nuclear morphology can vary across cellular states, including different phases of the cell cycle, and such variation may influence the interpretation of subcellular localization patterns. SVC partially accounts for these geometric differences by registering transcript coordinates into a unified nuclear-centered coordinate system. However, because morphology can also covary with biological states such as the cell cycle, we further evaluated whether the cell-cycle-associated subcellular co-localization patterns identified by SVC in the seqFISH+ dataset (Fig. 2l) could be explained by morphology-associated variation.

We first characterized gross morphological variation across cell-cycle phases by measuring cell area and eccentricity for each cell and jointly visualizing these two features (Supplementary Fig. 13). G1, S, and G2/M cells showed substantial overlap in this two-dimensional morphology space, with a silhouette coefficient of -0.07 for the cell-cycle labels, indicating little separation among phases based on these two gross morphological features alone. This substantial morphological overlap is consistent with the morphology-aware analyses below, in which most cell-cycle-associated co-localization patterns remained significant after accounting for cell morphology.

Next, we conducted two complementary morphology-aware control analyses to evaluate whether, and to what extent, the NIH/3T3 cell-cycle co-localization pattern was driven by morphology-associated effects. First, we evaluated an SVC model trained without the cell and nuclear morphology inputs to assess whether the original result was driven by cell and nuclear morphology. Second, we applied PCA directly to the flattened cell and nuclear morphology masks to obtain a low-dimensional morphology representation. We then regressed the SVC latent representations on this morphology representation to obtain morphology-residualized embeddings, which had morphology-associated variation removed, and then repeated the co-localization analysis using these residualized embeddings. The corresponding co-localization boxplots are shown in Supplementary Fig. 14. We found that the cell-cycle-associated co-localization patterns were largely robust to both control analyses, with 11 of the 12 tests remaining significant. For example, nuclear-localized gene pairs still showed significantly higher co-localization in G1 and S phases than in G2/M phase, with 62-73% and 63-66% of gene pairs showing higher co-localization in G1 and S, respectively, across the two control analyses. Similarly, protrusion-related gene pairs showed significantly higher co-localization in G1 phase than in both S and G2/M phases, with 73-84% and 69-84% of gene pairs showing higher co-localization in G1 in the two respective comparisons. As a complementary visualization, we examined the subcellular distribution of the protrusion-related gene *Ddr2* in representative cells with broadly comparable cell area and shape across the three cell-cycle phases (Supplementary Fig. 15). *Ddr2* showed more prominent protrusion-associated localization in G1 cells than in S and G2/M cells, whereas a relatively larger fraction of transcripts appeared within the nuclear region in S and G2/M cells.

The main exception in the quantitative analysis (Supplementary Fig. 14) was the comparison between the G2/M and S phases for cytoplasm-localized genes. In the original analysis, 64% of gene pairs showed higher co-localization in G2/M, and the difference was significant. After morphology residualization, this proportion decreased to 47% and the difference was no longer significant, whereas it remained significant in the morphology-ablated model, with 58% of gene pairs showing higher co-localization in G2/M. This result indicates that the G2/M-associated cytoplasmic co-localization pattern is associated with morphology-related changes, consistent with expected cell-cycle-associated remodeling of cell and nuclear organization. At the same time, the persistence of the other cell-cycle-associated patterns after morphology residualization suggests that SVC-derived subcellular localization embeddings retain localization-related biological signals beyond cell morphology.

Importantly, the role of morphology in subcellular localization is context dependent. Differences in cell and nuclear geometry can introduce technical variation in localization measurements, but morphology can also reflect biologically meaningful cell-state differences that are intrinsically linked to RNA organization. Therefore, removing morphology-associated variation may in some settings also remove genuine biological signal. We thus treat morphology adjustment primarily as a sensitivity analysis and compare results with and without morphology control to distinguish localization patterns that are robust to morphological variation from those that are closely coupled to it.

#### Supplementary Note 7. Robustness of SVC to spatial and morphological noise

We further evaluated the robustness of SVC predictions to spatial jitter in RNA coordinates and noise in morphological images. We quantified the relative change in prediction performance compared with the clean-input condition, using the unperturbed measured maps as ground truth and evaluating subcellular, cross-cell, and cross-gene PCC and RMSE across five random seeds.

First, to evaluate robustness to noisy RNA coordinates, we introduced spatial jitter to the transcript coordinates in test cells. The perturbation was controlled by two factors: the perturbation probability $p$, which specifies the fraction of cell-gene images to be jittered, and the noise level, which scales the standard deviation of the isotropic Gaussian displacement applied to each transcript in the selected images. The displacement scale was normalized separately for each dataset based on transcript density.

We varied one key factor at a time and conducted the following two sets of experiments: experiment 1 fixes $p = 0.5$ and sweeps the noise level over 0, 0.5, 1, 1.5 and 2, quantifying how prediction quality degrades as the magnitude of localization noise grows; experiment 2 fixes noise level = 1 and sweeps $p$ over 0, 0.25, 0.5, 0.75 and 1, quantifying how prediction quality degrades as a larger fraction of images is corrupted. For the seqFISH+ dataset, representative subcellular images under different noise levels, together with the results from Experiments 1 and 2, are shown in Supplementary Fig. 23. The results show that SVC is robust to spatial jitter on the seqFISH+ dataset. As the noise level increased to 2 or the fraction of jittered images increased to 1.0, the overall prediction performance remained largely stable across the evaluated metrics. The main observed degradation was a modest decrease in subcellular PCC, with relative changes of -3.7% and -4.3% under the strongest perturbation settings in Experiments 1 and 2, respectively.

We also performed the same RNA-coordinate jitter experiments on the MERFISH U2-OS dataset. Representative subcellular images under different noise levels, together with the results from Experiments 1 and 2, are shown in Supplementary Fig. 25. On the MERFISH U2-OS dataset, SVC also showed robust performance under RNA-coordinate spatial jitter. As the noise level increased to 2 or the fraction of jittered cell-gene images increased to 1.0, the relative performance changes across the evaluated metrics ranged from -0.3% to -17.6%. We observed that the cross-cell and cross-gene RMSE metrics were more affected by spatial jitter than in the seqFISH+ dataset. This difference may be related to the more limited input information available in the MERFISH U2-OS dataset compared with seqFISH+, including its smaller gene panel (119 genes versus 1,000 genes), which provides less observed-gene information for prediction, and the lack of cell-cycle annotations that could serve as explicit cell identity embeddings. Overall, even under the strongest RNA-coordinate perturbation, the performance degradation remained within 18% across all evaluated metrics, supporting the robustness of SVC to spatial jitter.

Second, to test SVC's robustness to noisy morphology inputs, we applied two sets of perturbation experiments to morphological images of test cells: experiment 1: to evaluate sensitivity to noisy morphology boundaries, we perturbed the cell and nuclear masks by applying a Gaussian-smoothed random field to their signed distance transforms and then re-thresholding the perturbed maps to obtain binary masks. The perturbation strength was controlled by noise level $\sigma_{\mathrm{px}}$, which represents the approximate root-mean-square boundary displacement in mask pixels. We applied perturbations to each test cell with probability 0.5 and evaluated noise level $\sigma_{\mathrm{px}} \in \{0, 1, 2, 3, 4\}$, where $\sigma_{\mathrm{px}} = 0$ corresponds to the clean input; experiment 2: we replaced the cell and nuclear morphology images of a subset of test cells with images sampled from other cells in the training data. The perturbation level was controlled by $p_{\mathrm{swap}}$, the fraction of test cells whose morphology was replaced. We evaluated the fraction of swapped cells $p_{\mathrm{swap}} \in \{0, 0.25, 0.5, 0.75, 1.0\}$, where $p_{\mathrm{swap}}$ =0 corresponds to the clean input and $p_{\mathrm{swap}}$=1.0 replaces the morphology of all test cells. For the seqFISH+ dataset, representative morphology perturbations and the results from these two experiments are shown in Supplementary Fig. 24. SVC showed strong robustness to morphology perturbations. As noise level increased to 4 or the fraction of swapped cells increased to 1.0, the relative performance degradation remained below 3% across all evaluated metrics. For the MERFISH U2-OS dataset, representative morphology perturbations and the results from these two experiments are shown in Supplementary Fig. 26. SVC also remained robust to morphology perturbations. As the noise level increased to 4 or the fraction of swapped cells increased to 1.0, the relative performance degradation remained below 11% across all evaluated metrics. Compared with seqFISH+, the MERFISH U2-OS dataset showed moderately greater sensitivity to morphology perturbations. This may be partly because, for the MERFISH U2-OS dataset, no additional cell-cycle annotations were available as cell identity embeddings, making morphology a relatively more important source of information for capturing variation in cellular phenotypes. Overall, these results support the robustness of SVC to morphology noise across datasets, while also suggesting that morphology provides complementary cell-level context that contributes to prediction performance.

#### Supplementary Note 8. Morphology control analyses for SVC-defined clusters in MERFISH human osteosarcoma cells

In the MERFISH human osteosarcoma dataset, SVC identified cell clusters characterized by distinct cell-cycle-associated subcellular gene co-localization patterns (Fig. 3g). Because cell and nuclear morphology can also vary across the cell cycle, we performed complementary control analyses to determine whether these SVC-defined clusters could be accounted for by morphology alone and whether SVC latent representations retained cluster-associated information after removing the modeled morphology-associated variation.

Specifically, we constructed a quantitative morphology-only representation from the cell and nuclear morphology masks. The $48 \times 48$ cell and nuclear masks were flattened into a 4,608-dimensional vector, and we applied PCA to obtain a low-dimensional embedding that captures the major axes of morphological variation. We then asked whether this morphology embedding could explain the SVC-defined clusters in two ways. First, we projected the SVC cluster labels into the morphology-only PCA space and observed no clear separation among the SVC clusters (silhouette score = -0.081), suggesting that morphology does not explain SVC clusters (Supplementary Fig. 29a). Second, we applied the same spectral clustering procedure to the morphology-only representation to obtain morphology-based clusters and compared them with the original SVC clusters. Morphology-only clustering showed very low agreement with SVC clusters (ARI = 0.052; NMI = 0.058), and in particular did not clearly isolate the abnormal SVC cluster (Supplementary Fig. 29b). Together, these results indicate that the SVC clusters are not driven by cell or nuclear morphology.

Next, we tested whether localization-derived features contain cluster-associated information beyond morphology. To do so, we removed morphological effects by regressing the original SVC latent representations on the morphology-only representation and using the resulting residuals as morphology-residualized SVC latent features. By construction, these residual features capture the component of SVC variation beyond what is explained by cell or nuclear morphology. We then compared morphology-only features and morphology-residualized SVC latent features using (i) supervised visualization and (ii) cross-validated cluster classification. For visualization, we fitted linear discriminant analysis (LDA) using the original SVC cluster labels and projected each feature set onto the first two discriminant axes to examine whether the SVC clusters were separable in that feature space. For quantitative evaluation, we trained the same cross-validated classifier to predict the original SVC cluster labels from each feature set and evaluated performance using balanced accuracy, macro-F1, and macro-AUROC. These results are shown in Supplementary Fig. 30a and b. In the LDA visualization, morphology-only features showed substantial overlap among SVC clusters, whereas morphology-residualized SVC latent features yielded markedly clearer separation of the original SVC clusters.

Consistently, cross-validated classification showed that morphology-only features provided only limited recovery of the SVC clusters (balanced accuracy = 0.459, macro-F1 = 0.412, macro-AUROC = 0.682). In contrast, morphology-residualized SVC latent features achieved higher cluster recovery, with balanced accuracy = 0.620, macro-F1 = 0.565, and macro-AUROC = 0.830. These results indicate that the SVC-defined clusters capture substantial localization-associated variation beyond what can be explained by cell or nuclear morphology alone.

As an additional control, we directly computed cell area and eccentricity as two measures of cell morphology. Similar to the low-dimensional embeddings, these two measurements also overlapped substantially across SVC clusters (silhouette coefficient = -0.01; Supplementary Fig. 30c), indicating that morphology provides limited information for explaining the SVC-defined clusters and is insufficient to provide clear separation of clusters.

#### Supplementary Note 9. Comprehensive benchmarks of SVC on two tissue-level Xenium datasets

To evaluate SVC in heterogeneous tissue settings, we extended the subcellular localization and gene-expression prediction benchmarks to the Xenium mouse brain and human breast cancer datasets. For both Xenium tissue datasets, SVC was trained on section 1 and evaluated on the held-out section 2 using the 10-fold masked-gene prediction strategy described in the Methods section.

For subcellular localization prediction, we evaluated SVC using the same metrics and baseline comparisons as in the cell-culture datasets, including PCC, cosine similarity, and RMSE, with comparisons against the same four baselines. The corresponding results for the Xenium mouse brain dataset are shown in Supplementary Fig. 32. For the Xenium mouse brain dataset, SVC consistently outperformed the four baselines, with positive relative gains across all three metrics. To further assess performance across heterogeneous cell populations, we stratified the analysis by seven major cell types, including excitatory cortical neurons (Exc-Cortical), excitatory hippocampal neurons (Exc-Hippocampal), inhibitory neurons, astrocytes, oligodendrocytes, microglia-PVM, and vascular cells (Supplementary Fig. 33). These major cell types were obtained by merging the 42 annotated subclasses according to neurotransmitter identity and cell lineage, with excitatory neurons further separated into cortical and hippocampal (dentate gyrus, CA fields, and subiculum) groups based on anatomical origin. In each group, the predicted mean distances to the nuclear center were highly concordant with the observed distances, and the predicted gene expression levels closely matched the observed expression levels. These analyses showed that, in this complex tissue-level mouse brain dataset, gene expression and subcellular localization patterns varied across cell types. SVC preserved this cell-type-specific heterogeneity and accurately predicted both gene expression levels and subcellular localization trends. The *in situ* visualizations further supported this observation (Supplementary Fig. 34). For example, SVC recovered an extranuclear cytoplasmic distribution of *Arc* in excitatory neurons and a nuclear-proximal localization pattern of *Opalin* in oligodendrocytes, consistent with the observed localization patterns.

Similarly, the corresponding subcellular localization prediction results for the Xenium human breast cancer dataset are shown in Supplementary Figs. 55-57. For the cell-type-resolved analysis, the 20 cell types annotated in the original study were merged into seven major groups: invasive tumor (invasive and proliferative invasive tumor), DCIS (two DCIS subtypes), stromal, myoepithelial (ACTA2+ and KRT15+), vascular (endothelial and perivascular-like), myeloid (macrophages, dendritic cells, and mast cells), and lymphoid (B cells, CD4+ T cells, and CD8+ T cells). The two hybrid categories and unlabeled cells were excluded. SVC also showed improved subcellular localization prediction performance over the baselines on the Xenium human breast cancer dataset. The visualizations further confirmed that SVC accurately recovered transcript-specific localization patterns across different cell types.

For example, SVC captured the nuclear-proximal localization pattern of *VWF* in vascular cells and the more diffuse cytoplasmic distribution of *ERBB2* in both invasive tumor and DCIS regions, consistent with the observed localization patterns.

We additionally benchmarked cell-level gene-expression imputation in both Xenium tissue datasets using the same evaluation framework as for the seqFISH+ and MERFISH U2-OS datasets, including comparisons with the same competing methods and the same hyperparameter-selection procedure (Supplementary Note 5; Supplementary Fig. 7). SVC achieved competitive or improved performance relative to the competing cell-level imputation methods (Supplementary Figs. 36 and 58). Together, these analyses demonstrate that the predictive performance of SVC extends from relatively homogeneous cell-culture systems to complex tissue environments, while preserving cell-type-specific variation in both gene expression and subcellular transcript organization.

#### Supplementary Note 10. Bidirectional cross-modal retrieval using SVC representations

SVC integrates subcellular gene patterns with cell morphology and spatial context to learn context-aware representations for subcellular expression prediction. We further investigated whether these learned representations preserve sufficient cross-modal information to support direct retrieval between gene-pattern representations and their associated cellular context. To this end, we developed a simple embedding-based extension of SVC that enables bidirectional retrieval between subcellular gene patterns and morphology or spatial-context representations. Specifically, for each retrieval direction, cells were split 70/30 into a fitting set and a held-out set. PCA was fit to the query and target representations in the fitting set to obtain 50-dimensional representations, followed by a linear ridge mapping from the query space to the target space. The fitted PCA transformations and ridge mapping were then applied to the held-out cells, and each mapped query was matched against the target representations of all held-out cells using cosine similarity for nearest-neighbor retrieval. We evaluated the bidirectional retrieval framework on the Xenium mouse brain dataset, focusing on three cell types: dentate gyrus granule cells, endothelial cells, and oligodendrocytes, and on the top 100 highly expressed genes. We first performed a morphology-to-gene-pattern retrieval analysis. For each cell, its morphology embedding served as the query, whereas its latent representation of gene tokens (i.e., the gene-pattern embedding), extracted after ablating morphology and cell identity, served as the retrieval target.

Retrieval performance was summarized by ROC curves under two criteria for defining positive retrievals (Supplementary Fig. 37). First, for exact cell-instance retrieval, the subcellular gene-pattern embedding from the same cell was treated as the positive target. Second, for subcellular gene-pattern similarity retrieval, the five cells with the highest ground-truth gene-pattern cosine similarity were treated as positives. We also visualized representative query cells and their top 5 retrieved cells using both morphology masks and aggregate transcript subcellular images of genes. This analysis showed above-random but moderate retrieval performance, with AUROC = 0.734 for exact cell-instance matching and AUROC = 0.688 for top-5 gene-pattern similarity retrieval. These results suggest that the SVC morphology embedding captures cell-level information associated with subcellular gene-pattern organization. Consistent with the representative retrieval examples, morphology-derived information was more informative for retrieving broad transcript-organization patterns, such as relatively concentrated versus diffuse localization, than for resolving fine-grained gene-specific localization features or subtle polarity patterns. This is consistent with the design of SVC, in which morphology serves as complementary context for encoding continuous variation in cellular phenotypes and provides useful but incomplete information for subcellular gene-pattern prediction. Accurate prediction of missing gene patterns also depends on other inputs, including observed gene patterns and gene-gene relationships learned by the model.

We next performed the reverse retrieval analysis, using gene-pattern embeddings to retrieve two types of cell-level context: cell/nuclear morphology and spatial neighborhood composition. Spatial context was defined by the cell-type composition of the 20 nearest neighboring cells. For each cell, we extracted a subcellular gene-pattern embedding after ablating morphology, spatial-context and cell-identity information, and used this embedding as the query for the two retrieval tasks. Retrieval performance was assessed using ROC curves for two tasks: exact cell-instance retrieval and top-5 target-similarity retrieval (Supplementary Fig. 38). For morphology retrieval, AUROC was 0.744 for exact cell-instance matching and 0.731 for top-5 morphology-similarity retrieval. For spatial-context retrieval, AUROC was 0.895 for exact cell-instance matching and 0.900 for top-5 spatial-context similarity retrieval. The higher retrieval accuracy for spatial neighborhood composition indicates that SVC gene-pattern embeddings more closely preserve the true spatial-context relationships among cells. These results suggest that SVC gene-pattern embeddings encode information associated with both cell morphology and spatial neighborhood context. Together with the morphology-to-gene-pattern retrieval analysis, this supports that SVC learns context-aware relationships across modalities. Overall, this extension provides a practical cross-modal retrieval capability for users when needed, while serving as a complementary analysis rather than the primary modeling objective of SVC.

#### Supplementary Note 11. Control analysis distinguishing subcellular co-localization from cell-level co-expression

To determine whether the gene-pair subcellular spatial relationships recovered by SVC reflected shared subcellular organization rather than cell-level co-expression, we performed a within-cell spatial-shuffle test. For each analyzed cell, we independently permuted the pixel values across the spatial locations that cell occupies within each gene image, preserving the overall cell-level co-expression relationship while disrupting the subcellular spatial arrangement of the transcripts. We repeated this procedure 1,000 times and compared the observed mean Earth Mover's Distance (EMD) across cells with the corresponding permutation null distribution, where lower EMD indicates greater similarity in subcellular transcript distributions.

We first applied this analysis to the co-localized gene pairs identified from the Xenium 5K subcellular imputation analysis. For *Acsbg1-Slc39a12* and *Cldn5-Cobll1*, the observed mean EMDs were substantially lower than those obtained after spatial shuffling (0.41 versus 0.78 and 0.66 versus 0.85, respectively; Supplementary Fig. 52). The corresponding values in the measured Xenium mouse brain data were 2.13 versus 2.67 and 1.97 versus 3.17, respectively. None of the 1,000 permutations produced a mean EMD as low as the observed value for any comparison (empirical $P$ value < 0.001). These results demonstrate that the recovered co-localization relationships preserve the relative intracellular locations of the transcripts and cannot be explained by cell-level co-expression alone.

We next extended the same analysis to the ten cell-type-specific top co-localization pairs identified by SVC (Fig. 4d). Using the measured transcript maps, we examined whether these pairs exhibited subcellular co-localization beyond that expected from cell-level co-expression alone. For all ten pairs, the observed mean EMD was substantially lower than the corresponding spatial-shuffle null, and none of the 1,000 permutations produced a mean EMD as low as the observed value (empirical $P$ value < 0.001; Supplementary Fig. 39).

We further examined the relationship between subcellular co-localization and cell-level co-expression across gene pairs in a cell-type-specific manner. The two measures were only moderately correlated, with Pearson correlation ranging from 0.34 to 0.52 across ten cell types (Supplementary Fig. 40). Among the top 1% most co-localized pairs in each cell type, only 17-55% (median 39%) are also among the top 1% most co-expressed pairs. Several highly co-localized pairs showed only modest cell-level co-expression, including *Cldn5-Pglyrp1* in endothelial cells and *Meis2-Wfs1* in Meis2 neurons. We further applied the same spatial-shuffle and co-localization versus co-expression analyses to the six cell-type-specific gene pairs identified in the Xenium human breast cancer dataset (Fig. 5d), with consistent results (Supplementary Figs. 64 and 65). Together, these analyses across both tissue datasets show that the cell-type-specific gene pairs identified by SVC capture shared subcellular spatial organization that cannot fully be explained by cell-level co-expression alone.

#### Supplementary Note 12. Cross-dataset and cross-tissue transferability of SVC

To evaluate the transferability of SVC beyond the datasets used for model training, we performed two complementary experiments examining generalization across imaging datasets and tissue contexts. In both settings, SVC was trained exclusively on the source dataset and directly applied to the target dataset without retraining or fine-tuning. Prediction performance was evaluated using the same four baseline predictors and the same metrics used in the main benchmarking analyses, including PCC, cosine similarity, RMSE, EMD, KL divergence, and JS divergence.

For cross-dataset transfer, we trained SVC on the Xenium mouse brain dataset and applied it to an independent mouse brain dataset generated using MERFISH. The two datasets shared 130 genes. For evaluation, we retained 24,957 MERFISH cells with more than 50 total transcripts across these shared genes. The median number of transcripts per cell was 77 in the MERFISH dataset, compared with 173 in the Xenium dataset across the shared genes. For quantitative evaluation, we performed 10-fold masked gene prediction on the shared genes, for which measured MERFISH expression provided ground truth for assessing prediction accuracy. In each fold, the corresponding genes were masked in the MERFISH cells and their subcellular transcript localizations were predicted using the SVC model trained only on Xenium data. Prediction performance was evaluated by comparing the predicted and observed transcript images using EMD, KL divergence, and JS divergence, together with PCC, cosine similarity, and RMSE. Because the MERFISH dataset was highly sparse, we restricted the evaluation to cell-gene images containing at least three observed transcripts. The relative gains of SVC over the four baselines across all evaluation metrics in this cross-dataset transfer experiment are shown in Supplementary Fig. 44. For reference, we also show the within-dataset relative gains for Xenium mouse brain, in which SVC was trained on section 1 and evaluated on section 2, over the baselines, enabling direct comparison with the cross-dataset setting. Representative *in situ* subcellular localization predictions for the MERFISH mouse brain dataset are shown in Supplementary Fig. 45.

In this cross-dataset transfer setting, SVC retained positive median gains over all four baselines across all evaluation metrics, demonstrating that its predictive advantage was largely preserved despite the change in platform and transcript-density distribution. The gains over Baselines 1 and 2 remained substantial. Notably, SVC also outperformed the strong Baseline 4, which predicts each target cell-gene image using the mean image of the same gene from other test cells of the same cell type. Across most metrics, SVC's gain over Baseline 4 showed moderate variation between the within-dataset and cross-dataset settings. For KL divergence specifically, the gain over Baseline 4 was larger under cross-dataset transfer (0.211 vs 0.099). Representative *in situ* prediction visualizations further showed qualitative agreement between predicted and measured subcellular transcript distributions of the unseen MERFISH dataset. Together, these results indicate that SVC learns transferable, cell-specific subcellular localization patterns that generalize across imaging platforms.

For cross-tissue transfer, we trained SVC on the Xenium human breast cancer dataset and directly applied it to an independent Xenium human lung cancer dataset. The training and test datasets originated from distinct organs and independent samples, while both represented epithelial solid-tumor microenvironments. The two datasets shared 141 genes. We retained 5,385 lung cancer cells with more than 50 total transcripts across these genes. The target dataset was substantially sparser, with a median of 60 transcripts per cell, compared with 104 in the breast cancer training dataset. In addition, because the cell-type annotations in the target dataset could not be directly aligned with those in the training dataset, we used the clustering labels included in the original output files of the Xenium lung cancer experiment and mapped each cluster to the breast cancer cell type with the highest cosine similarity between their mean expression profiles across the shared genes. We then applied the same 10-fold masked gene prediction and evaluation protocol described above, restricting evaluation to cell-gene pairs with at least three observed transcripts. The relative gains of SVC over the four baselines are shown in Supplementary Fig. 46. For reference, we also show the corresponding within-dataset gains for Xenium breast cancer, trained on section 1 and evaluated on section 2. Representative *in situ* subcellular localization predictions for the lung cancer dataset are shown in Supplementary Fig. 47.

In the cross-tissue setting, SVC achieved positive median relative gains over all four baselines across nearly all evaluation metrics, indicating robust generalization from the breast cancer training tissue to the lung cancer test tissue. The gains were generally largest over the weaker controls. Relative gains over Baseline 1 increased for all metrics except RMSE, indicating a greater performance separation between SVC and this baseline under cross-tissue evaluation. SVC also consistently outperformed the within-cell pixel-shuffling baseline, Baseline 2, indicating that its predictions preserved meaningful subcellular spatial patterns beyond cell-level transcript abundance. Importantly, SVC retained positive gains over the stronger population-average baselines, Baselines 3 and 4, although the changes relative to the within-tissue setting varied by metric. The median KL divergence gain increased from 0.146 to 0.435 over Baseline 4 and from 0.139 to 0.360 over Baseline 3, while the corresponding PCC gain increased from 0.051 to 0.070 for both baselines. At the gene level, SVC achieved lower KL divergence for 99.2% of genes relative to Baseline 3 and for 100% of genes relative to Baseline 4. In contrast, the median RMSE gains over these population-average baselines decreased in the cross-tissue setting, consistent with absolute pixel-level errors being more sensitive to tissue-specific distribution shifts. Together, these experiments show that SVC retains substantial predictive performance when transferred both across platforms and across tissue contexts without retraining, supporting the generalizability of its learned subcellular representations beyond the source datasets.

#### Supplementary Note 13. Out-of-distribution (OOD) generalization across anatomically distinct tissue regions

To further evaluate the generalizability of SVC beyond anatomically matched tissue regions, we examined its performance under a more stringent out-of-distribution setting in the Xenium human breast cancer dataset. Specifically, we assessed whether representations learned from one anatomical region could be transferred to a spatially distinct and completely held-out region of the same tissue section. To this end, we divided section 1 into upper and lower tissue regions and performed reciprocal train-test experiments, training SVC on one region and evaluating it on the other.

In the first experiment, SVC was trained only on the upper region and evaluated on the held-out lower region; in the second experiment, SVC was trained only on the lower region and evaluated on the held-out upper region. In each case, the held-out region was completely unseen during training. We then performed spatial domain detection across the full section to examine whether SVC-derived representations could generalize to tissue regions not observed during training. The domain detection results from both experiments are shown in Supplementary Fig. 59. The upper and lower regions showed distinct anatomical structures, with the upper region enriched for DCIS-associated and stromal areas and the lower region containing a large invasive carcinoma region. Despite the distinct structures between the two regions, SVC recovered spatially coherent domains in the held-out region in both train-test splits. The inferred domains were also concordant with both the H&E-based tissue annotation shown in Fig. 5a and the spatial domains obtained using the full section shown in Fig. 5b. These results indicate that SVC-derived representations can generalize to anatomically non-overlapping tissue regions that were not observed during training.

We further evaluated subcellular localization prediction performance in both train-test splits, as shown in Supplementary Fig. 60. As expected, restricting training to only half of the tissue section led to a modest decrease in absolute performance of all methods compared with models trained on the full section. Nevertheless, relative performance across methods remained similar, and SVC consistently outperformed all four baselines across the held-out regions, providing additional quantitative support for its stable performance under the OOD setting.

#### Supplementary Note 14. Generalization to cell-type-specific out-of-distribution subcellular localization patterns

Subcellular transcript localization can vary substantially across cell types, providing a stringent setting for evaluating whether SVC can generalize beyond localization patterns represented in the training data. We therefore examined whether SVC could predict cell-type-specific localization patterns that fall outside the range observed across the training cell types. To evaluate this capability from complementary aspects of subcellular organization, we designed two OOD settings based on nuclear proximity and transcript polarity.

In the first OOD setting, we examined whether SVC could predict cell-type-specific nuclear-proximity shifts of the same transcripts. We held out B cells as the target cell type and retrained SVC after removing all B cells from the training and validation sets. We selected genes whose B-cell mean distance to the nuclear center fell outside the range observed across all other expressing cell types and ranked them by their deviation from the mean across training cells. Only cell types containing at least 100 cells were considered. A gene was considered expressed in a cell type if it was detected in at least half of its cells and at least 20 cells contained at least 5 transcripts of that gene. The three genes with the most pronounced perinuclear shifts and the three with the most pronounced peripheral shifts were used for OOD evaluation. We then compared SVC predictions in held-out B cells with the four baselines. Because the held-out cell type is absent from training, Baselines 1 and 3 were constructed from training cells of all remaining cell types. Among these, Baseline 3, the per-gene training mean, represents memorization of localization patterns observed during training. These results are shown in Supplementary Fig. 61. SVC predictions more closely followed the identity line between predicted and observed B-cell distances to the nuclear center. In contrast, Baseline 3, the per-gene training mean baseline, deviated from the B-cell ground truth and instead reflected the localization patterns observed in the training cell types. Consistently, pixel-level evaluation showed that SVC outperformed the four baselines across all three metrics, with particularly pronounced improvements in PCC and cosine similarity. These results indicate that SVC maintained a consistent performance advantage in this held-out-cell-type OOD setting, rather than simply reproducing localization patterns observed during training.

In the second OOD setting, we evaluated a distinct aspect of subcellular localization: transcript polarity. Unlike the mean distance to the nuclear center, which measures the overall radial position of transcripts, polarity quantifies directional asymmetry by measuring the displacement of the transcript centroid from the nuclear center. Higher polarity therefore indicates that transcripts are preferentially concentrated toward one side of the cell. Such asymmetric RNA localization is biologically important because it is associated with functional and structural asymmetries within cells[2]. Because polarity is sensitive to transcript count, the polarity scores were computed after count-matched sampling using the observed transcript count of each cell-gene pair.

We held out Myoepi_ACTA2+ myoepithelial cells from the training and validation sets and evaluated SVC on genes showing OOD polarity in this held-out cell type. Using the same selection criterion as in the first setting, we ranked genes whose Myoepi_ACTA2+ polarity fell outside the range observed across all other expressing cell types by their deviation from the mean across training cells. *LUM* and *POSTN*, two extracellular-matrix-related genes, were the top two candidates, with a clear separation from the remaining genes, and were selected for OOD evaluation. The strong polarity of these two genes is consistent with the apicobasal orientation of myoepithelial cells at the epithelial-stromal interface. This design provides a more challenging OOD test of whether SVC can predict a cell-type-specific polarized localization pattern that was absent from the training data. The quantitative performance and representative localization predictions are shown in Supplementary Figs. 62 and 63, respectively. In this held-out Myoepi_ACTA2+ setting, SVC recovered the polarity shift that was absent from the training data. For both *LUM* and *POSTN*, Baseline 3 predicted a largely symmetric localization pattern, whereas the true Myoepi_ACTA2+ localization was strongly polarized. SVC predictions moved substantially toward the true polarized localization, indicating that SVC captured the cell-type-specific polarity pattern. Consistently, pixel-level evaluation showed that SVC outperformed all four baselines across PCC, cosine similarity, and RMSE. Together, these two OOD experiments show that SVC consistently outperformed the baselines across distinct subcellular localization settings, including nuclear-proximity and polarity shifts. These results provide evidence that SVC captures generalizable subcellular transcript-localization relationships, rather than simply memorizing localization patterns observed during training.

#### Supplementary Note 15. *In silico* perturbation analysis of *ERBB2* depletion in the Xenium human breast cancer data

We also conducted an *in silico* depletion of the *ERBB2* gene within tumor-related cells (DCIS #1, DCIS #2, invasive tumor and proliferative invasive tumor cells) across the tissue and quantified downstream responses of the 60 sufficiently expressed genes (Methods). Joint visualization of the predicted cell-level and subcellular effects again revealed significant but only partially overlapping gene responses, with *SCD* the only gene significant on both readouts (Supplementary Fig. 73). Five genes showed significant subcellular localization changes, all characterized by shifts away from the nuclear center (Supplementary Fig. 74). As a representative example, the aggregated spatial map and radial profiles of *SCD* showed reduced localization near the nuclear center and increased localization at the cell periphery following *in silico ERBB2* depletion (Supplementary Fig. 75). Previous work has shown that *ERBB2* upregulates *SCD* in breast cancer cells[3], consistent with the predicted reduction in *SCD* abundance following *ERBB2* depletion. The predicted relative outward redistribution of *SCD* may reflect lipid-sensitive regulation of its transcript stability during ERBB2-dependent lipid-metabolic remodeling[4].

#### Supplementary Note 16. Robustness of SVC to cell segmentation

Segmentation quality is an important consideration for spatial transcriptomics analyses and computational methods. To assess the sensitivity of SVC to cell segmentation, we conducted an additional robustness experiment. We selected an approximately $1{,}275 \times 1{,}275\ \mu m$ region of interest from Section 2 of the Xenium breast cancer dataset and focused on stromal cells within this region. We then compared the original 10x Genomics segmentation used throughout the SVC analyses with an independently generated alternative segmentation. Specifically, nuclei were segmented from the DAPI image using the Cellpose 2.0 nuclei model (*diameter* = 25, *flow_threshold* = 0.9, and *cellprob_threshold* = -2.0), and each nuclear mask was expanded outward by 10 μm to define the corresponding cell boundary. All transcripts in the original dataset were then reassigned according to these alternative cell boundaries. Compared with the original 10x segmentation, the Cellpose-based segmentation yielded smaller stromal cells (median area, 335 versus 519 $\mu m^2$), identified more cells within the same region (1,493 versus 1,272), and resulted in an approximately 26% lower median transcript count per cell (207 versus 280). Representative regions showing the original 10x segmentation and the newly generated Cellpose-based segmentation overlaid on the DAPI images are presented in Supplementary Fig. 76.

We then used the same SVC model trained on section 1 using the original segmentation and evaluated its consistency across the two test segmentations for prediction performance and gene co-localization analysis. Prediction PCCs at the cell, gene, and subcellular levels under the two test segmentations are shown in Supplementary Fig. 77a. For gene co-localization analysis, we calculated pairwise co-localization scores among the 50 most highly expressed genes in stromal cells. The scatter plot compares the scores obtained under the two segmentation strategies for all gene pairs (Supplementary Fig. 77b), while Supplementary Fig. 77c shows the co-localization scores under both segmentations for the top 10 gene pairs identified using the original segmentation. Across the subcellular, cell, and gene levels, prediction PCCs under the Cellpose-based segmentation were slightly lower than those under the original segmentation, with mean differences of only approximately 0.01-0.04, indicating that the predictive performance of SVC remained largely stable across the two segmentation strategies. In addition, gene co-localization analyses of stromal cells were highly consistent between the original and Cellpose-based segmentations. Pairwise co-localization scores showed strong agreement across the two segmentations (Spearman correlation = 0.96; Pearson correlation = 0.98), and *LUM-POSTN* was identified as the strongest co-localized gene pair in both. These results suggest that the co-localization structure recovered by SVC was largely preserved under the alternative segmentation. One possible explanation for this relative stability is that SVC is designed to model relative subcellular localization relationships among genes, which support both the prediction of missing genes from observed genes and the inference of gene co-localization from latent representations.

Such relational information may be less sensitive to moderate segmentation differences. More broadly, this relative stability is consistent with prior observations from ELLA showing that multi-cell subcellular modeling can mitigate the impact of segmentation inaccuracies on localization analysis[5].
